# spicyR enables robust, well-powered co-localisation tests in large spatial omics cohorts

**DOI:** 10.64898/2026.11.17.767438

**Authors:** Sadiq Dohadwalla, Shreya Rao, Farhan Ameen, Elijah Willie, John T. Ormerod, Ellis Patrick

**Affiliations:** Sydney Precision Data Science Centre, The University of Sydney; School of Mathematics and Statistics, The University of Sydney; Centre for Cancer Research, The Westmead Institute for Medical Research, The University of Sydney; Charles Perkins Centre, The University of Sydney, Sydney

## Abstract

Spatial omics technologies can now profile finely resolved cell populations in patient tissue at scale, offering a way to link their arrangement to outcomes. Yet images often differ in size, shape and cell content, and existing tests between patient groups mistake these differences for co-localisation or weight every image equally. We present spicyR v2, which avoids both problems with a fast, permutation-free test that accounts for these differences and weights each image by how precisely it measures co-localisation. In simulations and 74 analyses of 42 public datasets, it controlled false discoveries, had more power than spicyR v1 and was robust to tears, holes and differences in cell number that misled other methods. These analyses also revealed that many studies relating co-localisation to clinical outcomes are underpowered, and spicyR v2 can estimate the number of patients a new study needs. spicyR v2 analyses 100 million cells in minutes, making robust, well-powered tests of co-localisation practical for large spatial omics cohorts.

## Introduction

Spatial omics technologies now measure dozens of proteins or thousands of genes in every cell of intact tissue [1–4], creating an opportunity to find spatial relationships between cell populations that inform patient outcomes. With only a handful of markers, multiplex immunofluorescence showed that melanoma patients who responded to anti-PD-1 therapy had more CD8 T cells within 20 *µ*m of a tumour cell [5]. Yet, although spatial omics profiles many more cell populations at once, relationships this interpretable are not routinely reported. Instead, many published findings have come from unsupervised clustering of cells into neighbourhoods [6–8], not from tests of specific pairs of cell populations. Such tests are hard to power. Studies often have only tens of patients but hundreds of pairs to test, and changes in co-localisation can be small relative to the variation between patients and the imprecision of images with few cells of a type. Hypothesis-driven research therefore needs tests that control false discoveries yet are powerful enough to detect these changes in cohorts of realistic size.

A comparison of co-localisation between patients starts with a score for each image. The simplest score counts how many cells of one type lie near each cell of another, for example how many CD8 T cells lie within 20 *µ*m of each tumour cell. This count cannot be compared between images on its own, because its expected value doubles when the neighbouring cells are twice as dense, even if the two populations have no affinity for each other. Methods therefore compare the count with the count expected by chance in the same image, and call two populations co-localised when their cells lie closer together than chance would predict. They differ in what they take chance to mean [9] and in how they report the difference from it, and these choices decide whether a score from a small biopsy can be compared with one from a large resection. Some take chance to mean that the cells of each population are scattered at random, independently of each other, with the same density everywhere in the image, so that the expected count depends on the area of the image, or of an outline drawn around the tissue. Ripley’s *K* function and its relatives report the observed count relative to this expectation [10–15]. Others keep every cell where it is and shuffle which cells carry each label many times, mimicking a null hypothesis known as random labelling [9, 16]. These methods most often report how far the observed count lies from the shuffled counts as a *z*-score or a permutation *p*-value [17–22]. For scores to be compared between patients, the same co-localisation must receive the same score whatever the size and shape of the tissue imaged, how densely its cells are packed and how they are divided between populations. Otherwise, a difference between patients’ tissues could be reported as a difference in co-localisation, giving a false discovery, or could hide a real difference.

Neither way of comparing with chance fully meets this requirement. Chance defined by area treats the whole image, or the outline drawn around the tissue, as if cells could lie anywhere in it [23]. When they cannot, as with a hole, a wide tear or a partly empty field of view, the empty space lowers the expected count more than the observed one, so the same arrangement of cells receives a higher score and can be reported as co-localisation. Comparing counts directly with random labelling avoids this problem, because the shuffled labels fall on the same cells as the real ones. However, a *z*-score measures how surprising an arrangement is, not how strongly two populations co-localise. Just as a *p*-value shrinks in a larger study of the same effect, a *z*-score grows with the number of cells imaged for the same co-localisation. Dividing it by the square root of the number of cells corrects for this [22], but it is unclear whether this division also corrects for how densely cells are packed and how they are divided between populations, which also set how much the count varies under random labelling. These properties differ between groups of patients, for example between cancer subtypes with more or less stroma [6, 7], and within groups, where a population can vary many-fold in abundance between patients [24]. Because real tissue has no known truth, methods have mainly been evaluated on simulated tissue [14, 22], so their rate of false discoveries in real cohorts is rarely measured.

Images also differ in how precisely they measure co-localisation, and averaging their scores with equal weights to compare groups of patients lets imprecise images add noise and costs power. An image with five cells of a population says far less about where those cells sit than one with five hundred. Such imprecise images become more common as populations are split into finer and rarer subsets. Weighting each image by its precision, as a meta-analysis weights its studies, reduces this loss. Most methods that weight images derive the weight from the numbers of cells [14, 25–27], as does our earlier method, spicyR v1 [12]. Such weights ignore how densely the cells are packed and how they are arranged, which also set how much an image’s co-localisation score varies under random labelling. PANORAMIC measures each image’s own precision by resampling its cells [15], but this must be repeated for every pair of populations in every image and becomes slow in large cohorts. Spatial statistics has long compared groups of subjects from their images [28–32], weighting images differently [33, 34], and has developed exact formulas for how neighbour counts vary under random labelling, which give this variance without resampling [35–41]. Leveraging these developments will matter more as cohorts grow, from the tens of patients and selected regions studied now to whole slides and consortia that combine images across laboratories [42, 43], whose images will differ more in size, shape and cell content and hold far more cells. Despite the many methods for analysing co-localisation (Supplementary Table S2), there remains a lack of tests that give every image a score that means the same thing whatever tissue was imaged, weight each score by the image’s own precision, and run fast enough for every pair of populations in large cohorts.

To fill this gap, we developed spicyR v2, a test of co-localisation between groups of patients. It scores each image against random labelling of the cells actually imaged, as a share of cells rather than a *z*-score, so that neither empty space nor image size changes what its score means. The exact variance of this score under random labelling follows from classical results for sampling without replacement [35, 44] and can be computed for every pair of populations in a single pass over the cells. We then derived a closed-form test between groups of patients that weights each image by this variance, so spicyR v2 needs no shuffling at any step and analyses 100 million cells in minutes. We demonstrate that spicyR v2 has more power than spicyR v1 while controlling false discoveries in real cohorts, and that it is robust to tears, holes and differences in cell number that other methods mistake for co-localisation.

## Methods

### Overview of spicyR v2

spicyR v2 tests whether the co-localisation of two cell types differs between groups of patients, or is associated with an outcome such as survival. As input, spicyR v2 takes segmented cells with their coordinates and cell types from one or more images per patient, and a group or outcome for each patient. We write patient for the independent unit of a study throughout, but the unit can equally be a donor or an animal, such as a mouse. For an ordered pair of cell types *A → B*, each image is scored by the extra fraction which is the share of *B* cells with an *A* cell nearby, in excess of, or short of, that expected under random labelling. Because the expected fraction comes from a random labelling of the cells actually imaged, in which every cell stays in place and the *A* cells keep their labels, tears, holes and differences in the size and cell content of images are not mistaken for co-localisation. The exact mean and variance of the extra fraction under this random labelling have closed forms that follow from sampling without replacement (Supplementary Section 3), so each image’s precision is known without permutations. A two-level model then combines images within patients and patients within groups, weighting each by its precision, as a meta-analysis weights its studies. The weights also allow for genuine variation between images and between patients, so that no single large image or patient dominates. To compare groups, we derived a closed-form sandwich estimator of the variance of the difference, which uses how much the patients’ effects actually spread rather than the weights. Together, these make the test fast and keep it valid when the precisions behind the weights are wrong. The same model also takes covariates, more than two groups, time-to-event outcomes and several radii at once.

Formally, spicyR v2 models the effect *Y*_*ij*_ of a pair in image *j* of patient *i* as

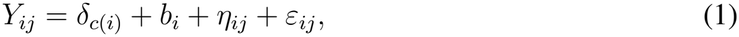

where *c*(*i*) is the group of patient *i, δ*_*c*_ is the mean effect of group *c, b*_*i*_ is the departure of patient *i* from its group’s mean, with variance *τ*^2^, *η*_*ij*_ is the departure of image *j* from its patient, with variance *ω*^2^, and *ε*_*ij*_ is the random-labelling noise of the image, whose variance 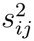 is calculated from the image, as described below.

The group means are estimated by fitting the two-level model (1) as a generalised estimating equation (GEE) [45], which here has a closed-form solution. The groups are compared with a cluster-robust variance with patients as the clusters, which stays valid, for example, when cells of a type aggregate.

### Neighbours and ordered pairs

For a pair of cell types, spicyR v2 asks whether cells of one type have cells of the other type among their neighbours more often than chance would give. Two cells are neighbours if they are within a radius *r* of each other; alternatively, a cell’s neighbours are its *k* nearest cells. The choice of radius determines which relationship between two cell types is tested. A radius of one or two cell diameters asks whether cells of the two types touch, as contact-dependent interactions require, whereas a larger radius asks whether they share a neighbourhood, such as the reach of a secreted signal or a tissue compartment. The radius, or *k*, is therefore best chosen for the relationship of interest before the analysis, and testing that single radius gives the most direct and powerful test. A pair of cell types is ordered and written *A → B*, where *A* is the anchor type, whose cells are held fixed, and *B* is the tested type, whose cells are checked for an *A* neighbour. *A → B* and *B → A* are different questions and are tested separately, and a cell type is also tested with itself (Supplementary Section 2).

### Co-localisation in one image

In each image, spicyR v2 compares the observed number *O* of *B* cells with an *A* neighbour with the number expected under random labelling, in which every cell stays in place, the *A* cells keep their labels, and the *B* labels are shuffled among the cells that are not *A*, which we call the candidates [9, 16]. Because the expectation comes from the cells actually imaged, empty regions such as tears and holes, and differences in the size and cell content of images, are not mistaken for co-localisation. Holding the *A* cells fixed makes *O* hypergeometric, so its mean *E* and variance *V* under random labelling have closed forms and no permutations are needed (Supplementary Section 3.2). Within one image we drop the subscript (*i, j*). If an image has *n*^*B*^ *B* cells and a share *q* of its candidates have an *A* neighbour, then *E* = *n*^*B*^*q*, so *n*^*B*^*q* of the *B* cells are expected to have an *A* neighbour and the other *n*^*B*^(1 *− q*) to lack one. The extra fraction divides the excess *O − E* by a room *R*, the number of *B* cells that could change, so that every image is on a common scale:

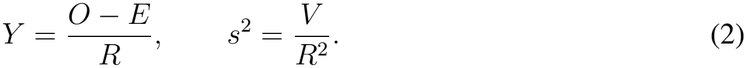

The room depends on the pair’s scale. There are two available: a gain scale or a loss scale. A pair is on the gain scale if, summed over all its images, across groups, more *B* cells have an *A* neighbour than expected (Σ (*O − E*) *≥* 0), so that *B* cells have, overall, gained *A* neighbours, and on the loss scale if fewer do, so that they have lost them. This scale is chosen once per pair and radius, from all images of all patients regardless of which group the patient belongs in. Therefore every image of the pair is on one scale, and that scale does not depend on the comparison (Supplementary Section 3.3). The mean and variance of each image’s *Y* are computed with this scale held fixed.

On the gain scale, the room is *R* = *n*^*B*^(1 *− q*), that is the number of *B* cells expected to lack an *A* neighbour, since only these could gain one. For example, if an image has 100 *B* cells, of which 30 are expected to have an *A* neighbour (*E* = 30), then *R* = 70; if 51 are observed with a neighbour (*O* = 51), the excess is 21 and *Y* = 21*/*70 = 0.3, so 30% of the *B* cells that would lack an *A* neighbour by chance have at least one. *Y* = 1 means that every *B* cell has an *A* neighbour.

On the loss scale, the room is *R* = *n*^*B*^*q*, that is the number of *B* cells expected to have an *A* neighbour, since only these could lose one. For example, if an image has 200 *B* cells, of which 80 are expected to have an *A* neighbour (*E* = 80), then *R* = 80; if 60 are observed with one (*O* = 60), the excess is *−*20 and *Y* = *−*20*/*80 = *−*0.25, so a quarter of the *B* cells that would have an *A* neighbour by chance do not. *Y* = *−*1 means that no *B* cell has an *A* neighbour.

The scale sets only the size of *Y*, not its sign: in every image, *Y* is positive (attraction) when more of its *B* cells have an *A* neighbour than expected and negative (avoidance) when fewer do. The alternative effect is the extra count: *O* is the total number of *A* neighbours of the *B* cells, and *Y* = (*O − E*)*/n*^*B*^ is the number of extra *A* neighbours per *B* cell, with variance *s*^2^ = *V/*(*n*^*B*^)^2^ under random labelling; *E* and *V* are again exact, as those of a total drawn without replacement [44]. The extra fraction is analogous to the cross-type nearest-neighbour function *G* at *r* [46], and the extra count to Ripley’s cross-*K* function [10]. The extra fraction measures how many *B* cells touch *A* cells, while the extra count also reflects how many *A* neighbours each *B* cell has. For a self-pair *A → A*, the *A* labels are shuffled among all cells, so its moments differ (Supplementary Section 3.8). For the extra fraction of a self-pair with a radius, one term of *V* counts the pairs of cells that share a neighbour, which takes time in proportion to the number of cells times the square of the number of neighbours per cell, so by default this term is estimated from 50 cells of each image and scaled up; the estimate of *V* is unbiased, and the same image always gives the same sample (Supplementary Remark 2). The count, and both effects with the *k* nearest neighbours, use the exact *V*. An image is not used for a pair when *V* = 0 or *R <* 1 (Supplementary Section 3.5).

### Combining images and patients

spicyR v2 combines the images of each patient, and the patients of each group, as a random-effects meta-analysis combines studies, with each image’s random-labelling variance 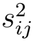 in the role of a study’s known sampling variance. Patient *i* has *J*_*i*_ images, and *m* patients are compared in all. Within a patient, image *j* gets the weight 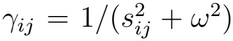, the inverse of its variance about the patient’s mean. An image with many cells has a small random-labelling variance 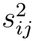, so it counts for more, but only up to a point: when the images of a patient genuinely differ (*ω*^2^ large), no image’s weight can exceed 1*/ω*^2^, so very large images count about equally. When they agree (*ω*^2^ near 0), the weights follow the random-labelling variances, and an image with many more cells counts for much more. The weighted mean of the images is the patient’s effect 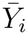.

Patients are weighted in the same way, *u*_*i*_ = 1*/*(1/Γ_*i*_ + *τ*^2^), where Γ_*i*_ = Σ_*j*_ *γ*_*ij*_, so a patient with more or larger images counts for more. When patients differ a lot (*τ*^2^ large), no patient’s weight can exceed 1*/τ*^2^, so patients count nearly equally. When patients agree (*τ*^2^ near 0), the weights follow the patients’ total information, and one patient with many large images can carry most of a group’s weight; a rule below on the number of patients that carry the weight guards against this. The group mean is

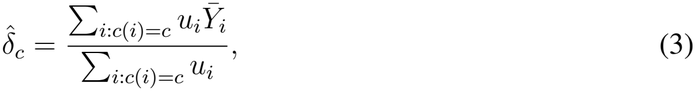

the GEE estimate under (1). *ω*^2^ and *τ*^2^ are estimated by the method of Paule and Mandel [47], shared across groups: 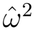 is the value at which the weighted spread of images within patients equals its expected value, Σ_*i*_(*J*_*i*_ *−* 1), and 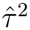 the value at which the weighted spread of patients within groups equals its expected value, *m −* 2 for two groups. 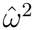 is set to 0 when the images of a patient spread no more than random labelling alone would give, and 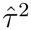 when patients spread no more than their images alone would give (Supplementary Section 4).

### Comparing groups

The groups are compared by the difference in their means, 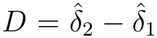. The random-labelling variances that set the weights can be wrong, for example when cells of a type aggregate, so the variance of *D* is instead estimated from how far the patients actually spread about their group’s mean. Each group’s variance *V*_*c*_ of 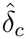 is estimated separately from its patients’ residuals 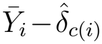, as a cluster-robust (sandwich) variance with patients as the clusters, so that patients are the units of inference. The plain sandwich variance is too small when a group has few patients, so we use the bias-reduced form of Bell and McCaffrey, known as CR2, which corrects this [48, 49]. We derived closed forms for this CR2 variance and for its Satterthwaite degrees of freedom under the two-level model (Supplementary Sections 5.4 and 5.6), so no general GEE solver is needed. The test is 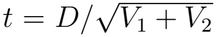 on Satterthwaite degrees of freedom [50]. A pair is not tested, and is reported with the reason, when a group’s variance cannot be estimated reliably:

- the group has fewer than two patients;
- the effect does not vary within either group (for example, no *B* cell has an *A* neighbour in any image), so the variance of *D* is zero;
- in one group, every image is at the end of the scale, with every *B* cell having an *A* neighbour (*Y* = 1) on the gain scale or none having one (*Y* = *−*1) on the loss scale, so that group’s spread is set by the scale, not measured;
- the weight rests on too few patients: the effective number of patients in group *c*, 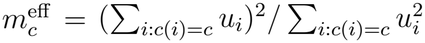, is below the smaller of 3 and half the *m*_*c*_ patients of the group that contribute;
- the Satterthwaite degrees of freedom are below 1. (Supplementary Section 5.8).

### Extensions

spicyR v2 also handles covariates, more than two groups, time-to-event outcomes such as survival, and several radii, all with the same GEE fit and CR2 variance.

#### Covariates and more than two groups

Covariates enter (1) as 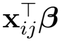, at the image or the patient level, and are estimated by the same GEE with CR2. One built-in covariate adjusts for abundance: the fraction of *B* cells with an *A* neighbour can differ between groups simply because *A* cells are more or less common in one group, so the log share of *A* cells among all cells of the image can be added as an image-level covariate, and the groups are compared at the same abundance. With more than two groups, the test is an overall Wald test that tests if all group means are equal, referred to an *F* distribution with the CR2 variance and the approximate Hotelling’s *T*^2^ correction of the degrees of freedom [49]; for two groups, *F* = *t*^2^ and the test is the *t*-test above (Supplementary Sections 6.1 and 6.2).

#### Time-to-event outcomes

For a time-to-event outcome, such as overall or disease-free survival, the question is whether a pair’s co-localisation is associated with how soon patients have the event. First, a Cox model [51] is fitted once, without the pair but with the pair’s covariates **w**_*i*_,

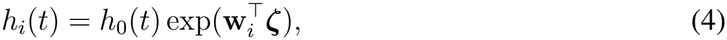

and gives each patient a martingale residual [52],

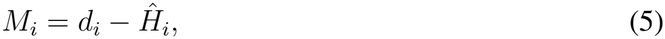

where *d*_*i*_ is 1 if patient *i* had the event (for example, died) and 0 if they were censored, and *Ĥ*_*i*_ is the number of events model (4) expects for patient *i* up to their last follow-up. So *M*_*i*_ is positive for a patient who had the event sooner than expected, and negative for one who had it later than expected or was censored without it. Model (1) is then fitted with *M*_*i*_ in place of the group means,

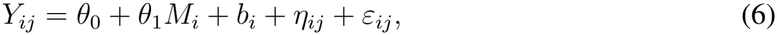

and *θ*_1_ = 0 is tested by the same GEE with CR2; a positive *θ*_1_ means that patients who had the event sooner than expected have higher co-localisation. Putting the effect on the left of the model, rather than into the Cox model as a predictor, lets each patient’s images enter with the same weights and CR2 variance as in the group comparison. *θ*_1_, the change in the mean effect per unit of *M*_*i*_, is hard to interpret, so spicyR v2 also reports a hazard ratio per standard deviation of the pair’s effect. It comes from a Cox model with the covariates **w**_*i*_ and each patient’s effect, shrunk towards the mean so that patients with few cells do not pull the hazard ratio towards 1. The hazard ratio only describes the size of the association, and the pair’s *p*-value is that of the test of *θ*_1_ (Supplementary Section 6.3). For a time-to-event outcome, patients with no follow-up time or event status are removed first.

#### Several radii

When the scale is unknown, several radii can be tested at once. Each radius gives its own *t* statistic, which is put on the normal scale because the degrees of freedom differ between radii, and the pair is summarised by the largest of these in absolute value, *z*_max_, whose *p*-value is the probability that the largest would be at least this large if the groups did not differ at any radius. spicyR v2 offers three methods to calculate *p*-values for this test. One comes from the joint normal distribution of the statistics, with their correlations estimated from each patient’s contribution to the cluster-robust variance at each radius [53, 54], and slows as radii are added. The other, which we call Davies’ bound, follows Davies [55]: it adds the chance that the statistic at the first radius exceeds *z*_max_ in absolute value to the expected number of times the statistics cross *z*_max_ or *−z*_max_ between neighbouring radii. It is the default because it is a closed form that takes microseconds per pair for any number of radii, against milliseconds to seconds for the joint *p*-value; it needs only the correlations between adjacent radii and is never smaller than the joint *p*-value, so it is slightly conservative relative to it (Supplementary Section 6.4). Because nearby radii count many of the same neighbours, their statistics are strongly correlated, and either *p*-value keeps much more power than a Bonferroni correction, though less than a single well-chosen radius. The third option, the Cauchy combination of the radii’s *p*-values [56], which needs no correlations, is also available.

### Software

spicyR v2 is the default method of the function spicy in the R package spicyR and in the Python package spicyr, which share one C++ core and give the same results to within a relative difference of 10^*−*7^ on a common set of test cases. Cells can be given as a data frame, as a SingleCellExperiment or SpatialExperiment object in R, or as an AnnData or SpatialData object in Python, and in either language as the path of a SpatialData or AnnData Zarr store, which is read a batch of images at a time so that the cells need never fit in memory. By default, spicy tests every ordered pair of cell types by the extra fraction at one radius, adjusts the *p*-values across pairs by the method of Benjamini and Hochberg [57] and reports each pair that is not tested with the reason. Options include the extra count, the *k* nearest neighbours, covariates and abundance adjustment, the Hartung–Knapp variance in place of CR2 [58], the exact variance of a self-pair’s fraction in place of the sampled one, and cell weights, such as a classifier’s confidence in each cell’s type. Supplementary Section 7 sets out the whole method as one algorithm, with the cost of each step.

## Datasets

### Datasets and comparisons

We assembled every spatial dataset we found with segmented and annotated cells and a question about two groups of patients, giving 74 two-group analyses of 42 public datasets (Supplementary Table S1). The datasets were measured by imaging mass cytometry (12 datasets), multiplexed ion beam imaging (9), CODEX or PhenoCycler (8), Xenium (5), MERFISH (4), CosMx (2), in situ sequencing (1) and multiplex immunofluorescence (1). Thirty-eight were of human and four of mouse tissue, and 28 were of cancer or pre-malignant tissue.

Each analysis compared two groups defined by one question, about tissue state (47 analyses), such as lung adenocarcinoma against squamous cell carcinoma, about patient outcome (23), such as response to treatment or recurrence, or about sex (4), where we expected few differences in co-localisation and which served as negative controls. Where a dataset supported several questions or levels of cell type annotation, each was a separate analysis.

spicyR v2 was also applied to three survival analyses of two cohorts, overall and disease-free survival in the discovery cohort of triple-negative breast cancer of Meyer et al. [59] (208 patients and 46 deaths, and 163 patients and 52 events) and overall survival in the lung cancer cohort of Cords et al. [60] (1,059 patients and 730 deaths; Supplementary Table S1).

The unit of analysis was the patient, or the donor or mouse, with images nested within units. In four analyses of heart transplant biopsies [61], some patients had biopsies in both groups, so the biopsies were the units, and every method treated a patient’s biopsies as independent. In the head and neck cancer cohort of Wu et al. [62], the deposited regions have no patient identifier, so regions with identical clinical fields were grouped, giving 18 units that each hold whole patients. The analyses ranged from 8 to 1,019 units (median 35). We used each dataset’s own segmentation, coordinates and cell type labels, without re-annotation, and removed cells with a missing group or cell type.

### Cell types and pairs tested

A cell type was tested if it had at least five cells in at least three images of each group. Cells of untested types were kept, so that every method counted neighbours and formed its null among all cells of the tissue. Every ordered pair *A → B* of tested cell types was tested, including pairs of a cell type with itself. The radius was 25 *µ*m, or the nearest number of pixels for the six datasets with coordinates in pixels (Supplementary Table S1). spicyR v2 used the fraction at a single radius, without abundance adjustment; where stated, the count, or the 10 nearest neighbours in place of the radius, was used instead.

## Methods compared

We compared spicyR v2 with 17 existing methods, in three groups by what each image is compared against (Supplementary Table S2). Every method was given the same cells, cell types and pairs, with a radius of 25 *µ*m (or its pixel equivalent) or the *k* = 10 nearest neighbours, and the same units as above in every comparison between groups. Other settings followed each method’s documentation, and departures from the package defaults, including changes we made to KAMP’s handling of self-pairs and to SPIAT’s handling of zero scores, are listed in Supplementary Table S2. Several methods give a score for each image but no test between groups; for these, we averaged each score over a patient’s images and compared the patient means between groups with a two-sample *t*-test (pooled variance), requiring at least four patients with a score. With their variants, this gave 28 methods and variants (four for spicyR v2), each run on real data and in the main simulation.

### Compared with shuffled labels

These methods compare each image with random relabellings of its own cells, the same reference as spicyR v2. Squidpy [18], COZI and SEA [22], Scimap [21], histoCAT [17], imcRtools [19], Giotto [20] and CellCharter [63] count the neighbours of each type around each cell and compare the counts with permutations of the labels within the image. For COZI, SEA, Scimap, histoCAT, imcRtools and Giotto we used the numbers of permutations of the benchmark of Schiller et al. [22], which are below the package defaults. Because a *z*-score for co-localisation grows with the number of cells *n* in an image even when co-localisation does not change, we also ran COZI and Squidpy with each image’s *z*-score divided by 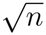, as Schiller et al. recommend for COZI. All of these methods except CellCharter give a score per image and were compared by the *t*-test; CellCharter compares groups directly, and we reimplemented its test to permute group labels between patients, so that all of a patient’s images move together. KAMP [40], which we implemented from the authors’ formulas, uses Ripley’s cross-type *K* at *r* minus its exact mean under random labelling, gives a score per image and was compared by the *t*-test. The smoppix package [27] uses the chance that a cell’s nearest partner is closer than a random other cell, and was compared by a weighted mixed model with a patient random effect.

### Compared with random points

spicyR v1 [12], PANORAMIC [15], SPOT [13], SpaceANOVA [25] and spatialFDA [14] compare each image with points placed at random across its window, through Ripley’s *L, G*, the pair correlation function or, for PANORAMIC, an edge-corrected local enrichment. Methods that integrate over distances used radii up to 50 *µ*m, or the range their authors recommend. Each has its own multi-sample test, and each was given the patient as the sample or random effect.

### Compared directly

SPIAT and DIMPLE compare the two cell types directly, with no chance model. SPIAT [64] uses a ratio of neighbour counts and was compared by the *t*-test. DIMPLE [65] uses the distance between smoothed maps of the two types’ density and was compared by a linear model of patient medians.

### Ordered and unordered pairs

Directional methods, such as spicyR v2, test *A → B* and *B → A* separately, whereas symmetric methods give one result for *A* and *B* together (Supplementary Table S2). Every comparison used ordered pairs, with a symmetric method’s result used for both orders.

## Data with controlled differences

We compared methods on data in which we controlled what differed between the groups, from our own tissue simulator, from a published simulator and from real cohorts that we altered.

### Evaluation measures

In each simulated study, the pairs whose co-localisation was changed in one group are the positives and the pairs whose co-localisation was not changed are the negatives. The power is the percentage of tests of positives with *p <* 0.05, and the false positive rate (FPR) the percentage of tests of negatives with *p <* 0.05, each with a 95% confidence interval computed from each study’s power and false positive rate (mean *±* 1.96 *·* standard errors over studies), because the pairs of a study share its tissue and patients. The area under the ROC curve (AUC), computed with *−* log_10_ *p* as the score, is the chance that a randomly chosen positive has a smaller *p*-value than a randomly chosen negative, with ties counted as one half, so 0.5 is no better than guessing and 1 separates them perfectly. Tests that a method did not return were left out, so each method has its own denominator.

### Simulated tissue

Real images differ widely in how much tissue they show and how many cells they hold, so we built a tissue simulator that reproduces this. Each image is a 1,000 *µ*m *×* 1,000 *µ*m square holding one to three overlapping discs of tissue with a gradient in cell density, with a median of 2,000 cells (about 500 to 8,000). Cells are labelled as tumour (60%), stroma (22%) or one of six rare types, *A* to *F* (3% each), and 0–90% of tumour cells are packed into nests at four times the mean density. To make *B* cells sit next to *A* cells, each *B* cell was moved, with probability *f*_*i*_, to a random point within 20 *µ*m of a randomly chosen *A* cell; this changes where the *B* cells are, but not how many there are. So that patients in a group differ, as real patients do, the probability varied between patients: logit(*f*_*i*_) = logit(*f*_*g*_) + *a*_*i*_, where *f*_*g*_ is the typical probability in the patient’s group and *a*_*i*_ ~ *N* (0, 0.5^2^).

### Power when information varies between images

An image with few cells measures co-localisation less precisely than an image with many, and spicyR v2 weights the images accordingly. This simulation asks whether doing so gives more power than other methods, without more false positives. We simulated 1,000 studies of 20 patients, 10 per group, with one image each. *B* cells were moved towards *A* cells with *f*_*g*_ = 0.10 in one group and 0.25 in the other, so *A → B* and *B → A* were the positives. The negatives were of two kinds: pairs that are co-localised equally in both groups (*F* cells moved towards *E* cells with *f*_*g*_ = 0.20 in both groups, giving *E → F* and *F → E*), and pairs that were not moved towards each other (*C → D, D → C, B → D* and *D → B*). Pairs of *A* and *C* were not scored, because moving *B* cells next to *A* cells can change the neighbourhoods of *A* cells indirectly. Every method was run at *r* = 25 *µ*m, or with the 10 nearest neighbours.

### Tissue artefacts

Groups of patients often differ in their tissue as well as in how their cells are arranged. One group’s samples may tear more, be imaged in smaller fields of view, have more holes, pack their tumour cells more densely or have more of a cell type. A method should not call a pair changed because of such differences. In each of the following designs, the tissue differed between the groups but co-localisation did not, and we report the percentage of tests with *p <* 0.05 among pairs whose arrangement did not change. Where a design gives several such rates, we report the largest, so that each value is the method’s worst case. For the simulated designs, this is the larger of the rates for pairs whose cells were not moved and for pairs co-localised equally in both groups.

### Simulated artefacts

#### Holes

In 200 studies from our simulator with 10 patients per group, holes covered 30–50% of the tissue in one group and 5–20% in the other.

#### Cell packing

In 200 studies from our simulator, 0–45% of tumour cells were packed into nests in one group and 45–90% in the other. In a second design of 500 studies, tested at *r* = 20 *µ*m, tumour nests held 150 cells in one group and 300, more densely packed, in the other, and 10% of immune cells were placed near tumour cells in both groups, so the anchor type of the pair tumour *→* immune was packed more densely in one group. We report the larger of the two rates.

#### Abundance

In 200 studies from our simulator without nests, each patient’s shares of the rare types were multiplied by a log-normal factor (standard deviation 1 on the log scale), cell types *D* and *F* were twice as common in one group, and the variation between patients in co-localisation was smaller (*a*_*i*_ *~ N* (0, 0.2^2^)).

#### Image size

In 200 studies from the in silico tissue simulator of Schiller et al. [22] (below), the images of one group were cropped to 30% of their area. As this tissue is uniform, cropping changed only the number of cells per image, unlike the cropped field of view of real tissue below. We report the larger of the rates at the weak and strong preferences.

### Artefacts added to real tissue

#### Cropped field of view

In two real cohorts [66, 67], patients were split at random into two pseudo-groups, stratified by their real group, and every image of one pseudo-group was cropped to a randomly placed window covering 30% of its area. Each cohort was split twice, and each split was used as it was and with a known change planted in the cropped pseudo-group, in which the 10% of cells of a type *A* farthest from a cell of type *B* swapped labels with as many cells of a type *C* nearest to *B* cells. False positives were counted among every pair without a planted change, and among the pairs involving neither *A* nor *C* with one, and the rate is averaged over the two cohorts.

#### Tears

In three real cohorts [7, 66, 67], every image of one group was torn by removing strips of cells 40 *µ*m wide until at least 40% of its cells were gone. Because these groups are the real ones, some pairs may truly differ, so the rate is the percentage of the pairs with *p ≥* 0.05 in the intact cohort that had *p <* 0.05 after tearing, averaged over the three cohorts.

### In silico tissue cohort

For the image-size design, we used the in silico tissue cohort of Schiller et al. [22], a simulator we did not design. In it, four cell types are assigned to packed cells (discs of radius 10 units) with a set neighbourhood preference, following Baker et al. [68]. We used their asymmetric design, in which type 0 cells take 45% or 60% of their neighbours from type 1, with the same preference in both groups and type 0 making up 25% of cells, and tested type 1 *→* type 0 at a radius of 45 units (about ten neighbours per cell). Each study had 10 patients per group with one image each, and each design was simulated 200 times.

## Evaluation on real data

On real data we do not know which pairs truly differ between the groups, so we compared each analysis with versions of itself in which the group labels were shuffled, and in which nothing can differ.

### Calibration with shuffled labels

A method is calibrated if, when the groups do not differ, it gives *p <* 0.05 in about 5% of tests. To check this on real tissue, we shuffled the group labels across patients 30 times in each analysis, keeping the group sizes and moving each patient’s images with the patient, so that any difference between the shuffled groups is due to chance. For a survival analysis, each patient’s follow-up time and event indicator were shuffled together across patients. Every method was given the same 30 shuffles. The cell types to test were chosen once, from the real groups, and kept for every shuffle. SpaceANOVA and spatialFDA were too slow to run on the shuffles of up to four of the largest analyses, which were left out for these methods only. We recorded two measures, each averaged over the analyses. The first is the percentage of tests with *p <* 0.05, which should be about 5%. The second is the percentage of shuffles with at least one discovery (a Benjamini– Hochberg adjusted *p <* 0.05), the chance that a study with no real difference reports a finding, which should be at most 5%.

### More changes than expected by chance

A calibrated method is useful only if it also finds real differences, so for each method and analysis we asked whether the real labels gave more pairs with *p <* 0.05 than the shuffles did. A method identified more changes than expected by chance in an analysis when the real count was larger than the count in every one of the 30 shuffles. If the groups did not differ, the real labels would be just one more shuffle, and the real count would be the largest of the 31 with probability at most 1/31 (3.2%). The groups can also differ in cell type composition or image size, which can raise the count as well. Identifying more changes than expected by chance therefore shows that the groups differ, but not necessarily in co-localisation, and the data with controlled differences measure how much such differences contribute to each method’s calls. Each dataset was given equal weight by averaging over its analyses first.

For spicyR v2, we related the changes found to cohort size through the ratio of changes found relative to chance, the number of pairs with *p <* 0.05 with the real labels divided by the median of the 30 counts of pairs with *p <* 0.05 from the shuffles (or by 0.5, if the median was smaller), so a ratio of 1 means no more pairs than chance. Where a study gave several analyses of the same question, for example at different levels of cell type annotation, only the one with the most patients was used. Cohort size was the number of patients in the smaller group, or, for a survival analysis, the smaller of the numbers of patients with and without an event.

### Power and sample size

Because spicyR v2 separates the variance within each patient from the variation between patients, and every variance has a closed form, a fitted model also gives the number of patients a study would need to detect a given change. For a pair, let *ū*_*c*_ be the mean weight *u*_*i*_ of the patients in group *c*, and *ū* = 2*/*(1*/ū*_1_ + 1*/ū*_2_) the harmonic mean of *ū*_1_ and *ū*_2_, and let *φ* be the ratio of the true variance of the difference *D* to the variance the model implies, estimated from the cohort’s CR2 variance by 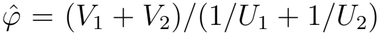 with *U*_*c*_ = Σ_*i*:*c*(*i*)=*c*_ *u*_*i*_, so that the calculation uses how much the cohort’s patients actually vary (*φ* = 1 when the model’s variance is accurate, and its estimate from a cohort is then close to 1). Assuming new patients resemble those in the cohort, the number of patients needed per group, for a two-sided test at level *α* with 80% power and equal group sizes, is then

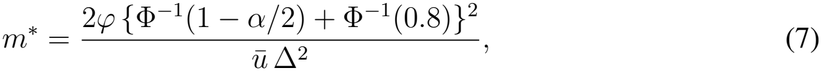

rounded up, where Δ is the difference between the group means and φ^*−*1^ is the inverse of the standard normal distribution function (Supplementary Section 6.5). Likewise, a cohort can detect, with 80% power at level *α*, a difference of at least 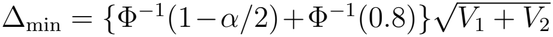 using its own CR2 variance. We set Δ = 0.10 on the scale of the extra fraction, meaning that the groups differ by 10 percentage points in the share of the *B* cells that would lack an *A* neighbour by chance but have one, on the gain scale, or that would have one by chance but do not, on the loss scale, and *α* = 0.05 or 0.001. We used the analyses with at least 20 patients in the smaller group, leaving out analyses which had the heart transplant biopsies as units. Where a study gave several analyses of the same question, the one with the most patients was used. For each cohort, we computed *m*^*\**^ for every pair and report the median over pairs, leaving out pairs with fewer than three patients in a group.

### Breast cancer case study

To see whether differences in abundance mislead methods in real tissue as they did in simulation, we examined one cohort in detail.

We used the Basel cohort of the imaging mass cytometry dataset of Jackson et al. [6], comparing 176 patients with HR+HER2-tumours with 49 with triple-negative tumours, with the authors’ 27 cell metaclusters, 24 of which passed the cell type rule. We examined two pairs. The first, metacluster 24 tumour cells *→* small elongated stromal cells, was chosen from the pairs that spicyR v2 discovered and no other method did, across all analyses. It was one of the pairs that passed every robustness check we applied, with a difference of the same sign at most other radii, with the extra count and with adjustment for abundance, few patients whose extra fraction was at its limit, and an analysis with more pairs at *p <* 0.05 than every shuffle. For this pair, 211 patients (168 and 43) were informative. The second, HR-low CK tumour cells *→* Vimentin-high stromal cells, was chosen from Squidpy’s discoveries in this analysis as an example in which the abundance of one cell type differed between subtypes and spicyR v2 found no change. To check that the first pair did not depend on the radius of 25 *µ*m, we also tested it at single radii from 10 to 50 *µ*m, and with 25 radii from 2 to 50 *µ*m combined into one test.

### Accounting for abundance

To ask how much of Squidpy’s result came from abundance, we compared patients’ mean Squidpy scores between subtypes by a linear model, without and with the patients’ mean log numbers of both cell types as covariates. For the second pair, we also compared subtypes within quintiles of the patients’ mean number of HR-low CK tumour cells, with the quintile as a covariate, so that only patients with similar numbers of these cells were compared. For both pairs, spicyR v2 was also fitted with the log share of the *A* cells in each image as an image-level covariate.

### Weighting of patients

spicyR v2 gives little weight to patients with few cells of the pair. To see how much this weighting contributed to the first pair, we also compared patients’ effects with every patient weighted equally, with and without the patient with the fewest cells, and by a Wilcoxon rank-sum test, and we also weighted Squidpy’s patient scores by spicyR v2’s patient weights.

## Computational performance

Cohorts now reach millions of cells per study and hundreds of patients, and a method that permutes labels in every image repeats its work for every permutation, whereas spicyR v2 computes the random-labelling moments once, in closed form. We therefore measured how the time and memory of each method grow with the number of cells.

### Synthetic studies

Each study had 50 images, with 1,000 to 2 million cells per image (50 thousand to 100 million in all), placed uniformly at random at 3,000 cells per mm^2^, so that larger images cover more tissue, and labelled with one of 10 cell types with equal probability. Every method tested all 90 ordered pairs of distinct cell types (10 *×* 9; no cell type was tested with itself) at *r* = 25 *µ*m or with the 10 nearest neighbours, on one core, with a limit of one hour. spicyR v2 was also run on eight cores, and on one core reading the cells from disk, from a SpatialData Zarr store [69] written by spicyR, with the time to read the cells included. We recorded the wall time of the method call and the peak memory of its processes, sampled every 0.1 s and including the data and the interpreter. Methods that run in Python, and spicyR v1, which runs in its own R library, were timed within their own process, so their times exclude handing the cells over from R and starting the interpreter. Each method ran alone on a node of the Gadi supercomputer (National Computational Infrastructure, Australia; two 24-core Intel Xeon Platinum 8274 processors, 190 GB of memory).

### Real data

To check these timings on real tissue, every method was timed once on one core on each of the 74 analyses it was run on, of 42 thousand to 5.5 million cells, with a limit of 24 hours.

## Statistics and reproducibility

All tests were two-sided, and a discovery was a pair with a Benjamini–Hochberg adjusted *p <* 0.05 [57] within its analysis. No statistical method was used to predetermine sample size, the experiments were not randomised and the investigators were not blinded, as the datasets and their groups came from published studies. All analyses used spicyR v2 (R package spicyR, development version 1.99.56, released in Bioconductor as version 2.0) and spicyR v1 (version 1.24.1) in R 4.6.1, and the versions of the other packages are listed in Supplementary Table S2.

## Results

### spicyR v2 weights each image by how precisely it measures co-localisation

Here we present spicyR v2, a method for testing changes in cell type co-localisation between groups of patients. Briefly, spicyR v2 assumes that cells have been segmented and assigned to cell types, and defines neighbours as the cells within a chosen radius or the *k* nearest cells. For each pair of cell types, it compares each image with a random labelling of its own cells (Figure 1a). Its score, the extra fraction, is the share of cells of one type with a cell of the other type nearby, in excess of, or short of, that expected under this random labelling. Because the expected share comes from the cells actually imaged, differences in the size, shape and cell content of images are not mistaken for co-localisation. An alternative score, the extra count of neighbours per cell, keeps its power when almost every cell already has a neighbour but, unlike the extra fraction, also rises when the other type is packed more densely (Figure 1a, right; Supplementary Proposition 4). The precision of each image is set by the variance of its extra fraction under random labelling, which depends on how many cells of each type the image contains and how they are arranged. spicyR v1 approximated this variance from the numbers of cells alone [12]. In contrast, spicyR v2 computes this variance exactly from the neighbourhood of every cell, using classical results for sampling without replacement [44] and without any permutations (Figure 1b; Methods, Supplementary Section 3). A two-level model then combines images within patients and compares patients between groups, weighting each image by its precision as a meta-analysis weights its studies (Figure 1c). Imprecise images therefore add little noise, and the variation between images caps the weight any one image can carry. A cluster-robust variance, which we derived in closed form with its degrees of freedom (Supplementary Section 5), is fast in large cohorts and kept the test calibrated in small ones. spicyR v2 reports a test for every pair of cell types in both directions, and the same model can combine several radii into one test, relate co-localisation to survival and can give the number of patients needed to detect a change (Figure 1d).

**Figure 1:**
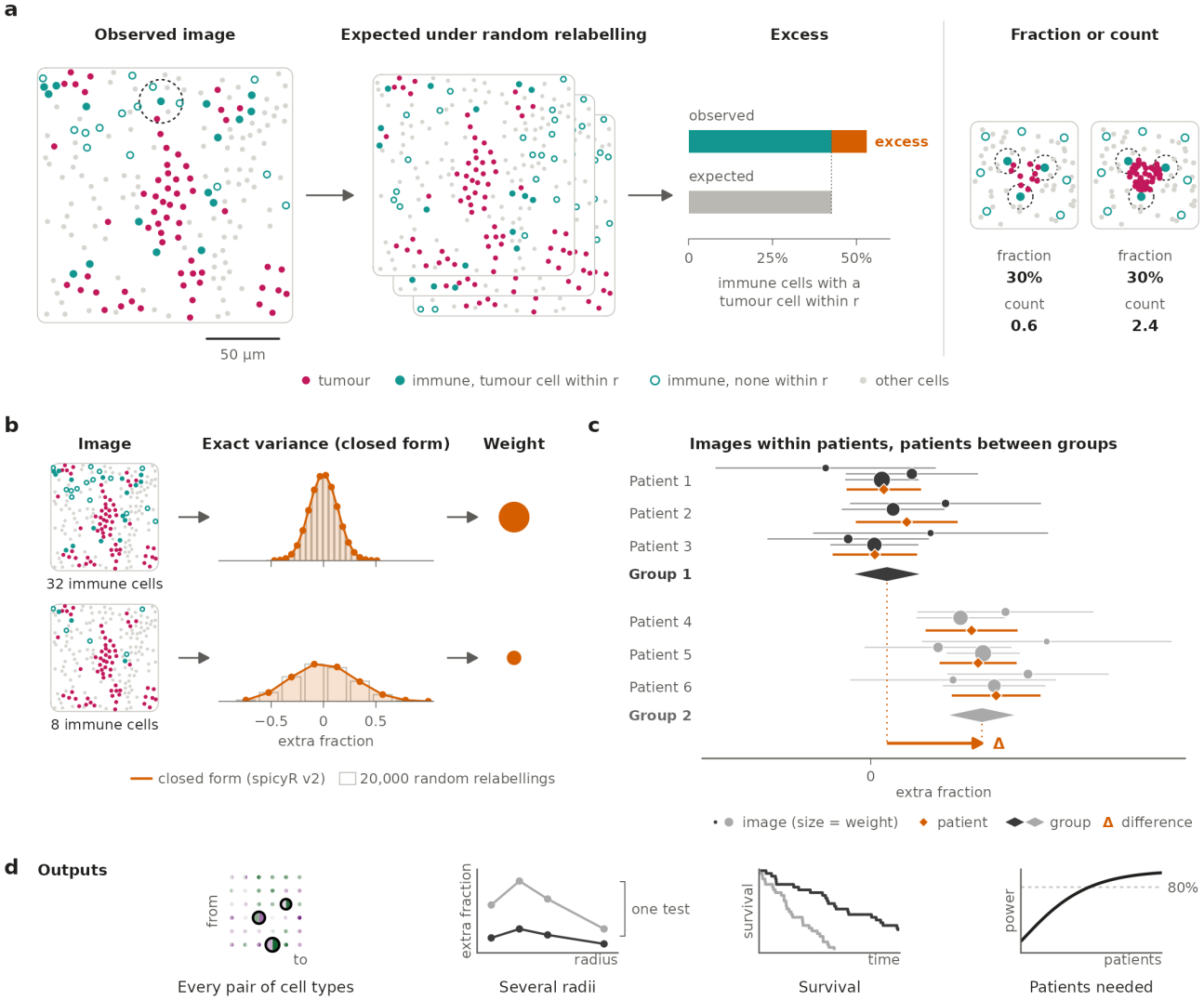
spicyR v2 scores each image against random labelling, weights it by its exact precision and compares patients between groups. (a) Each image is scored by counting the immune cells with a tumour cell within a radius *r* (dashed circle, *r* = 15 *µ*m), against the number expected if immune labels were shuffled among the non-tumour cells, with every cell kept in place. The excess, divided by the immune cells that could still gain a tumour neighbour, is the extra fraction. Image from a breast cancer imaging mass cytometry core [6]. Right, packing tumour cells more densely raises the number of tumour neighbours per immune cell (0.6 to 2.4) but not the share of immune cells with a tumour neighbour (30% in both). (b) Each image is weighted by how much its score varies under random labelling, which spicyR v2 calculates exactly from a formula (orange) rather than by shuffling (grey, 20,000 shuffles). An image with 32 immune cells varies less, and gets a larger weight (disc area), than one with 8. (c) Images are combined into patient estimates (orange diamonds) and patients are compared between groups (grey diamonds), as in a meta-analysis, with patients as the unit of the test. Lines, 95% intervals; *D*, difference between groups. (d) Outputs: tests of every ordered pair of cell types (anchor type *A*, rows; tested type *B*, columns), a single test combining several radii, associations with survival and the number of patients needed to detect a change. Values in (c) and (d) are illustrative.

### spicyR v2 is more powerful than spicyR v1 in simulated studies

To test whether modelling the precision of each image improves power, we simulated 1,000 studies of 20 patients whose images differed in their numbers of cells, in how densely the cells were packed and in the shape of the tissue. These differences can all affect how precisely an image measures co-localisation. In one group, a larger share of the cells of one type was moved close to another type, while other pairs did not change. With either of its scores, spicyR v2 separated changed from unchanged pairs better than every other method and variant we compared (Figure 2a). Its extra fraction had an area under the ROC curve (AUC) of 0.87, against 0.80 for spicyR v1 (Figure 2b). PANORAMIC and smoppix, which also model the variance of each image, came next (AUC 0.82 and 0.81). At *p <* 0.05, spicyR v2 had 1.74 times the power of spicyR v1 (53% v 30%), and both were slightly conservative (false positive rates 4.2% and 3.1%; Figure 2c). Together, these results show that in simulations spicyR v2 has more power than spicyR v1 without inflating the false positive rate.

**Figure 2:**
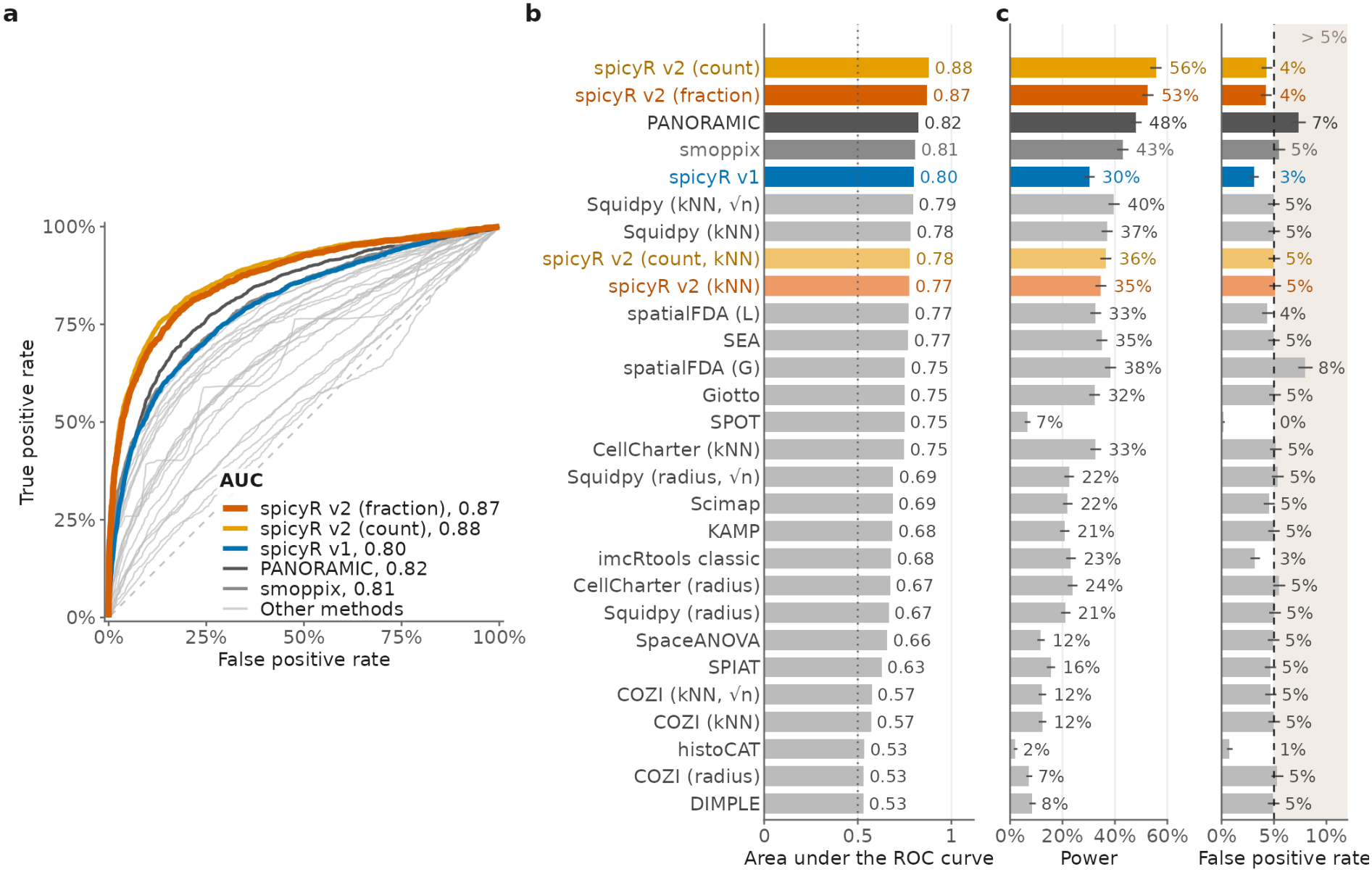
spicyR v2 is more powerful than spicyR v1 at a calibrated false positive rate when precision varies between images. Simulated studies of 20 patients (one image per patient, 10 per group) in which the number of cells varies between images (median coefficient of variation 0.60 over 1,000 studies). (a) ROC curves over the 1,000 studies (2,000 tests of pairs whose co-localisation changed and 6,000 of pairs that did not), with the area under each curve. spicyR v2 with the extra fraction (orange) and the extra count (amber), spicyR v1 (blue), PANORAMIC (dark grey), smoppix (mid grey) and 23 other methods, including variants of a few (light grey). Methods without their own test between groups (Squidpy, COZI, SEA, Scimap, histoCAT, imcR-tools, Giotto, KAMP and SPIAT) were compared by a *t*-test of their per-image scores (Methods); COZI answers a different question from the one simulated here (Discussion). (b) Area under the ROC curve for each method. (c) Power (changed pairs) and false positive rate (unchanged pairs) at *p <* 0.05, with 95% confidence intervals; dashed line and shading, false positive rates above 5%.

### spicyR v2 is calibrated and well powered in real data

To see whether these results hold on real data, we applied 28 methods and variants to 74 two-group analyses of 42 datasets, and spicyR v2 to three survival analyses (Supplementary Tables S2 and S1). Most analyses compared tissue states (47), such as diseased against healthy tissue or two subtypes of a cancer. Others compared patient outcomes (23), such as response to treatment or recurrence, and four compared the sexes as potential negative controls. The methods differed widely in how many discoveries they made (Figure 3a; Supplementary Fig. S2). At a Benjamini– Hochberg false discovery rate of 5%, spicyR v2 made 846 discoveries across all analyses. Of the 24 methods other than spicyR v2 and its four variants, 19 made more discoveries than spicyR v2, and spatialFDA’s *G* function and PANORAMIC made the most (5,589 and 4,995). However, a method can make more discoveries simply by making more false ones.

**Figure 3:**
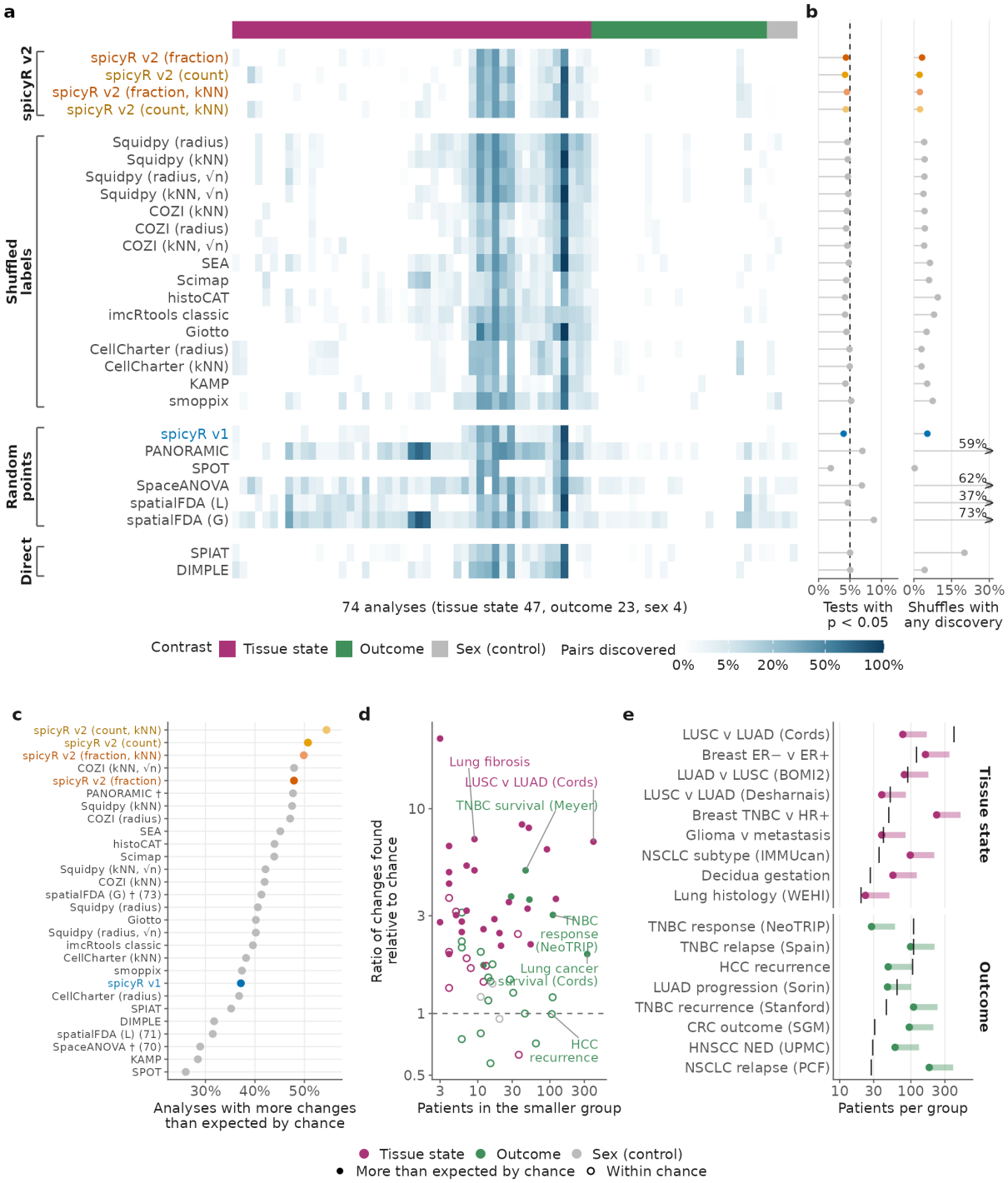
On real data, spicyR v2 is calibrated and identifies more changes than expected by chance in more datasets than any other method. (a) Percentage of cell type pairs discovered (BH-adjusted *p <* 0.05) by 28 methods (rows: spicyR v2, then the other methods grouped by what each image is compared against, either shuffled labels or random points, or compared directly with no reference) in 74 two-group analyses of published imaging cohorts (columns: 47 tissue-state, 23 outcome and 4 sex comparisons, clustered by their discoveries within each type; top strip, tissue state raspberry, outcome green, sex grey; white to dark blue, 0% to 100% of pairs). (b) The same analyses after shuffling patients between groups (30 shuffles per analysis): percentage of pair tests with *p <* 0.05 (left; dashed line, 5%) and percentage of shuffles in which at least one pair was discovered (BH-adjusted *p <* 0.05; right), averaged over each method’s analyses across datasets. Arrowheads mark values beyond the axis. (c) For each method, the percentage of analyses in which the real groups gave more pairs with *p <* 0.05 than every one of the 30 shuffles, averaged within each dataset and then over datasets. Orange, spicyR v2 and its variants; blue, spicyR v1; grey, other methods; †, more than 6% of tests with *p <* 0.05 under shuffled labels; in brackets, the number of analyses for methods that did not run in all 74. (d) For spicyR v2, the number of pairs with *p <* 0.05 in each analysis, divided by the median of the 30 counts of pairs with *p <* 0.05 from its shuffles (vertical axis; dashed line, 1, as many as the median shuffle), against the number of patients in the smaller group (horizontal axis; for survival, the smaller of the numbers of patients with and without an event). One point per study and comparison, including three survival analyses, coloured by contrast type. Filled points, analyses in which the real groups gave more pairs with *p <* 0.05 than every one of the 30 shuffles; open points, analyses in which they did not. (e) Patients needed per group to detect a change of 0.10 in the extra fraction with 80% power, for cohorts with at least 20 patients per group. For each cohort, this number was computed for every pair, and the median over pairs is shown. Dot, significance level 0.05; bar end, 0.001. The ticks are the number of patients in the cohort’s smaller group.

Because the true changes in real data are unknown, we repeatedly shuffled patients between groups, so that nothing truly differed between them. Any discovery after shuffling is therefore a false positive (Figure 3b; Supplementary Fig. S3). As in the simulation, spicyR v2 was slightly conservative, rejecting 4.4% of tests at *p <* 0.05 and discovering at least one pair (BH-adjusted *p <* 0.05) in 3.2% of shuffles. Of the 24 other methods, 18 were also calibrated, rejecting fewer than 5% of tests. The exceptions included spatialFDA’s *G* function (8.8%) and PANORAMIC (7.0%). These two methods made a discovery in 73% and 59% of shuffles, respectively. That is, many of their additional discoveries would be expected even if nothing differed between groups. However, of the 19 methods that made more discoveries than spicyR v2, 13 were calibrated. Their additional discoveries are therefore likely to reflect real differences between the groups. Despite making fewer discoveries, spicyR v2 identified more changes than expected by chance in more datasets than any other method (Figure 3c). That is, the real groups gave more pairs with *p <* 0.05 than every one of 30 shuffles, which would happen in only 1 analysis in 31 if the groups did not differ. Its variants did so in 48–54% of analyses, against at most 48% for other methods and 37% for spicyR v1. Even the most sensitive methods did so in only about half of the analyses, suggesting that whether a change can be found depends more on the dataset than on the method.

### Most spatial omics studies of patient outcome are underpowered

For every method, nearly all discoveries came from comparisons of tissue state rather than of patient outcome (Figure 3a). We classified a comparison as one of tissue state when its groups already differed at the time of sampling, such as diseased against healthy tissue or two subtypes of a cancer. We classified it as one of outcome when its groups differed in what happened to the patients afterwards, such as response to treatment or recurrence. On average, methods made discoveries in 6.2% of pairs in tissue-state analyses, against 0.28% in outcome analyses, a rate similar to that in the sex controls. Many tissue states, such as cancer subtypes, are defined by how the tissue looks, so they are likely to differ in co-localisation. To see whether outcome studies lacked a difference or lacked power, we compared each analysis’s ratio of changes found relative to chance with the number of patients in its smaller group (Figure 3d; Supplementary Fig. S4a). This ratio is the number of pairs with *p <* 0.05 divided by the median of the counts of pairs from its 30 shuffles. Taking one analysis per study and comparison, spicyR v2 identified more changes than expected by chance in 25 of 35 tissue-state comparisons, against 3 of 20 outcome comparisons other than survival. Many tissue-state comparisons did so with fewer than ten patients per group, whereas most outcome comparisons did not, even with more than 100. Survival was the exception, as all three survival analyses exceeded every shuffle. This suggests that changes in co-localisation related to outcome are still usually too small for current cohorts to detect.

Most outcome cohorts were too small to detect a 10% change in co-localisation (Figure 3e; Supplementary Fig. S4b,c). spicyR v2 can calculate the number of patients needed to detect a change, because it separates the variance within each patient from the variation between patients and every variance has a closed form. We took a 10% change to be a difference of 0.10 in the extra fraction between groups. Only eight of the 20 outcome comparisons had at least 20 patients in each group. In these eight, detecting this shift at *α* = 0.05 needed 28–181 patients per group, and four had at least that many. At *α* = 0.001, closer to the thresholds needed when many pairs are tested, only two had enough: the NeoTRIP trial in triple-negative breast cancer and a study of recurrence in hepatocellular carcinoma. Together, these results suggest that most outcome studies can detect only large changes in co-localisation, and that new studies should calculate the number of patients they need before collecting tissue.

### spicyR v2 is robust to experimental artefacts that mislead other methods

Shuffling patients in real data shows whether a method is calibrated, but not whether its dis-coveries reflect co-localisation or other differences between the groups’ tissue. To test this, we created differences between groups in which the tissue changed but the placement of cell types did not (Figure 4). We changed the shape of the tissue, by tearing or cropping the images of existing datasets or by simulating holes, or the number of cells, by simulating larger images, more densely packed cells or more abundant cell types. Every method and variant except spicyR v2 with the extra fraction and a radius had a false positive rate above 10% under at least one of these changes. Which change misled a method depended on how it defines what is expected by chance.

**Figure 4:**
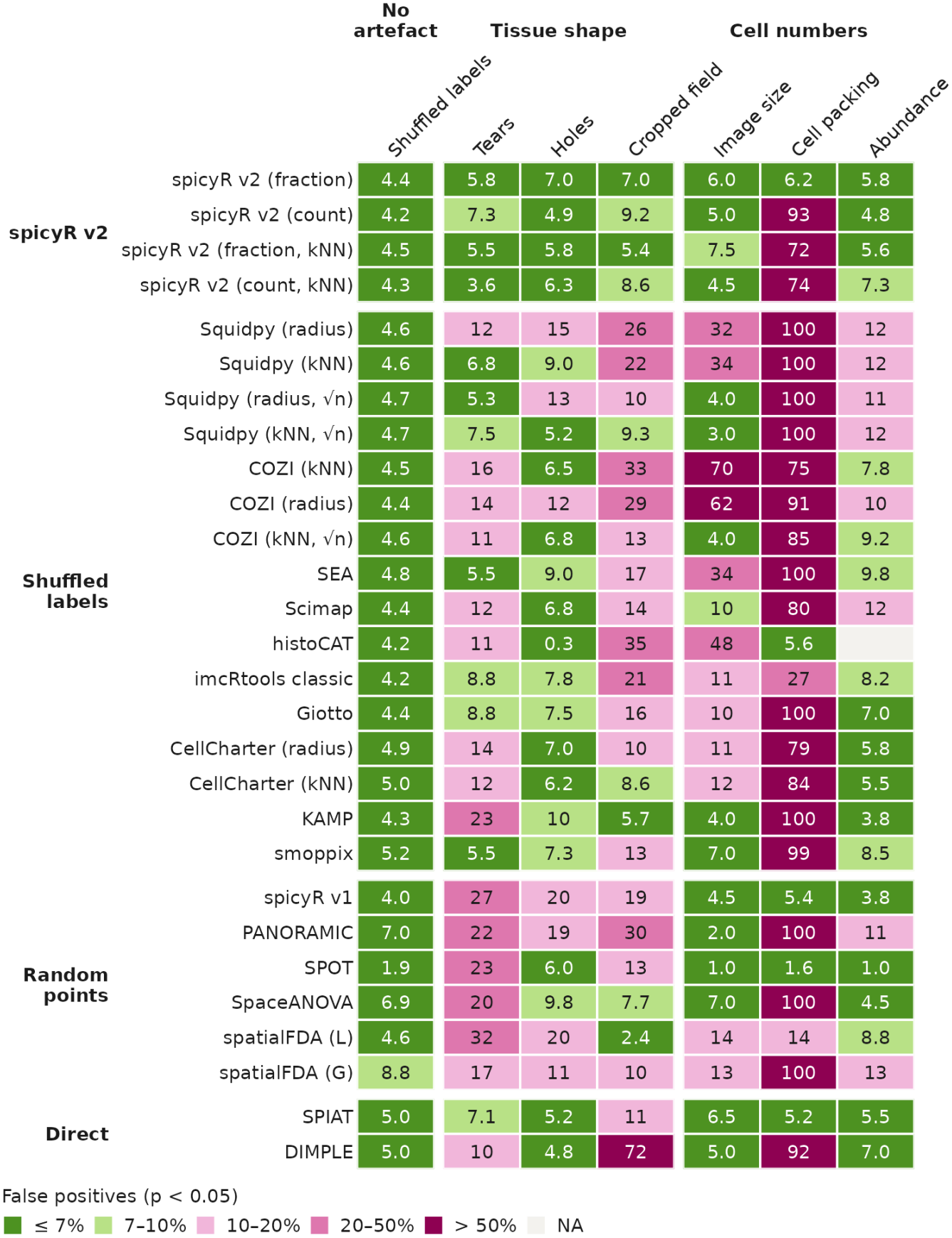
spicyR v2 is robust to differences in tissue shape and cell number. Percentage of false positives (*p <* 0.05 for pairs whose arrangement did not change) when the groups differ only in tissue shape (tears and cropped fields of view in existing datasets, simulated holes) or in cell numbers (simulated image size, packing of cells and abundance of cell types). The first column, patients shuffled between groups in the 74 real analyses (as in Fig. 3b), gives each method’s false positive rate when the groups differ in nothing.

Methods whose scores are scaled by the area of the image window were susceptible to tears and holes. When tears were more common in one group, spicyR v1, KAMP and PANORAMIC had false positive rates of 22-27%, and holes had a similar effect (19-20% for spicyR v1 and PANORAMIC). For these methods, the empty space of a tear or a hole still counts as tissue, which changes the expected neighbourhood although the cells have not moved.

In contrast, methods that permute cell type labels and report a *z*-score were susceptible to differences in cell number. When images were larger in one group, Squidpy’s radius and nearest-neighbour tests had false positive rates of 32% and 34%, because a *z*-score grows with the number of cells even when the excess does not. Dividing the *z*-score by 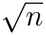 corrected for image size (4.0% and 3.0%), but not for how densely cells are packed (99.6% for the radius test) or for abundance (12%).

spicyR v2 with the extra fraction and a radius kept false positive rates close to 5% under every change (at most 7.0% at *p <* 0.05). Because it compares each image with random labelling of the cells actually imaged, a tear, a hole or a larger image changes the observed and the expected neighbourhoods alike. The extra count reported a change in 93% of tests when only packing differed, because packing one cell type more tightly gives each cell of the other type more neighbours, which is what the count measures. With *k* nearest neighbours rather than a radius, spicyR v2 also reported a change in 72% of tests when only packing differed, probably because a neighbourhood of *k* cells covers less space where cells are packed more densely. However, the extra count was slightly more powerful than the fraction in the simulation of Figure 2 (AUC 0.88 v 0.87). Together, these results suggest that the extra fraction is the safer default, and that the extra count is a slightly more powerful alternative when packing is unlikely to differ between groups.

### Abundance changes confound changes in co-localisation for other methods

The simulations of Figure 4 changed abundance between groups, but in real data the abundance of a cell type can also vary many-fold between patients of the same group. Since Squidpy’s *z*-score grows with the number of cells, this variation in abundance within a group can mask a real change in co-localisation. Conversely, a difference in abundance between groups can confound the comparison and create a false one. Both happened in the Basel breast cancer imaging mass cytometry cohort of Jackson et al. [6], when we compared 43 triple-negative informative patients with 168 HR+HER2-patients (Figure 5).

**Figure 5:**
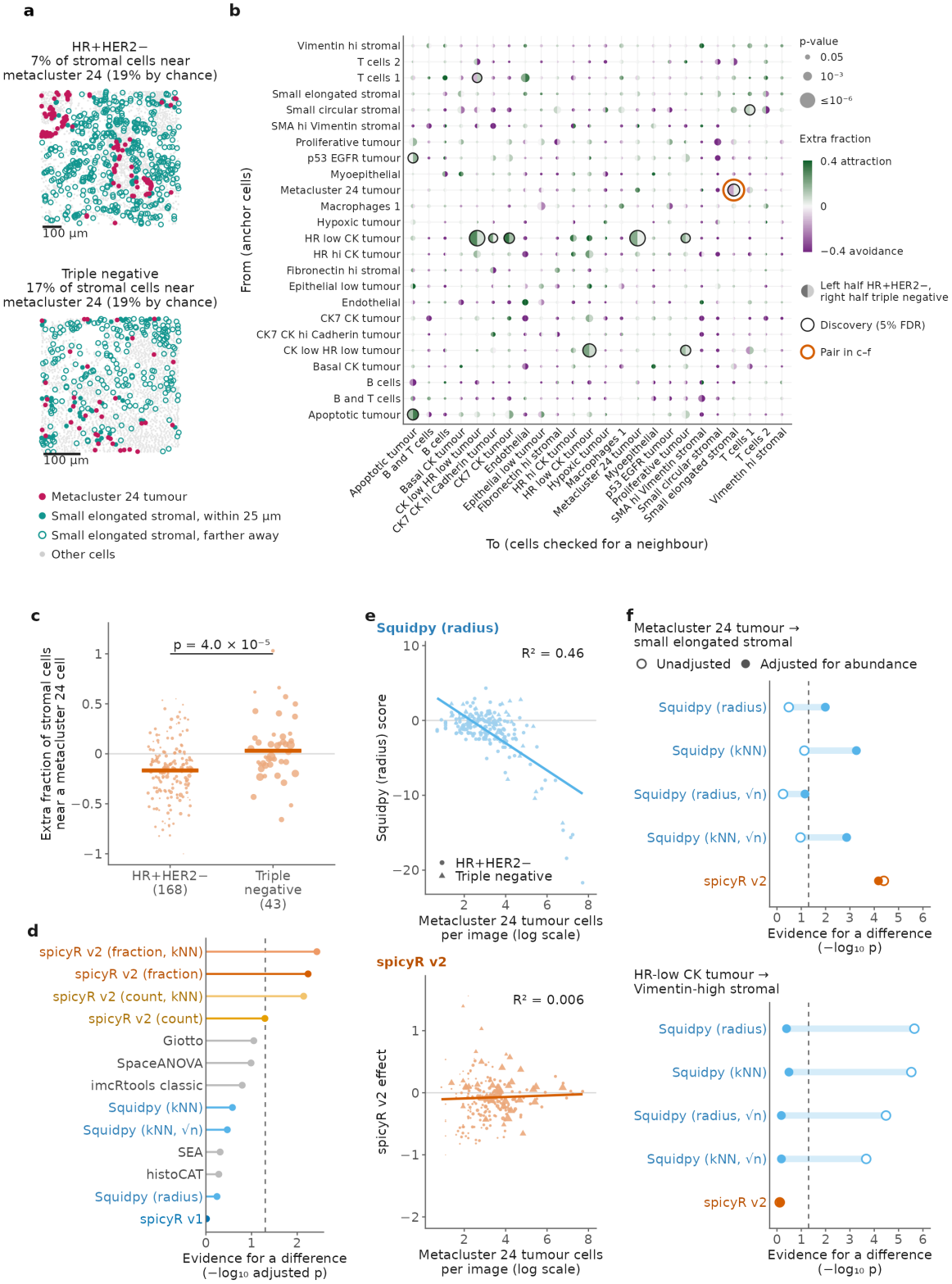
In breast cancer, spicyR v2 separates changes in co-localisation from differences in abundance. (a) Example images from an HR+HER2- and a triple-negative breast tumour [6]: metacluster 24 tumour cells (crimson), small elongated stromal cells within 25 *µ*m of one (filled teal) or farther away (open teal), and other cells (light grey). The two images were chosen with the same share of stromal cells expected to have a metacluster 24 neighbour by chance (19%), so they can visually be compared directly; the HR+HER2-image shows stronger avoidance than its group’s estimate (c), and the triple-negative image is close to random, as is its group. Above each image is the share of stromal cells with a metacluster 24 cell within 25 *µ*m, with the share expected by chance in brackets. Scale bars, 100 *µ*m. (b) Every ordered pair of the 24 cell types tested by spicyR v2 in this comparison: the halves of each disc give the two groups’ extra fractions (left, HR+HER2-; right, triple negative), disc size the p-value, black outlines the 12 discoveries at a 5% false discovery rate, and the orange ring the pair in (a, c–f). (c) Extra fraction of small elongated stromal cells with a metacluster 24 cell within 25 *µ*m, beyond that expected by chance, in 168 HR+HER2- and 43 triple-negative patients (each point is one patient, sized by its weight in the group estimate); orange bars, group estimates from spicyR v2. (d) Evidence for a difference for this pair by method (BH-adjusted within the analysis): the four spicyR v2 variants, the eight strongest other methods and spicyR v1; dashed line, adjusted *p* = 0.05. (e) Patient means of Squidpy’s radius score (top) and spicyR v2’s effect (bottom) against metacluster 24 abundance; lines are least-squares fits, with the variance explained (*R*^2^); circles, HR+HER2-patients; triangles, triple-negative patients. (f) p-values of Squidpy and spicyR v2 for a difference between groups, for metacluster 24 tumour to small elongated stromal cells (top) and for HR-low CK tumour to Vimentin-high stromal cells (bottom), without (open) and with (filled) adjustment for abundance (Methods); dashed line, *p* = 0.05. Squidpy called the bottom pair and spicyR v2 did not.

Variation in abundance between patients masked a change in co-localisation in Squidpy’s scores, but not in spicyR v2’s. Small elongated stromal cells avoided metacluster 24 tumour cells in HR+HER2-tumours, where on average 17% fewer had a metacluster 24 neighbour than expected by chance (extra fraction *−*0.17), but were placed close to random in triple-negative tumours (0.03; difference 0.20, 95% CI 0.11 to 0.29, *p* = 4.0 *×* 10^*−*5^; Figure 5b,c). In the two example images, a similar share of stromal cells would have a metacluster 24 cell within 25 *µ*m by chance (19% in both), but 7% did in the HR+HER2-tumour and 17% in the triple-negative tumour (Figure 5a). No other method found this pair at a 5% false discovery rate (Figure 5d). We chose this pair as a case study for that reason, and asked why the other methods did not detect it. The pair was a discovery at a 5% false discovery rate at every radius from 15 to 50 *µ*m, though not at 10 *µ*m, and at 25 *µ*m it was the highest-ranked pair that involved a non-tumour cell type (Supplementary Fig. S6). It was also a discovery under the omnibus test, which combines radii from 2 to 50 *µ*m into a single test (adjusted *p* = 0.020). Metacluster 24 cells were similarly common in both tumour subtypes but varied about 100-fold between patients. Squidpy’s *z*-score for this pair tracked this abundance (*R*^2^ = 0.46), while the extra fraction of spicyR v2 did not (*R*^2^ = 0.006; Figure 5e). Much of the variation in Squidpy’s score between patients therefore reflected how many metacluster 24 cells a patient had, not where they sat.

Accounting for the abundance of both cell types gave Squidpy evidence for the pair, for example for its nearest-neighbour test divided by 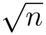 (*p* = 0.11 before and 1.4*×*10^*−*3^ after; Figure 5f, top; Supplementary Fig. S5a). spicyR v2 found the pair without adjustment, because its extra fraction is measured relative to the cells imaged and because its precision weights gave little weight to patients with a handful of these cells, such as one HR+HER2-patient with only 8 metacluster 24 and 75 stromal cells (Supplementary Fig. S5a).

A difference in abundance between subtypes created an apparent change in co-localisation for Squidpy, but not for spicyR v2 (Figure 5f, bottom). Squidpy found that HR-low CK tumour cells avoided Vimentin-high stromal cells more in HR+HER2-than in triple-negative tumours, a discovery at a 5% false discovery rate for each of its versions, including those divided by 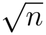. However, HR+HER2-tumours had about 10 times as many of these tumour cells (patient medians 71 and 7 per image), and Squidpy’s score fell with their number in both subtypes (Supplementary Fig. S5b). Accounting for abundance removed the difference for every version of Squidpy (*p ≥* 0.32), and patients with similar numbers of these tumour cells did not differ between subtypes (*p* = 0.90). spicyR v2 found no change with or without accounting for abundance (*p* = 0.77 and 0.71), as Vimentin-high stromal cells avoided these tumour cells to a similar extent in both subtypes (extra fraction *−*0.34 and *−*0.37). These two pairs show that in real data, differences in abundance can both mask real changes in co-localisation and create false ones for a score that grows with the number of cells. Together with the simulations of Figure 4, these results show that a calibrated test can still be misled by the shape of the tissue and the number of cells, and that spicyR v2 was robust to every difference we tested.

### spicyR v2 analyses large studies in minutes

Detecting modest changes in co-localisation needs cohorts of hundreds of patients (Figure 3e), and therefore a method that scales. Because each image’s variance and the test between patients both have closed forms, spicyR v2 needs no permutations and scales to very large studies (Supplementary Fig. S1). In synthetic studies of 50 images and 10 cell types, spicyR v2 analysed 100 million cells on one core in 105 s, less time than any other method took for 10 million cells; of the other methods, only spicyR v1 (32 min, 98 GB of memory) and KAMP (53 min, 51 GB) analysed 100 million cells within an hour (Supplementary Fig. S1a). With eight cores it took 26 s, and reading cells from disk rather than holding them in memory kept its peak memory to 2.0 GB (Supplementary Fig. S1a,b). On the 74 real analyses, it needed a median of 1.5 s per million cells, about 80 times faster than Squidpy and 110 times faster than spicyR v1 (Supplementary Fig. S1c). Together, these results show that spicyR v2 can analyse cohorts of the size needed to detect modest changes in co-localisation, with the memory of a laptop.

## Discussion

Here we have presented spicyR v2, a method for testing changes in co-localisation between groups of patients. It scores each image against a random labelling of its own cells [16] and, building on the pooling of replicated point patterns [33, 34], weights each image by how precisely it measures co-localisation. This precision comes from the exact variance of the extra fraction under random labelling, which depends on the arrangement of the cells rather than their number alone and is computed in one pass over the cells; with a closed-form test between patients, spicyR v2 needs no permutations. In simulations this gave spicyR v2 more power than spicyR v1 [12], and on public data it remained calibrated and identified more changes than expected by chance in more datasets than other methods. Unlike other calibrated methods, it was robust to tears, holes, cropping and differences in cell number, and was fast enough to test every pair of cell types in studies of 100 million cells in minutes. Complementing the neighbourhood analyses that many studies already perform, spicyR v2 now makes hypothesis-driven analysis of co-localisation reliable and fast enough to become standard in spatial omics research.

Methods for co-localisation differ mainly in what they expect by chance, and none of these choices are wrong, because each answers a different question. spicyR v2 keeps cells of one type in place and shuffles the labels of the remaining cells among them, asking whether cells of a second type sit closer to the first than the other cells in the same tissue do. For example, when a cell type is more abundant in one group of patients, every cell has more neighbours of that type. spicyR v2 reports a change only if cells of one type gain these neighbours more than the other cells in the same tissue do. Because the imaged cells stay in place, tears, holes and differences in density are held fixed, which suits comparisons between patients [9]. A change reported by spicyR v2 is therefore a change in where cells of one type sit relative to the other cells of the same tissue, not evidence that the cells interact or a change in how many of them there are. Methods that compare cells with points placed across the image window [10, 11] instead ask whether cells sit closer than expected anywhere in the image. This is the right question when empty space, such as the lumen of a duct or vessel, is part of the biology rather than an artefact. Methods like COZI [22] and Kontextual [23] ask different questions again. COZI looks only at cells that already touch another type and asks how densely they are surrounded by it. Since our simulations changed how many cells of one type sit near another, not how densely they are surrounded, the power of COZI in them does not measure what it was designed to find, though its calibration under shuffled labels still applies. Kontextual asks whether a cell of one type sits nearer to another type than the rest of its parent population (for example, other T cells for a CD8 T cell) does, which removes the influence of the tissue structure that the parent population defines, and was not part of our comparison. Many of the methods we compared, including Squidpy and COZI, were designed to find co-localisation within a single image, and the relationships they detect may well matter, so we included them by comparing patients’ mean scores with a *t*-test, which treats patients as the units of inference [70]. Their results therefore reflect this test as well as their scores, and the choice of score matters. A permutation *z*-score [17, 18] is well suited to asking whether cells in one image are arranged non-randomly, because it measures the evidence against random arrangement. However, compared between groups of images as if it were an effect size, it mixes how strongly cells co-localise with how much evidence each image provides, so a larger or more densely packed image scores higher with the same arrangement. Dividing it by 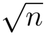, as recommended for COZI, corrects for image size but not for packing or abundance. A method for comparing patients should therefore be chosen both for the spatial relationship it quantifies and for what it expects by chance, as together these decide whether artefacts are mistaken for co-localisation.

Despite its strengths, spicyR v2 has some limitations. The extra fraction loses power at large radii, where almost every cell already has a neighbour of the other type. There, the extra count, which keeps its power but responds to how densely cells are packed, or methods that summarise many distances [14, 15] may be the better choice. Results also depend on the radius or choice of *k* nearest cells, which sets the relationship being tested: in the breast cancer case study, the change was found from 15 to 50 *µ*m but not at 10 *µ*m. The radius should therefore be chosen for the relationship of interest, such as a contact distance for cells that touch. When the scale is uncertain, testing several radii together protects against missing a change that occurs at a different radius, though it can have less power than an appropriate single radius. The precision of each image assumes that labels are shuffled at random, so when cells of a type cluster, an image can be weighted more than it deserves, which can cost power. Because spicyR v2 uses this precision only to weight images, and judges significance from the variation between patients through a cluster-robust variance [48, 49], a wrong precision does not invalidate the test. Finally, like every method, spicyR v2 inherits the segmentation and cell type annotation of each dataset, whose errors may systematically differ between groups. Errors of this kind, such as those between co-horts processed in different laboratories, would appear as changes in co-localisation. Therefore, segmentation and annotation should be done in the same way for every group. In our analyses, however, one of the more notable limitations in finding changes in co-localisation with spicyR was not its formulation but the size of the cohorts it was applied to.

Most of the cohorts that compared patients by clinical outcome, such as response to treatment or survival, were too small to detect a 10% change in co-localisation (a difference of 0.10 in the extra fraction), especially at the strict thresholds needed when many pairs of cell types are tested. A spicyR v2 model fitted to a pilot or published cohort can give the number of patients a new study needs, complementing simulation-based power analysis [68]. We recommend that future studies be sized in this way, ideally from a large published cohort, or from a small pilot allowing for the uncertainty in its estimate. Detecting such changes will need hundreds of patients, and cohorts of that size will come from combining studies across laboratories [42] and from imaging whole slides rather than selected regions, or predicting cell types across whole slides from histology [43]. Multi-cohort, whole-slide studies will hold hundreds of millions to billions of cells from tissue sectioned and imaged differently in each laboratory. A test of co-localisation for such studies must not mistake these differences for changes in co-localisation, and must be fast enough for this many cells. As spatial omics moves from single cohorts to multi-cohort, whole-slide studies, spicyR v2 provides a test of co-localisation that is robust to variation between tissues and fast enough for studies of this size, so that a change found in one cohort can be tested in the next.

## Data availability

Publicly available data were used for all evaluations. All data were downloaded as described in the originating manuscripts.

## Code availability

The *spicyR* R package is available from Bioconductor at https://www.bioconductor.org/packages/release/bioc/html/spicyR.html under the GNU General Public License (version 2 or later), and the Python package *spicyr* at https://github.com/SydneyBioX/spicyrpy. The code for every analysis and figure will be available at https://github.com/SydneyBioX/spicyR_v2_paper.

## Acknowledgments

The authors thank all their colleagues, particularly those at the Sydney Precision Data Science Centre and the Westmead Institute for Medical Research for their support and intellectual engagement. This work was supported by Australian Research Council Discovery Project grants to EP (DP260102836) and JTO (DP210100521). The funders had no role in the study design, the collection, analysis and interpretation of the data, the writing of the manuscript or the decision to submit it for publication. This research was undertaken with the assistance of resources and services from the National Computational Infrastructure (NCI), which is supported by the Australian Government, accessed through the Sydney Informatics Hub HPC Allocation Scheme, which is supported by the Deputy Vice-Chancellor (Research), University of Sydney.

## Author contributions

SD and EP conceived the study. SD developed the method and performed the analyses, with contributions from JTO, EW, FA and SR. EP supervised the study. All authors contributed to writing and reviewing the manuscript and approved the final version.

