## Supplementary Information for "spicyR enables robust, well-powered co-localisation tests in large spatial omics cohorts"

**This file contains:**

- Supplementary Figures S1–S6
- Supplementary Tables S1–S2
- Supplementary Sections 1–7: the method in full, with derivations and proofs
- Notation
- Supplementary references

### Contents

**Supplementary Figures** 4

**Supplementary Tables** 11

**1 Overview** 18

**2 Data** 19

**3 The effect in one image** 20

**7 Algorithm and cost** **83**

**8 Notation** **86**

### Supplementary Figures

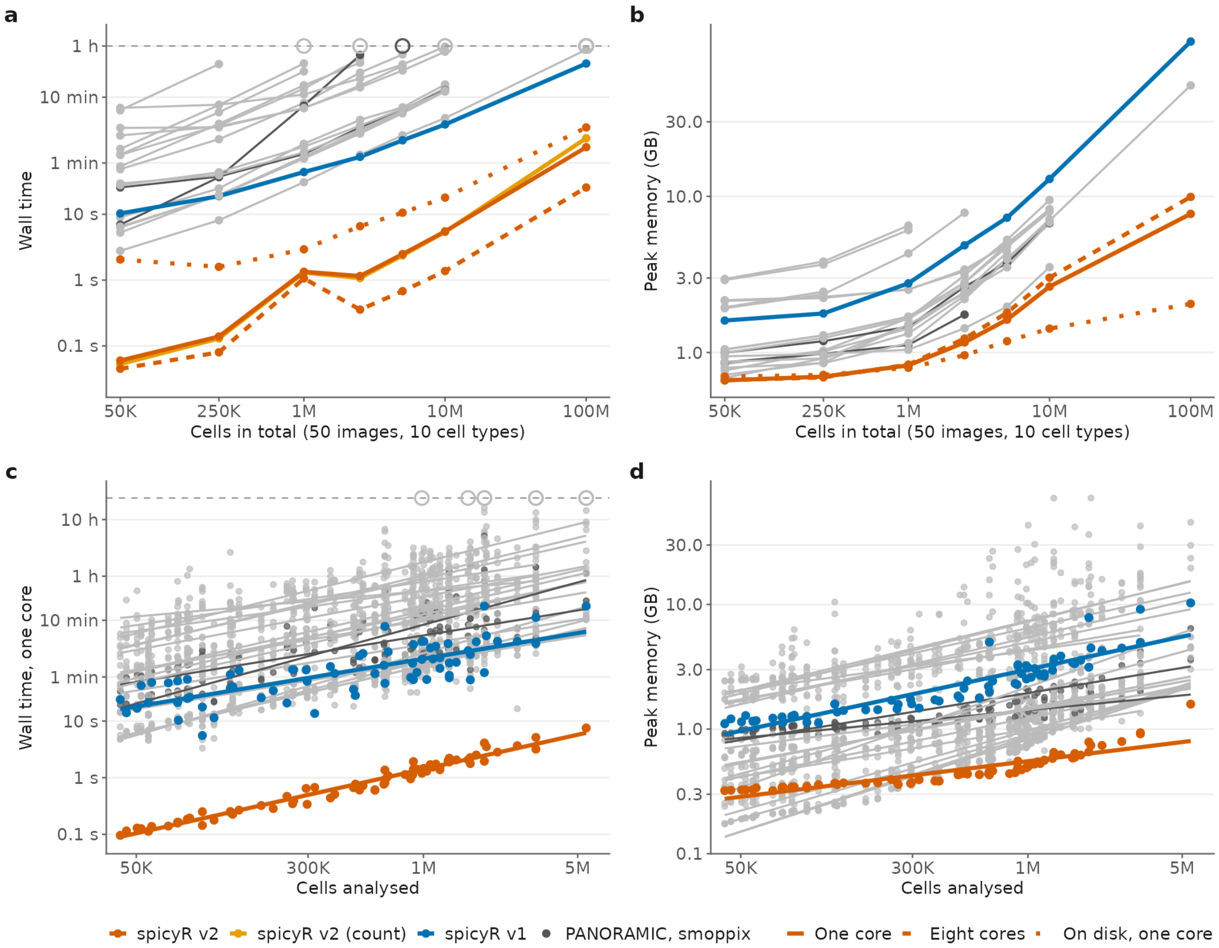

Supplementary Fig. S1: **spicyR v2 is faster and uses less memory than other methods.** Each point is one run of a method; orange, spicyR v2; amber, spicyR v2 with the count; blue, spicyR v1; dark grey, PANORAMIC and smoppix; light grey, other methods. (a) Run time for synthetic studies of 50 images and 10 cell types, testing every ordered pair, with a one-hour limit in place (grey dashed line; open circles, runs stopped at the limit). Other methods ran on one core, and spicyR v2 on one core (solid line), on eight cores (orange dashed) and on one core reading the cells from a file on disk rather than holding them in memory (dotted). (b) Peak memory of the same runs as in panel (a), counting everything held in memory (the cells, the method's calculations and R or Python itself); runs stopped at the time limit are not shown. (c) Run time on one core for 74 real analyses of 42 thousand to 5.5 million cells; each point is one analysis, and each line is a log-log fit; open circles, runs stopped at the 24-hour limit (grey dashed line). The real analyses ran on shared computers alongside other jobs, so these times are approximate. (d) Peak memory of the same analyses as in panel (c), counting everything held in memory (the cells, the method's calculations and R or Python itself), with log-log fits; runs stopped at the time limit are not shown.

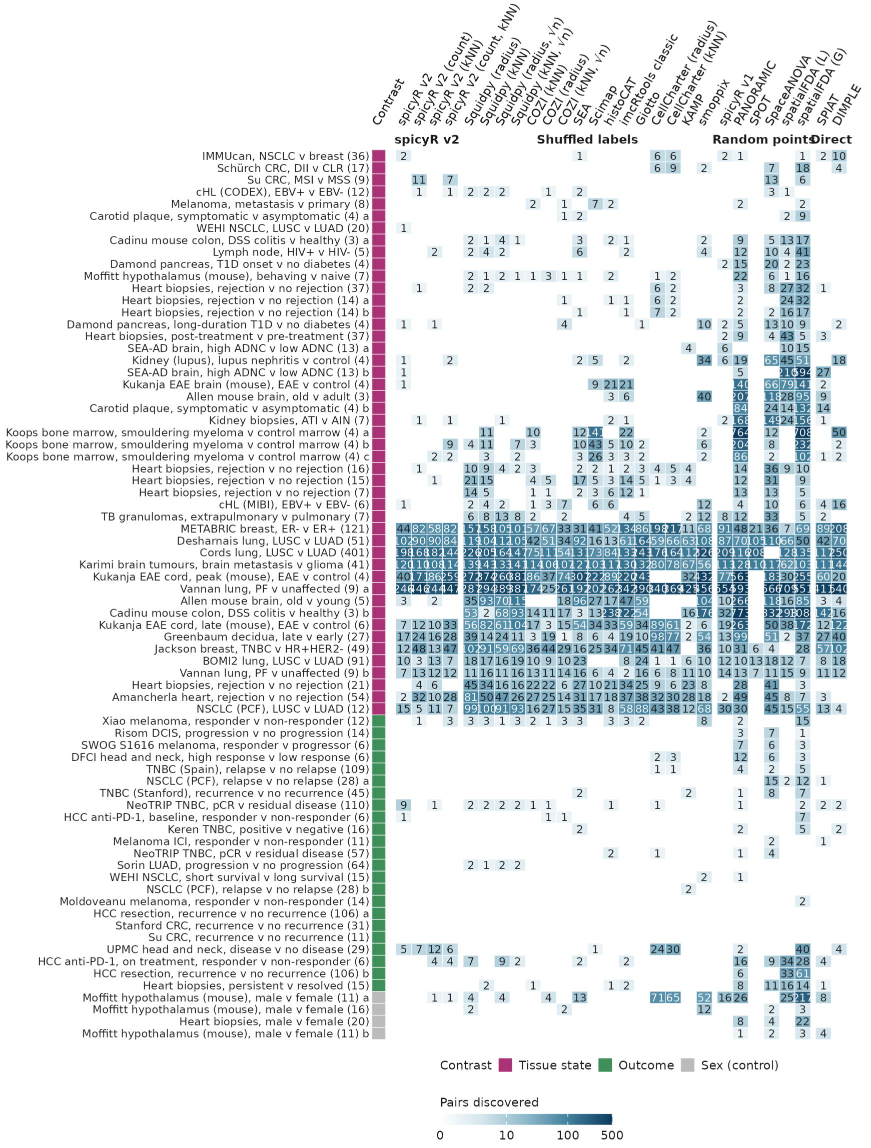

Supplementary Fig. S2: **Discoveries by method and analysis.** Number of pairs discovered (BH-adjusted $p < 0.05$) by each of 28 methods in each of 74 analyses, ordered as in Fig. 3a; number of patients in the smaller group in brackets; letters distinguish analyses of the same cohort at different cell type resolutions or subsets; dash, no result.

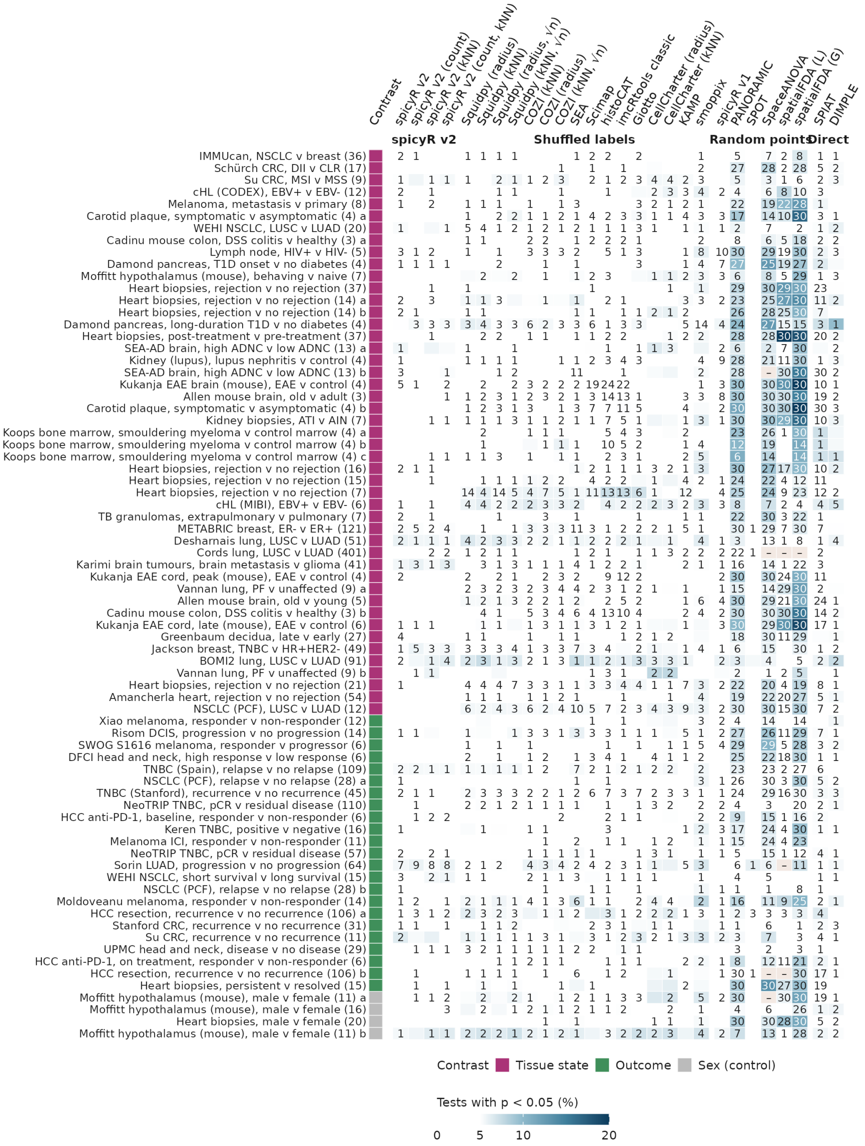

Supplementary Fig. S3: **Calibration by method and analysis under shuffled labels.** Fill, percentage of pair tests with $p < 0.05$ over 30 shuffles of patients between groups (white up to 5%); number, shuffles with any discovery; dash, fewer than 30 shuffles or no result. Rows as in Supplementary Fig. S2.

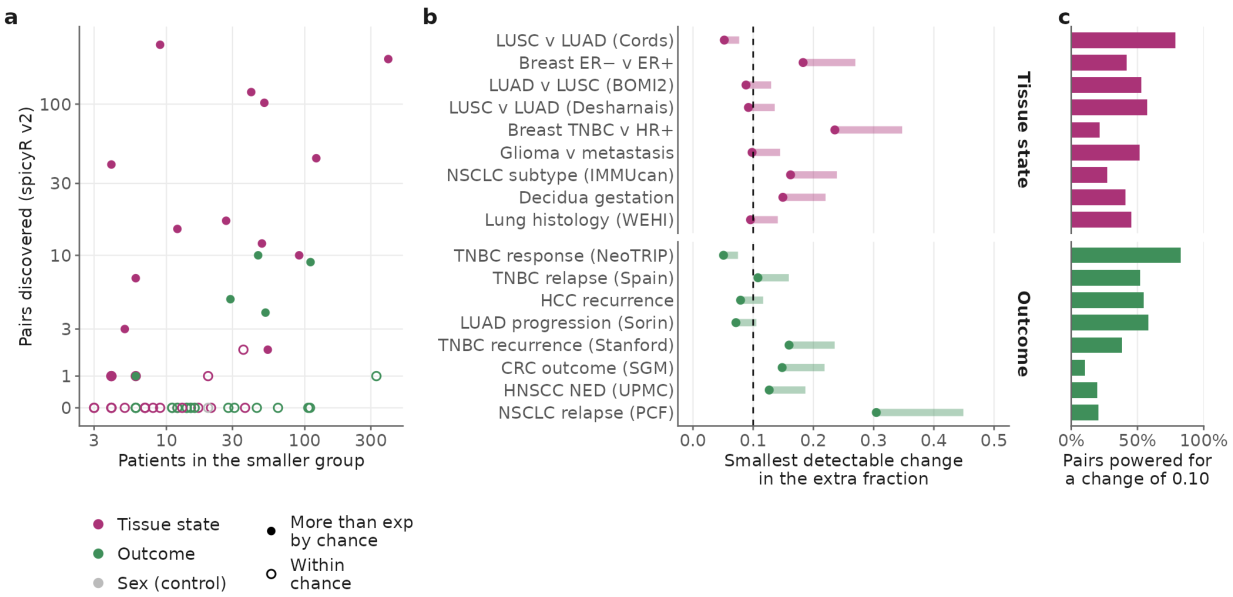

Supplementary Fig. S4: **Power of the real cohorts.** (a) Number of pairs discovered by spicyR v2 with the real groups (BH-adjusted $p < 0.05$) against the number of patients in the smaller group (for survival outcomes, the smaller of the numbers of patients with and without an event); filled, more pairs with $p < 0.05$ than in every shuffle. (b) Smallest change in the extra fraction that each cohort could detect with 80% power with the patients it has, for cohorts with at least 20 patients per group. For each cohort, this change was computed for every pair, and the median over pairs is shown. Dot, significance level 0.05; bar end, 0.001. Cohorts to the right of the dashed line (0.10) could not detect a 10% change. (c) For each cohort, the percentage of its pairs that could detect a change of 0.10 with 80% power at a significance level of 0.05.

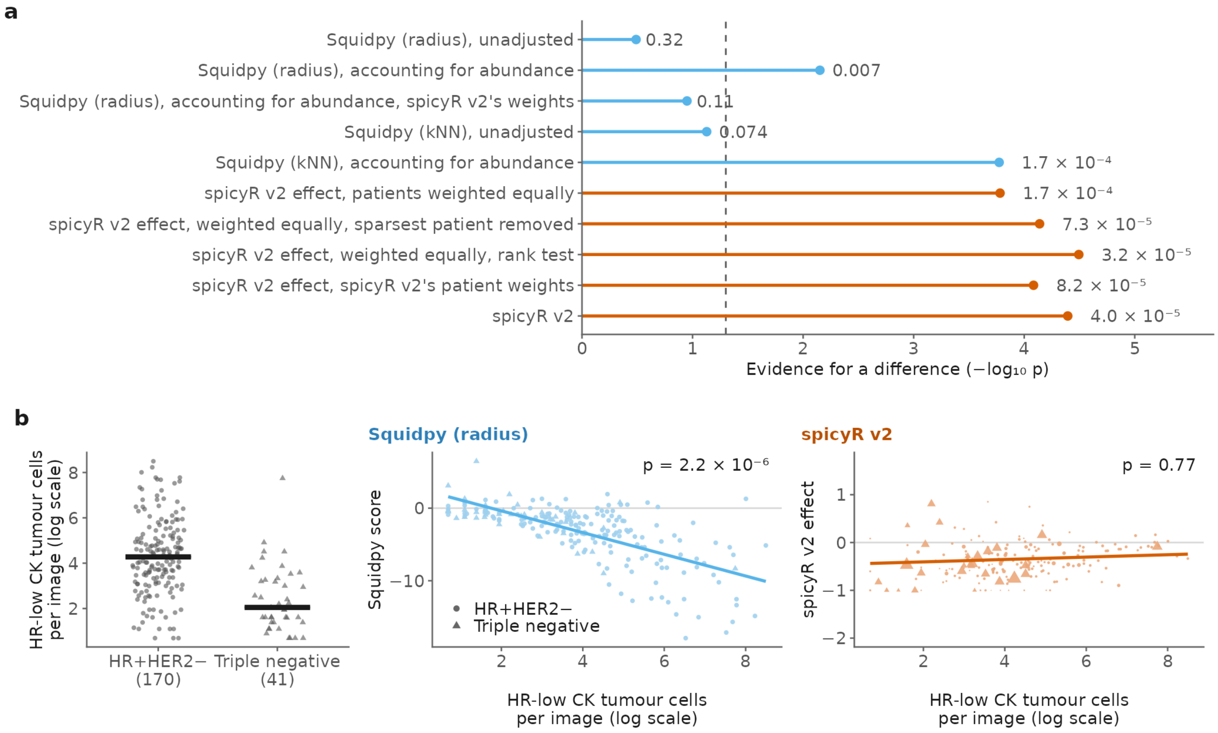

Supplementary Fig. S5: **Weighting and abundance in the Jackson example.** (a) p-values for the metacluster 24 tumour to small elongated stromal pair under different tests. Squidpy (radius and kNN): its patient scores compared between subtypes without and with adjustment for abundance, and, for the radius score, also weighted by spicyR v2's patient weights. spicyR v2: its patient effects compared with every patient weighted equally (also without the sparsest patient, and by a Wilcoxon rank-sum test), by a weighted linear model with spicyR v2's patient weights, and by the full spicyR v2 model. Dashed line, $p = 0.05$. (b) The pair in Fig. 5f (bottom): HR-low CK tumour cells per image in each subtype (left; patient means, log scale; bars, medians), and Squidpy (radius)'s score (middle) and spicyR v2's effect (right) against that abundance, with the $p$-value for the difference between subtypes.

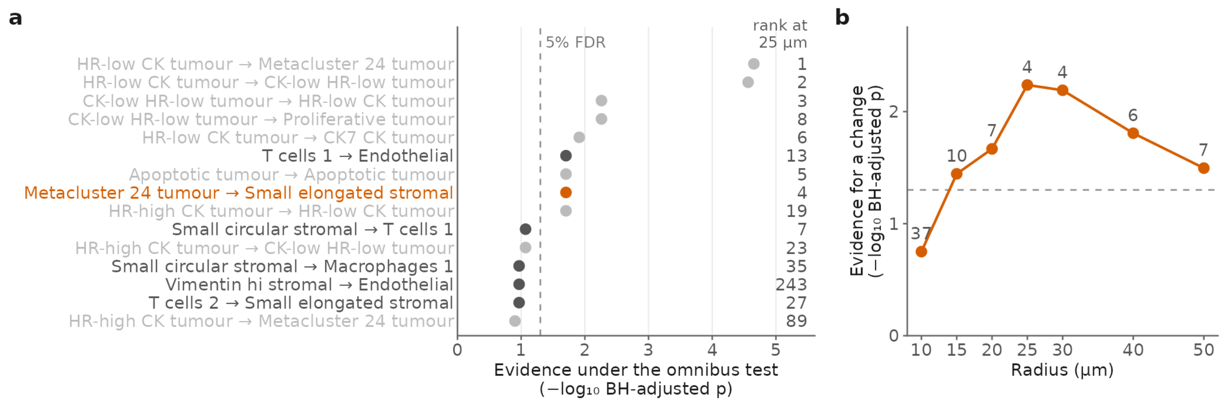

Supplementary Fig. S6: **The breast cancer pair of Fig. 5 is a discovery at every radius from 15 to 50 $\mu$m.** (a) The 15 pairs with the most evidence for a difference between triple-negative and HR+HER2- tumours under spicyR v2's omnibus test, which combines radii from 2 to 50 $\mu$m (BH-adjusted within the 576 tested pairs). On the right, each pair's rank among the 576 pairs when tested at 25 $\mu$m alone. Pairs of two tumour populations in grey, the metacluster 24 tumour to small elongated stromal pair in orange and other pairs in dark grey; dashed line, 5% false discovery rate. (b) Evidence for a difference for the metacluster 24 tumour to small elongated stromal pair, tested at one radius at a time from 10 to 50 $\mu$m; the number above each point is the pair's rank among the 576 pairs at that radius; dashed line, 5% false discovery rate. At 5 $\mu$m, too few stromal cells had a metacluster 24 cell nearby to test the pair.

#### Supplementary Tables

Table S1: **The 74 two-group analyses and three survival analyses of real data.** One row per analysis. Type: T, tissue state; O, outcome; S, sex (negative control); V, survival. Units: the patients (or, where marked, B, biopsies; D, donors; G, clinical groups; M, mice) in the reference and comparison groups, the units of inference; for survival, the patients with a follow-up time and event status, and the number of events (E: deaths, or recurrences and deaths for disease-free survival). Cell types: cell types tested, those with at least five cells in at least three images of each group (for survival, of the cohort); all cells were kept as neighbours. $r$: radius in $\mu$m, or in pixels (px) chosen to be about 25 $\mu$m. Accession: the deposit of the data analysed.

| Reference | Technology | Tissue | Contrast (reference v comparison) | Type | Units | Images | Cells | Cell types | $r$ | Accession |
|---|---|---|---|---|---|---|---|---|---|---|
| Koops et al. [1] | Xenium | Bone marrow, myeloma | control bone marrow v smouldering myeloma | T | 5 v 4 | 13 | 1,309,534 | 17 (broad) | 25 | Mendeley 10.17632/8bcwmrxp6w.3 |
| Koops et al. [1] | Xenium | Bone marrow, myeloma | control bone marrow v smouldering myeloma | T | 5 v 4 | 13 | 1,309,534 | 12 (major) | 25 | Mendeley 10.17632/8bcwmrxp6w.3 |
| Koops et al. [1] | Xenium | Bone marrow, myeloma | control bone marrow v smouldering myeloma | T | 5 v 4 | 13 | 1,309,534 | 37 (refined) | 25 | Mendeley 10.17632/8bcwmrxp6w.3 |
| Gabitto et al. [2] | MERFISH | Brain, Alzheimer disease | no or low v intermediate or high AD neuropathological change | T | 13 v 14 D | 69 | 1,887,729 | 24 (subclass) | 25 | SEA-AD (sea-ad-spatial-transcriptomics) |
| Gabitto et al. [2] | MERFISH | Brain, Alzheimer disease | no or low v intermediate or high AD neuropathological change | T | 13 v 14 D | 69 | 1,887,729 | 137 (supertype) | 25 | SEA-AD (sea-ad-spatial-transcriptomics) |
| Eling et al. [3] | IMC | Breast and lung cancer | breast cancer v NSCLC | T | 44 v 36 | 80 | 208,739 | 15 | 25 | Zenodo 12912567 |
| Jackson et al. [4] | IMC | Breast cancer (Basel) | HR+HER2- v triple-negative | T | 176 v 49 | 306 | 692,125 | 24 | 25 | Zenodo 3518284 |
| Danenberg et al. [5] | IMC | Breast cancer (METABRIC) | ER-positive v ER-negative | T | 481 v 121 | 673 | 983,167 | 32 | 25 | Zenodo 5850952 |
| Risom et al. [6] | MIBI-TOF | Breast DCIS | DCIS non-progressor v progressor (later invasive recurrence) | O | 44 v 14 | 58 | 50,413 | 21 | 51 px | Mendeley d87vg86zd8 |
| Pauli et al. [7] | Xenium | Carotid plaque | asymptomatic v symptomatic plaque | T | 8 v 4 | 12 | 99,625 | 12 (low) | 25 | Zenodo 17520216 |
| Pauli et al. [7] | Xenium | Carotid plaque | asymptomatic v symptomatic plaque | T | 8 v 4 | 12 | 99,625 | 46 (substates) | 25 | Zenodo 17520216 |
| Schürch et al. [8] | CODEX | Colorectal cancer | Crohn-like reaction v diffuse inflammatory infiltration | T | 17 v 18 | 140 | 258,385 | 23 | 65 px | Mendeley 10.17632/mpjzbt-fgfr.1 |
| Wu et al. [9] | CODEX | Colorectal cancer | no recurrence v recurrence | O | 71 v 31 | 186 | 396,127 | 15 | 25 | Zenodo 13179600 |
| Su et al. [10] | IMC | Colorectal cancer | no recurrence v recurrence | O | 11 v 38 | 154 | 825,352 | 10 | 25 | Zenodo 13901180 |
| Su et al. [10] | IMC | Colorectal cancer | MSS v MSI | T | 33 v 9 | 132 | 680,555 | 10 | 25 | Zenodo 13901180 |
| Greenbaum et al. [11] | MIBI-TOF | Decidua | early (6-12 wk) v late (14-20 wk) gestation | T | 27 v 39 D | 211 | 293,956 | 24 | 25 | figshare 10.6084/m9.figshare.16663465 |
| Karimi et al. [12] | IMC | Glioma, brain metastasis | glioma v brain metastasis | T | 139 v 41 | 327 | 984,147 | 17 | 25 | Zenodo 7884599 |
| Wu et al. [9] | CODEX | Head and neck cancer | low v high pathological tumour response | O | 6 v 12 G | 58 | 136,680 | 20 | 25 | Zenodo 13179600 |

*Supplementary Table S1, continued*

| Reference | Technology | Tissue | Contrast (reference v comparison) | Type | Units | Images | Cells | Cell types | $r$ | Accession |
|---|---|---|---|---|---|---|---|---|---|---|
| Wu et al. [9] | CODEX | Head and neck cancer | no evidence of disease v recurrence/death (NED v non-NED) | O | 52 v 29 | 308 | 2,165,215 | 17 | 25 | Zenodo 13179600 |
| Amancherla et al. [13] | Xenium | Heart transplant | no rejection v acute cellular rejection (biopsies) | T | 97 v 54 B | 151 | 133,344 | 20 | 25 | GEO GSE290577 |
| Amancherla et al. [13] | Xenium | Heart transplant | no rejection v cellular rejection (one random biopsy per patient) | T | 39 v 16 | 55 | 45,113 | 19 | 25 | GEO GSE290577 |
| Amancherla et al. [13] | Xenium | Heart transplant | no rejection v cellular rejection (biopsies after rejection treatment) | T | 87 v 21 B | 108 | 85,222 | 18 | 25 | GEO GSE290577 |
| Amancherla et al. [13] | Xenium | Heart transplant | no rejection v cellular rejection (after treatment; patients grouped by worst biopsy) | T | 44 v 15 | 108 | 85,222 | 18 | 25 | GEO GSE290577 |
| Amancherla et al. [13] | Xenium | Heart transplant | no rejection v cellular rejection (after treatment; concordant patients) | T | 44 v 7 | 92 | 69,870 | 18 | 25 | GEO GSE290577 |
| Amancherla et al. [13] | Xenium | Heart transplant | no rejection v cellular rejection grade $\geq 2$ (biopsies before treatment) | T | 37 v 44 B | 81 | 76,944 | 21 | 25 | GEO GSE290577 |
| Amancherla et al. [13] | Xenium | Heart transplant | no rejection v cellular rejection (before treatment; patients grouped by worst biopsy) | T | 14 v 32 | 81 | 76,944 | 17 | 25 | GEO GSE290577 |
| Amancherla et al. [13] | Xenium | Heart transplant | no rejection v cellular rejection (before treatment; concordant patients) | T | 14 v 18 | 53 | 52,629 | 17 | 25 | GEO GSE290577 |
| Amancherla et al. [13] | Xenium | Heart transplant | rejection resolved v persisted after treatment (pre-treatment biopsies) | O | 16 v 15 | 56 | 58,703 | 21 | 25 | GEO GSE290577 |
| Amancherla et al. [13] | Xenium | Heart transplant | female v male recipients (negative control) | S | 20 v 35 | 151 | 133,344 | 22 | 25 | GEO GSE290577 |
| Amancherla et al. [13] | Xenium | Heart transplant | no-rejection biopsies before v after treatment | T | 37 v 87 B | 124 | 87,298 | 21 | 25 | GEO GSE290577 |
| Qiu et al. [14] | CODEX | Hepatocellular carcinoma | no recurrence v recurrence | O | 106 v 172 | 288 | 3,231,437 | 71 | 65 px | Mendeley km2df7y256 |
| Qiu et al. [14] | CODEX | Hepatocellular carcinoma | no recurrence v recurrence | O | 106 v 172 | 288 | 3,231,437 | 9 (lineages) | 65 px | Mendeley km2df7y256 |
| Wu et al. [15] | PhenoCycler | Hepatocellular carcinoma | anti-PD-1 non-responder v responder (post-treatment resections) | O | 7 v 6 | 24 | 1,035,214 | 14 | 25 | Zenodo 19123188 |
| Wu et al. [15] | PhenoCycler | Hepatocellular carcinoma | anti-PD-1 non-responder v responder (pre-treatment samples) | O | 7 v 6 | 24 | 491,852 | 14 | 25 | Zenodo 19123188 |
| Yeo et al. [16] | PhenoCycler | Hodgkin lymphoma | EBV-negative v EBV-positive | T | 12 v 15 | 27 | 934,478 | 9 | 25 | Zenodo 18434741 |
| Yeo et al. [16] | MIBI-TOF | Hodgkin lymphoma | EBV-negative v EBV-positive | T | 14 v 6 | 42 | 1,409,210 | 13 | 25 | Zenodo 18434741 |
| Baker et al. [17] | Xenium 5K | Kidney | acute interstitial nephritis v acute tubular injury | T | 8 v 7 | 15 | 240,740 | 45 | 25 | Zenodo 21793755 |
| Danaher et al. [18] | CosMx | Kidney, lupus nephritis | control v lupus nephritis | T | 4 v 8 | 275 | 532,275 | 22 | 25 | Zenodo 13964258 |
| Sorin et al. [19] | IMC | Lung adenocarcinoma | no progression v progression | O | 340 v 64 | 404 | 1,593,404 | 17 | 25 | Zenodo 7760826 |
| Nordling et al. [20] | Vectra mIF | Lung cancer | LUAD v LUSC | T | 174 v 91 | 452 | 2,661,850 | 9 | 25 | Zenodo 17912877 |
| Cords et al. [21] | IMC | Lung cancer | LUAD v LUSC | T | 618 v 401 | 1,887 | 5,461,238 | 30 | 25 | Zenodo 7961844 |

*Supplementary Table S1, continued*

| Reference | Technology | Tissue | Contrast (reference v comparison) | Type | Units | Images | Cells | Cell types | $r$ | Accession |
|---|---|---|---|---|---|---|---|---|---|---|
| Desharnais et al. [22] | IMC | Lung cancer | LUAD v LUSC | T | 51 v 51 | 204 | 893,148 | 17 | 25 | Zenodo 14625562 |
| Marceaux et al. [23] | MIBI-TOF | Lung cancer | long v short overall survival (deposited groups) | O | 49 v 15 | 154 | 602,822 | 13 | 25 | Zenodo 17480094 |
| Marceaux et al. [23] | MIBI-TOF | Lung cancer | LUAD v LUSC | T | 44 v 20 | 154 | 602,822 | 13 | 25 | Zenodo 17480094 |
| Kulasinghe et al. [24] | PhenoCycler | Lung cancer | no relapse v relapse after adjuvant chemotherapy | O | 33 v 28 | 110 | 1,187,511 | 14 (tumour subtypes) | 25 | Zenodo 22749853 |
| Kulasinghe et al. [24] | PhenoCycler | Lung cancer | no relapse v relapse after adjuvant chemotherapy | O | 33 v 28 | 110 | 1,187,511 | 34 | 25 | Zenodo 22749853 |
| Kulasinghe et al. [24] | PhenoCycler | Lung cancer | LUAD v LUSC | T | 45 v 12 | 103 | 1,103,785 | 34 | 25 | Zenodo 22749853 |
| Vannan et al. [25] | Xenium | Lung fibrosis | control v pulmonary fibrosis | T | 9 v 26 | 45 | 1,630,319 | 40 | 25 | GEO GSE250346 |
| Vannan et al. [25] | Xenium | Lung fibrosis | control v pulmonary fibrosis | T | 9 v 26 | 45 | 1,630,319 | 4 (lineages) | 25 | GEO GSE250346 |
| Liu et al. [26] | MIBI-TOF | Lymph node, HIV | HIV-negative v HIV-positive | T | 5 v 5 D | 166 | 1,734,664 | 22 | 25 | BioImage Archive S-BIAD2864 |
| Moldoveanu et al. [27] | IMC | Melanoma | ICI non-responder v responder | O | 16 v 14 | 30 | 112,497 | 10 | 25 | Zenodo 5903179 |
| Dong et al. [28] | CosMx | Melanoma | non-responder (SD/PD) v responder (CR/PR) | O | 14 v 11 | 26 | 56,761 | 11 | 140 px | Zenodo 14708000 |
| Dong et al. [28] | CosMx | Melanoma | primary v metastasis (node or distant) | T | 8 v 17 | 26 | 56,761 | 11 | 140 px | Zenodo 14708000 |
| Campbell et al. [29] | MIBI-TOF | Melanoma | combination-arm progressor v responder (baseline) | O | 6 v 7 | 64 | 297,271 | 18 | 25 | Zenodo 15547911 |
| Xiao et al. [30] | IMC | Melanoma | anti-PD-1 non-responder v responder (pre-treatment) | O | 12 v 14 | 158 | 662,266 | 20 | 25 | Zenodo 6838169 |
| Allen et al. [31] | MERFISH | Mouse brain | adult (24 wk) v old (90 wk) mice | T | 3 v 5 M | 21 | 255,278 | 43 | 25 | CELLxGENE 31937775 |
| Allen et al. [31] | MERFISH | Mouse brain | young/adult (4 and 24 wk) v old (90 wk) mice | T | 7 v 5 M | 31 | 378,918 | 43 | 25 | CELLxGENE 31937775 |
| Kukanja et al. [32] | ISS | Mouse CNS, EAE | control v EAE mice (brain, peak) | T | 4 v 4 M | 8 | 321,543 | 41 | 25 | Zenodo 8037425 |
| Kukanja et al. [32] | ISS | Mouse CNS, EAE | control v EAE mice (spinal cord, late) | T | 6 v 6 M | 12 | 77,601 | 37 | 25 | Zenodo 8037425 |
| Kukanja et al. [32] | ISS | Mouse CNS, EAE | control v EAE mice (spinal cord, peak) | T | 4 v 4 M | 24 | 146,007 | 35 | 25 | Zenodo 8037425 |
| Cadinu et al. [33] | MERFISH | Mouse colon, colitis | healthy v DSS day 9 mice | T | 3 v 6 M | 14 | 900,427 | 7 (tier 1) | 25 | Dryad 10.5061/dryad.rjdfn2zh3 |
| Cadinu et al. [33] | MERFISH | Mouse colon, colitis | healthy v DSS day 9 mice | T | 3 v 6 M | 14 | 900,427 | 58 (tier 2) | 25 | Dryad 10.5061/dryad.rjdfn2zh3 |
| Moffitt et al. [34] | MERFISH | Mouse hypothalamus | female v male mice (all animals) | S | 16 v 20 M | 181 | 874,768 | 15 | 25 | Dryad 10.5061/dryad.8t8s248 |
| Moffitt et al. [34] | MERFISH | Mouse hypothalamus | naive v behaving male mice | T | 7 v 13 M | 100 | 482,199 | 15 | 25 | Dryad 10.5061/dryad.8t8s248 |
| Moffitt et al. [34] | MERFISH | Mouse hypothalamus | female v male mice (sex-matched behaviours) | S | 11 v 14 M | 137 | 668,741 | 81 (fine) | 25 | Dryad 10.5061/dryad.8t8s248 |
| Moffitt et al. [34] | MERFISH | Mouse hypothalamus | female v male mice (sex-matched behaviours) | S | 11 v 14 M | 137 | 668,741 | 15 | 25 | Dryad 10.5061/dryad.8t8s248 |

*Supplementary Table S1, continued*

| Reference | Technology | Tissue | Contrast (reference v comparison) | Type | Units | Images | Cells | Cell types | $r$ | Accession |
|---|---|---|---|---|---|---|---|---|---|---|
| Damond et al. [35] | IMC | Pancreas, type 1 diabetes | non-diabetic v long-duration T1D donors | T | 4 v 4 D | 555 | 1,137,088 | 13 | 25 | Mendeley 10.17632/cydmws-fztj.1 |
| Damond et al. [35] | IMC | Pancreas, type 1 diabetes | non-diabetic v recent-onset T1D donors | T | 4 v 4 D | 564 | 1,292,937 | 14 | 25 | Mendeley 10.17632/cydmws-fztj.1 |
| Keren et al. [36] | MIBI-TOF | TNBC | no recurrence v recurrence | O | 21 v 16 | 37 | 184,137 | 17 | 65 px | ExperimentHub EH8420 |
| Ranek et al. [37] | MIBI-TOF | TNBC (Spain) | no relapse v relapse | O | 205 v 109 | 548 | 1,928,329 | 31 | 65 px | Zenodo 14290163 |
| Ranek et al. [37] | MIBI-TOF | TNBC (Stanford) | no recurrence v recurrence | O | 97 v 45 | 277 | 1,073,456 | 31 | 65 px | Zenodo 14290163 |
| Wang et al. [38] | IMC | TNBC (NeoTRIP) | residual disease v pCR (both arms) | O | 133 v 110 | 766 | 1,003,614 | 21 | 25 | Zenodo 20414755 (deposited by Meyer et al.) |
| Wang et al. [38] | IMC | TNBC (NeoTRIP) | residual disease v pCR (chemotherapy + atezolizumab arm) | O | 67 v 57 | 394 | 513,383 | 21 | 25 | Zenodo 20414755 (deposited by Meyer et al.) |
| McCaffrey et al. [39] | MIBI-TOF | Tuberculosis granuloma | pulmonary v extrapulmonary granulomas | T | 8 v 7 | 30 | 42,310 | 14 | 25 | Mendeley 10.17632/dr5fkgtrb6 |
| Meyer et al. [40] | IMC | TNBC (discovery cohort) | overall survival | V | 208 (46 E) | 437 | 908,423 | 10 | 25 | Zenodo 20414755 |
| Meyer et al. [40] | IMC | TNBC (discovery cohort) | disease-free survival | V | 163 (52 E) | 348 | 720,875 | 10 | 25 | Zenodo 20414755 |
| Cords et al. [21] | IMC | Lung cancer | overall survival | V | 1,059 (730 E) | 1,960 | 5,640,390 | 30 | 25 | Zenodo 7961844 |

Table S2: **Methods compared.** Approach as grouped in the figures; the per-image statistic and its settings; how groups of patients were compared; and whether the method tests one direction of a pair (D) or is symmetric (S). Every method used a radius of 25 $\mu$m (or its pixel equivalent) or $k = 10$ nearest neighbours unless stated. For methods that give a score per image, the comparison of groups was our two-sample $t$-test (pooled variance) of patient means. Numbers of permutations for COZI, SEA, Scimap, histoCAT, imcRtools classic and Giotto follow the benchmark of Schiller et al. [41]; the packages' defaults are 1,000, and 100 for COZI.

| Method | Package (version) | Approach | Statistic and settings | Comparison of groups | Dir. |
|---|---|---|---|---|---|
| spicyR v2 (extra fraction, extra count; radius or $k$NN) | spicyR (1.99.56), R | spicyR v2 | Extra fraction or count of neighbours over random labelling of the observed cells, with its variance under random labelling in closed form (sampled for a self-pair's fraction) | Two-level model of images within patients (Paule–Mandel variances), group means compared by their CR2 variances with patients as clusters, Satterthwaite df | D |
| Squidpy (radius, $k$NN) [42] | squidpy (1.8.2), Python | Shuffled labels | Neighbourhood enrichment $z$-score, 1,000 permutations; also divided by $\sqrt{n}$ | Patient means, $t$-test | radius: S; $k$NN: D |
| COZI (radius, $k$NN) [41] | cozipy (0.1.4), Python | Shuffled labels | Conditional-ratio $z$-score, 300 permutations, $k = 10$ (authors: 5); also divided by $\sqrt{n}$ | Patient means, $t$-test | D |
| SEA [41] | scimap (2.2.11, fork of Schiller et al.), Python | Shuffled labels | Neighbourhood $z$-score normalised by total counts, 300 permutations | Patient means, $t$-test | D |
| Scimap [43] | scimap (2.2.11, fork of Schiller et al.), Python | Shuffled labels | Signed neighbour-frequency score, 300 permutations | Patient means, $t$-test | D |
| histoCAT [44] | imcRtools (1.19.1), R | Shuffled labels | Conditional interaction test, 200 permutations, significance indicator | Patient means, $t$-test | D |
| imcRtools classic [45] | imcRtools (1.19.1), R | Shuffled labels | Classic interaction test, 200 permutations, significance indicator | Patient means, $t$-test | D |
| Giotto [46] | Giotto (4.2.3), R | Shuffled labels | Cell proximity enrichment, 100 simulations | Patient means, $t$-test | S |
| CellCharter (radius, $k$NN) [47] | cellcharter (0.3.7), Python | Shuffled labels | Differential neighbourhood enrichment between groups | Group labels permuted between patients (reimplemented), at least 10,000 permutations, so that the smallest attainable $p$-value is about $2 \times 10^{-4}$ | D |
| KAMP [48] | Implemented from the authors' formulas, R | Shuffled labels | Cross-type $K$ at $r$ minus its exact random-labelling mean, convex-hull window; for pairs of a cell type with itself, the univariate random-labelling mean | Patient means, $t$-test | S |
| smoppix [49] | smoppix (1.5.4), R | Shuffled labels | Nearest-neighbour probabilistic index against the background of all cells | Weighted linear mixed model with a patient random effect | S |
| spicyR v1 [50] | spicyR (1.24.1), R | Random points | $L$-function at $r$ against complete spatial randomness, weighted | Weighted linear mixed model with a patient random intercept | S |
| PANORAMIC [51] | panoramic (0.99.4), R | Random points | Edge-corrected local enrichment at $r$, concave-hull window | Random-effects meta-analysis with a patient level | D |
| SPOT [52] | SPOT (0.1.0), R | Random points | Cross-type $L$ at 26 radii from 0 to 50 $\mu$m (SPOT's tutorial uses 100 radii from 0 to a quarter of the shortest image side) | Logistic regression per radius on patient averages, combined by the Cauchy combination test | S |

*Supplementary Table S2, continued*

| Method | Package (version) | Approach | Statistic and settings | Comparison of groups | Dir. |
|---|---|---|---|---|---|
| SpaceANOVA [53] | SpaceANOVA (0.1.0), R | Random points | Pair correlation function at 26 radii from 0 to 50 $\mu$m, 20 permutations for the hole adjustment (documentation example; default 50) | Functional ANOVA with patients as subjects | S |
| spatialFDA $L$ and $G$ [54] | spatialFDA (1.5.2), R | Random points | Cross-type $L$ (isotropic correction) or $G$ (reduced-sample correction) at 26 radii from 0 to 50 $\mu$m | Functional additive mixed model with a patient random effect | $L$: S; $G$: D |
| SPIAT [55] | SPIAT (1.15.0), R | Direct | Log normalised mixing score at $r$; images with a score of zero kept by adding half the pair's smallest positive score | Patient means, $t$-test | D |
| DIMPLE [56] | DIMPLE (1.0.0), R | Direct | Jensen–Shannon distance between kernel intensity surfaces (bandwidth 30 and pixel size 10, in coordinate units) | Linear model of patient medians | S |

### 1 Overview

spicyR v2 needs no permutations because each image's score has an exact mean and variance under random labelling. This supplement derives those formulas and builds the rest of the method on them, from the extra fraction and the extra count in a single image to the model that combines a patient's images and the test that compares patients between groups. Along the way it shows what each score responds to, so that users can choose between them, and extends the method to covariates, more than two groups, time-to-event outcomes and several radii. The last section sets out the whole method as one algorithm, with its cost.

**The question.** We have spatial omics images of tissue from patients in two groups, and in every image each cell has a location and a cell type. For two cell types $A$ and $B$, we ask whether $B$ cells lie near $A$ cells more, or less, in one group than in the other. Covariates, more than two groups and time-to-event outcomes are treated in Section 6.

**What is measured.** In each image, what is observed is compared with *random labelling*: the cells stay where they are, and the $B$ labels are shuffled at random among the cells that are not $A$. Two effects are defined on this basis (Section 3).

- The *extra fraction*, the default. It compares the share of $B$ cells with an $A$ neighbour with the share expected under random labelling. Positive values mean *attraction* ($B$ cells have an $A$ neighbour more often than expected), negative values mean *avoidance* (less often), and 0 means the share is exactly as expected. The difference is divided by its *room*, the largest difference possible, reached if every $B$ cell had an $A$ neighbour (for attraction) or if none did (for avoidance).
- The *extra count*, an option. It compares the average number of $A$ neighbours of a $B$ cell with the number expected under random labelling. It is the difference between the two, in $A$ cells per $B$ cell, and has no fixed upper limit. Positive values mean attraction, negative values mean avoidance, and 0 means the number is exactly as expected.

The mean and variance of both under random labelling are exact, so no permutations are needed. The one exception is a term of the variance of a self-pair's extra fraction with a radius, which is by default estimated from a sample of cells (Remark 2).

**The model.** Each image contributes one effect, and the group means are estimated with a generalised estimating equation (GEE). Its working covariance comes from a two-level model that combines the images of each patient, and the patients of each group, as a random-effects meta-analysis combines studies. Each image's random-labelling variance plays the role of a study's known variance, and the model adds a variance between images of the same patient and a variance between patients, so that images and patients are weighted by their precision. The two-level model sets only these weights. The two means are compared with a CR2 sandwich variance clustered by patient, on Satterthwaite degrees of freedom, which does not rely on the model being right.

**The method in four steps.** Figure S7 shows the four steps and where each is described.

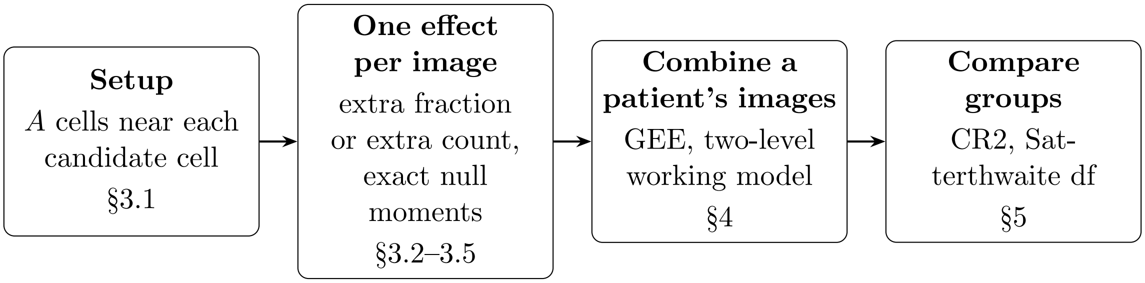

Supplementary Fig. S7: The method, from cells to a test between groups.

**Outline.** Section 2 sets out the patients, images and pairs, and Sections 3–5 follow the four steps of Figure S7. Section 6 gives the extensions, Section 7 the whole algorithm and its cost, and Section 8 the notation.

### 2 Data

**Patients and groups.** We write *patient* for the independent unit of a study, because the word is unambiguous, but the unit can be any subject from which the images come, such as an animal in a mouse study. There are $m$ patients, $i = 1, \ldots, m$. Each patient is in one of two groups, $c(i) \in \{1, 2\}$, with $m_c$ patients in group $c$ and $m = m_1 + m_2$. Group 1 is the reference, and every contrast is group 2 minus group 1. Covariates and more than two groups are treated in Sections 6.1 and 6.2.

**Images.** Patient $i$ has $J_i \geq 1$ images, $j = 1, \ldots, J_i$. Image $(i, j)$ is a set of $n_{ij}$ cells, each with a location and one cell-type label. We assume that images from different patients are independent. Images from the same patient may be dependent in any way. An image whose group is missing is removed before anything else.

**Patients are the units.** A difference between groups is a claim about patients, which should hold for new patients from the same groups. A patient's images share that patient's biology, and the cells of an image share their tissue, so treating images or cells as independent would overstate the sample size and give false positives. Every variance below is therefore clustered by patient, and every degree of freedom counts patients. The patient of every image is assumed known.

**Pairs.** A *pair* is an ordered pair of cell types, written $A \to B$. $A$ is the *anchor* type, whose cells are held fixed, and $B$ is the *tested* type, whose cells are the centres and whose labels are shuffled. For each $B$ cell we look at the $A$ cells near it. So $A \to B$ and $B \to A$ are different questions (Section 3.6). The pair $A \to A$ is a *self-pair* (Section 3.8). In image $(i, j)$, $n^A_{ij}$ and $n^B_{ij}$ are the numbers of $A$ and $B$ cells.

**Neighbours.** Two cells are neighbours if they are within distance $r$ of each other (the radius graph). Alternatively, each cell's neighbours are its $k$ nearest cells (the $k$-nearest-neighbour graph); this graph is not symmetric. Results are stated for the radius graph, with the $k$-nearest-neighbour version noted where it differs.

### 3 The effect in one image

#### 3.1 Setup

This section works with one image and one pair $A \to B$ with $A \neq B$. Within one image we drop the subscript $(i, j)$: the image has $n$ cells, $n^A$ of type $A$ and $n^B$ of type $B$.

**Candidates.** The *candidates* are the cells that are not of type $A$. There are $M = n - n^A$ of them, indexed by $\ell = 1, \ldots, M$. They are the cells that could carry a $B$ label: an $A$ cell cannot, because the $A$ labels are held fixed and each cell has one label. Cells of types other than $A$ and $B$ are candidates too. The $B$ cells are $n^B$ of the $M$ candidates, and $p = n^B/M$ is the share of candidates that are $B$.

**The number of each candidate.** Each candidate $\ell$ is given one number $x_\ell$, which depends on the effect:

- for the extra fraction, $x_\ell = 1$ if at least one $A$ cell is within $r$ of $\ell$, and $x_\ell = 0$ otherwise;
- for the extra count, $x_\ell$ is the number of $A$ cells within $r$ of $\ell$.

A candidate with three $A$ cells within $r$ has $x_\ell = 1$ for the extra fraction and $x_\ell = 3$ for the extra count. With the $k$-nearest-neighbour graph, "within $r$ of $\ell$" becomes "among the $k$ nearest cells to $\ell$".

**The observed total.** The observed total is the sum over the $B$ cells,

$$O \coloneqq \sum_{\ell \in B} x_\ell.$$

For the extra fraction, $O$ is the number of $B$ cells with an $A$ neighbour. For the extra count, $O$ is the total number of $A$ neighbours of the $B$ cells.

#### 3.2 The random-labelling null

**Definition.** Under *random labelling*, the positions of all cells and the $A$ labels are fixed, and the $n^B$ $B$ labels are placed on $n^B$ of the $M$ candidates, with every set of $n^B$ candidates equally likely. Equivalently, the $B$ labels are shuffled among the candidates.

**Why this null.** Random labelling takes the tissue as observed: where the cells are, how dense they are, and where the $A$ cells sit. It asks only whether the $B$ cells sit near $A$ cells more or less often than a random choice of $n^B$ candidates would. A null of complete spatial randomness instead places cells in empty space, and so needs a window, the tissue area and an edge correction. Random labelling needs none of these.

Write $\bar{x} := \frac{1}{M}\sum_{\ell=1}^{M} x_\ell$ and $\sigma_x^2 = \frac{1}{M}\sum_{\ell=1}^{M}(x_\ell - \bar{x})^2$ for the mean and variance of the $x_\ell$ over the candidates.

**Proposition 1.** *Let $x_1, \dots, x_M$ be fixed numbers, and let $O := \sum_{\ell \in B} x_\ell$, where $B$ is a set of $n^B$ of the $M$ candidates chosen at random, with every such set equally likely. Then*

$$\mathbb{E}(O) = n^B \bar{x}, \qquad \mathrm{Var}(O) = \frac{n^B(M - n^B)}{M - 1}\,\sigma_x^2.$$

*Proof.* This is the standard result for a total under simple random sampling without replacement [57, Theorems 2.1 and 2.2, pp. 22–24].

For each candidate $\ell$, let $I_\ell = 1$ if $\ell$ is in $B$ and $I_\ell = 0$ otherwise. Then

$$O = \sum_\ell x_\ell I_\ell \qquad \text{and} \qquad \sum_\ell I_\ell = n^B.$$

Since $n^B$ of the $M$ candidates are in $B$ and every candidate is equally likely to be one of them,

$$\Pr(I_\ell = 1) = \frac{n^B}{M}. \tag{1}$$

Now take two distinct candidates $\ell \neq \ell'$. Given that $\ell$ is in $B$, the remaining $n^B - 1$ members of $B$ are among the other $M - 1$ candidates, so $\ell'$ is also in $B$ with probability $(n^B - 1)/(M - 1)$. Hence

$$\Pr(I_\ell = 1 \text{ and } I_{\ell'} = 1) = \frac{n^B}{M} \cdot \frac{n^B - 1}{M - 1} = \frac{n^B(n^B - 1)}{M(M - 1)}. \tag{2}$$

By (1), $\mathbb{E}(I_\ell) = n^B/M$, so

$$\mathbb{E}(O) = \sum_\ell x_\ell \frac{n^B}{M} = n^B \bar{x}.$$

Subtract the mean from each $x_\ell$: let $d_\ell := x_\ell - \bar{x}$. By the definitions of $\bar{x}$ and $\sigma_x^2$,

$$\sum_\ell d_\ell = 0 \qquad \text{and} \qquad \sum_\ell d_\ell^2 = M\sigma_x^2.$$

Because $\sum_\ell I_\ell = n^B$,

$$\sum_\ell d_\ell I_\ell = \sum_\ell x_\ell I_\ell - \bar{x}\sum_\ell I_\ell = O - n^B \bar{x}.$$

By definition, $\mathrm{Var}(O) = \mathbb{E}\big[\{O - \mathbb{E}(O)\}^2\big]$, and so

$$\mathrm{Var}(O) = \mathbb{E}\Big[\Big(\sum_\ell d_\ell I_\ell\Big)^2\Big].$$

Expanding the square gives one term for each candidate and one for each ordered pair of distinct candidates:

$$\Big(\sum_\ell d_\ell I_\ell\Big)^2 = \sum_\ell d_\ell^2 I_\ell^2 + \sum_{\ell\neq\ell'} d_\ell d_{\ell'} I_\ell I_{\ell'}.$$

Since $I_\ell$ is 0 or 1, $I_\ell^2 = I_\ell$, whose mean is $n^B/M$ by (1). The product $I_\ell I_{\ell'}$ is 1 exactly when both candidates are in $B$, so its mean is given by (2). Taking expectations term by term,

$$\mathrm{Var}(O) = \frac{n^B}{M}\sum_\ell d_\ell^2 + \frac{n^B(n^B-1)}{M(M-1)}\sum_{\ell\neq\ell'} d_\ell d_{\ell'}.$$

Since $\sum_\ell d_\ell = 0$,

$$0 = \Big(\sum_\ell d_\ell\Big)^2 = \sum_\ell d_\ell^2 + \sum_{\ell\neq\ell'} d_\ell d_{\ell'}, \qquad \text{so} \qquad \sum_{\ell\neq\ell'} d_\ell d_{\ell'} = -\sum_\ell d_\ell^2 = -M\sigma_x^2.$$

Substituting $\sum_\ell d_\ell^2 = M\sigma_x^2$ and the sum over pairs,

$$\mathrm{Var}(O) = n^B\sigma_x^2 - \frac{n^B(n^B-1)}{M-1}\,\sigma_x^2 = n^B\sigma_x^2\Big(1-\frac{n^B-1}{M-1}\Big) = \frac{n^B(M-n^B)}{M-1}\,\sigma_x^2. \qquad \square$$

The factor $(M-n^B)/(M-1)$ is the finite-population correction: it is 1 when $n^B = 1$ and 0 when every candidate is $B$. When the $x_\ell$ are 0 or 1, as for the extra fraction, $O$ follows a hypergeometric distribution (Section 3.3).

**Corollary 1.1.** *Under random labelling, the mean and variance of the observed total $O$ of an image are*

$$E = n^B\bar{x}, \qquad V = \frac{n^B(M-n^B)}{M-1}\,\sigma_x^2,$$

*for both effects, and with either the radius graph or the $k$-nearest-neighbour graph.*

*Proof.* The $x_\ell$ are set by the $A$ cells, which random labelling does not move, so $O$ is a sum of fixed numbers over a random set of $n^B$ candidates and Proposition 1 applies. $\square$

Sections 3.3 and 3.4 choose $x_\ell$ and the scale of the effect. Some images cannot show any effect for a pair, and Section 3.5 says which, and why they are not used.

**Self-pairs.** For a self-pair $A \to A$ the $x_\ell$ are not fixed, because shuffling the labels also moves the anchor cells. Section 3.8 treats that case.

#### 3.3 Extra fraction

**The numbers.** For the extra fraction, $x_\ell = 1$ if candidate $\ell$ has at least one $A$ cell within $r$, and $x_\ell = 0$ otherwise. Then $O$ is the number of $B$ cells with an $A$ neighbour. The mean $\bar{x}$ is the share of candidates with an $A$ neighbour; we write it $q$.

**Corollary 1.2.** *For the extra fraction, $O$ follows a hypergeometric distribution under random labelling, with*

$$E = n^B q, \qquad V = \frac{n^B(M - n^B)}{M - 1}\, q(1 - q).$$

*Proof.* Of the $M$ candidates, $Mq$ have an $A$ neighbour. Under random labelling the $B$ cells are $n^B$ candidates drawn without replacement, and $O$ counts how many of them are among those $Mq$. This is the definition of a hypergeometric distribution. For the moments, apply Corollary 1.1 with $\bar{x} = q$. Since each $x_\ell$ is 0 or 1, $x_\ell^2 = x_\ell$, so $\sigma_x^2 = \frac{1}{M} \sum_\ell x_\ell^2 - \bar{x}^2 = q - q^2 = q(1 - q)$. □

**Two rooms.** Under random labelling, $n^B q$ of the $B$ cells are expected to have an $A$ neighbour and $n^B(1 - q)$ are not. There are two ways to measure the excess $O - E$ against them:

- the *gain room* $n^B(1 - q)$: the $B$ cells expected *without* an $A$ neighbour, which could gain one;
- the *loss room* $n^B q$: the $B$ cells expected *with* an $A$ neighbour, which could lose one.

For example, with $n^B = 100$ and $q = 0.3$, the gain room is 70. If 51 rather than 30 $B$ cells have an $A$ neighbour, the excess is 21, which is $21/70 = 0.3$ of the gain room.

**Choosing the scale.** One of the two rooms is used for every image of a pair at a given radius. It is the gain room if

$$\sum_{\text{images}} (O - E) \;\geq\; 0,$$

and the loss room otherwise, where the sum is over every image of every patient. The groups are not used. The scale is chosen afresh at each radius, so one pair can be on the gain scale at one radius and on the loss scale at another. There are two reasons for this.

1. *One scale per pair.* Every image of the pair is divided by the same kind of room, so the effects of different images, and of different groups, are on the same scale and can be combined and compared.
2. *No dependence on the groups.* The scale would be the same if the patients' groups were shuffled, so the scale is fixed before the groups are compared.

This is the only step in Section 3 that uses more than one image.

**The extra fraction.** Write $R$ for the chosen room. The extra fraction of an image is its excess as a share of the room,

$$Y := \frac{O - E}{R}.$$

Under random labelling, $Y$ has mean and variance

$$\mathbb{E}(Y) = 0, \qquad \operatorname{Var}(Y) = s^2 = \frac{V}{R^2},$$

since $\mathbb{E}(O) = E$ and $\mathrm{Var}(O) = V$, with the scale held fixed. Note that the room $R$ is chosen from the $O$ of every image, so it is not strictly fixed under random labelling, but for inference this is still valid because these moments are only used to weight the images: the variance of the test comes from CR2 (Section 5). In the example above, $Y = 21/70 = 0.3$.

**The scale.** Since $0 \le O \le n^B$ and $E = n^B q$, the effect always lies in

$$\begin{cases} -\dfrac{q}{1-q} \;\le\; Y \;\le\; 1 & \text{with the gain room,} \\[2ex] -1 \;\le\; Y \;\le\; \dfrac{1-q}{q} & \text{with the loss room.} \end{cases}$$

Each room is the excess at the extreme in its own direction, so in the pair's direction the effect ends at 1 in size. With the gain room, the most an image can show is every $B$ cell having an $A$ neighbour. The excess then equals the room, and $Y = 1$. With the loss room, the least is no $B$ cell having one. The excess is then minus the room, and $Y = -1$.

The other direction has no such end, because the room was not sized for it. Suppose the pair is on the gain scale, and an image has $n^B = 100$ and $q = 0.8$, so $E = 80$ and the gain room is 20. If no $B$ cell in this image has an $A$ neighbour, the excess is $-80$, four times the room, and $Y = -4 = -q/(1-q)$. Section 3.5 shows why images whose chosen room is tiny are not used.

#### 3.4 Extra count

**The numbers.** For the extra count, $x_\ell$ is the number of $A$ cells within $r$ of candidate $\ell$. Then $O$ is the total number of $A$ neighbours of the $B$ cells. The mean $\bar{x}$ is the average number of $A$ neighbours of a candidate; we write it $\lambda$.

**Corollary 1.3.** *For the extra count, under random labelling,*

$$E = n^B \lambda, \qquad V = \frac{n^B (M - n^B)}{M - 1}\, \sigma_x^2.$$

*Proof.* Apply Corollary 1.1 with $\bar{x} = \lambda$. □

Unlike the extra fraction, $O$ here has no named distribution: the $x_\ell$ can take any whole-number value, and the distribution of $O$ depends on all of them. Only its mean and variance are used.

**The extra count.** The extra count of an image is its excess per $B$ cell,

$$Y := \frac{O - E}{n^B} = \frac{O}{n^B} - \lambda.$$

It is the average number of $A$ neighbours of a $B$ cell, minus the number expected. It is measured in $A$ cells per $B$ cell. Under random labelling, $Y$ has mean and variance

$$\mathbb{E}(Y) = 0, \qquad \mathrm{Var}(Y) = s^2 = \frac{V}{(n^B)^2},$$

since $\mathbb{E}(O) = E$, $\mathrm{Var}(O) = V$, and $E$ and $n^B$ are fixed. For example, with $n^B = 100$ and $\lambda = 0.5$, $E = 50$. If the $B$ cells have 80 $A$ neighbours in total, the excess is 30 and $Y = 0.3$: each $B$ cell has, on average, 0.3 more $A$ neighbours than expected.

**The scale.** The extra count divides by $n^B$ in both directions, so, unlike the extra fraction, it needs no choice of scale. Since $O \geq 0$, the effect is at least $-\lambda$, reached when no $B$ cell has an $A$ neighbour. It has no fixed upper limit: the largest possible $Y$ depends on how many $A$ cells the most crowded candidates have.

#### 3.5 Uninformative images

Some images cannot show any effect for a pair at a radius. Such an image is not used for that pair at that radius. There are two kinds.

**No variance.** For a pair $A \to B$, by Proposition 1, $V = 0$ in exactly three cases:

1. the image has no $B$ cells ($n^B = 0$);
2. every candidate is $B$ ($n^B = M$);
3. all the $x_\ell$ are equal ($\sigma_x^2 = 0$).

In each case $O$ takes the same value under every labelling. If $n^B = M$, the only possible $B$ is the set of all candidates, so $O = \sum_\ell x_\ell = M\bar{x} = E$. If every $x_\ell$ equals some $c$, any $n^B$ candidates give $O = n^B c = E$. So $O - E = 0$ whatever the tissue looks like: the image cannot show attraction or avoidance, and its effect would be 0 by construction rather than from the data. Including it would add a zero that carries no information and pull the estimate towards 0.

*Remark* 1 (What the three cases mean). The method asks whether the $B$ cells sit near $A$ cells more or less often than the other candidates would. Each case removes one side of that comparison.

- *No $B$ cells.* There is nothing to measure.
- *Every candidate is $B$.* All cells other than the $A$ cells are $B$ cells, for example an image containing only tumour cells ($A$) and T cells ($B$). There are no other cells to compare the $B$ cells with: they sit exactly where the non-$A$ cells sit, because they are the non-$A$ cells.
- *All $x_\ell$ equal.* Usually this means that no candidate has an $A$ cell within $r$ ($A$ is absent from the image, or too sparse at this radius), or, for the extra fraction, that every candidate has one ($A$ is widespread, or $r$ is large). Either way, every possible placement of the $B$ cells has the same exposure to $A$, so the placement cannot reveal a preference.

An uninformative image is not evidence of no effect. Any effect it has cannot be seen at this radius. If $B$ cells cluster tightly around $A$ cells but every cell is within $r$ of some $A$ cell, the preference is invisible to the extra fraction at $r$ and may be visible at a smaller radius. This is why the check is made for each pair and each radius separately.

**Almost no room.** For the extra fraction, an image can have a variance above 0 and still be unable to show the pair's effect: when its chosen room $R$ is less than one cell. Fewer than one $B$ cell could then move in the pair's direction, while on the other side, where the effect has no end (Section 3.3), a single cell can make the effect very large. For example, suppose the pair is on the gain scale, and an image has $n^B = 200$ and $q = 0.9999$, so $E = 199.98$ and the gain room is $R = 0.02$ cells. If 199 $B$ cells have an $A$ neighbour, one fewer than all of them, the excess is $-0.98$ and $Y = -0.98/0.02 = -49$.

So an image with $R < 1$ is not used for that pair at that radius. This extends the first kind. On the gain scale $R = 0$ exactly when $q = 1$, and on the loss scale when $q = 0$, and both are case 3 above. Three points about the rule:

- *The scale comes first.* The scale is chosen from all images (Section 3.3), and only then are the images with $R < 1$ set aside. So the rule cannot change the scale.
- *It does not depend on the B labels.* $R$ is $n^B(1-q)$ or $n^B q$. Here $n^B$ is a count and $q$ is set by the $A$ cells, and neither changes when the $B$ labels are shuffled. So the rule is fixed before the $B$ labels are looked at.
- *It does not bound the effect.* An image with $R \geq 1$ can still have an effect outside $[-1, 1]$ on the other side, but no longer one that a single cell decides.

The extra count needs no such rule: it divides by $n^B$, a whole number of cells that is at least 1 in every image used.

#### 3.6 The relation between the extra fraction and the extra count

**Breadth and depth.** The extra fraction asks how many $B$ cells have an $A$ neighbour: the *breadth* of contact. A $B$ cell next to one $A$ cell and a $B$ cell next to ten count the same. The extra count asks how many $A$ neighbours the $B$ cells have in total, so it also reflects the *depth* of contact: a $B$ cell gaining its fifth $A$ neighbour adds to the extra count but not to the extra fraction. Proposition 2 makes this observation exact: it splits the extra count into a part due to breadth and a part due to depth.

**Notation.** Here both effects are used at once, so quantities are labelled by effect. Write $Y_F$ and $Y_C$ for the extra fraction and the extra count, and $O_F$ and $O_C$ for their observed totals:

- $O_F$ is the number of $B$ cells with at least one $A$ neighbour;
- $O_C$ is the total number of $A$ neighbours of the $B$ cells.

Write $c_\ell$ for the number of $A$ cells within $r$ of candidate $\ell$ (the extra count's $x_\ell$). Candidate $\ell$ has an $A$ neighbour exactly when $c_\ell \geq 1$, so the extra fraction's $x_\ell$ is 1 when $c_\ell \geq 1$ and 0 otherwise.

**Splitting an average into breadth and depth.** Take any group of cells and ask how many $A$ neighbours they have on average. Two things decide the answer: how many of the cells have any $A$ neighbour at all (breadth), and how many $A$ neighbours those cells have (depth). Cells with no $A$ neighbour add nothing to the total, but they still count in the number of cells.

To make this precise, suppose an image has $N$ cells, $K$ of them have at least one $A$ neighbour, and their $c_\ell$ add up to $T$. The cells with $c_\ell = 0$ add nothing to $T$, so $T$ is also the sum of $c_\ell$ over the $K$ cells with an $A$ neighbour. Hence

$$\underbrace{\frac{T}{N}}_{\text{average over all cells}} = \underbrace{\frac{K}{N}}_{\text{breadth}} \times \underbrace{\frac{T}{K}}_{\text{depth}}.$$

Here $T/K$ is the average number of $A$ neighbours of a cell *that has at least one.*

We use this split for two groups.

- *All candidates.* Here $N = M$, $K = Mq$ and $T = M\lambda$. Breadth is $q$, and depth is

  $$\bar{d} := \frac{M\lambda}{Mq} = \frac{\lambda}{q},$$

  the typical number of $A$ neighbours of a candidate that has at least one. The split reads $\lambda = q\,\bar{d}$.

- *The $B$ cells.* Here $N = n^B$, $K = O_F$ and $T = O_C$. Breadth is $O_F/n^B$, and depth is

  $$d_B := \frac{O_C}{O_F},$$

  the number of $A$ neighbours of a $B$ cell that has at least one. The split reads $O_C/n^B = (O_F/n^B)\,d_B$.

For example, take an image with $M = 10$ candidates whose numbers of $A$ neighbours are

| candidate $\ell$ | 1 | 2 | 3 | 4 | 5–10 |
|---|---|---|---|---|---|
| $c_\ell$ | 1 | 2 | 3 | 2 | 0 |

and let the $B$ cells be candidates 2, 3 and 5, so $n^B = 3$. Then

| group | average over all cells | breadth | depth | check |
|---|---|---|---|---|
| all candidates | $\lambda = 8/10 = 0.8$ | $q = 4/10 = 0.4$ | $\bar{d} = 8/4 = 2$ | $0.4 \times 2 = 0.8$ |
| $B$ cells | $O_C/n^B = 5/3 \approx 1.67$ | $O_F/n^B = 2/3$ | $d_B = 5/2 = 2.5$ | $\frac{2}{3} \times 2.5 = \frac{5}{3}$ |

The $B$ cells touch $A$ more often than a typical candidate (2/3 against 0.4), and those that touch $A$ have more $A$ neighbours than a typical candidate that touches $A$ (2.5 against 2).

**Edge cases.** Depth needs $K > 0$. For the candidates, $q = 0$ means no candidate has an $A$ neighbour; then every $x_\ell$ is 0, the image is uninformative (Section 3.5) and is dropped, so $\bar{d}$ is always defined. For the $B$ cells, $O_F = 0$ means no $B$ cell has an $A$ neighbour, and $d_B$ is an average over no cells. It is not needed: below, $d_B$ appears only multiplied by $O_F/n^B = 0$, so that term is 0 whatever value $d_B$ is given.

**Proposition 2.** *For any informative image,*

$$Y_C = \underbrace{\bar{d}\Big(\frac{O_F}{n^B} - q\Big)}_{\textit{breadth}} + \underbrace{\frac{O_F}{n^B}\,\big(d_B - \bar{d}\big)}_{\textit{depth}}.$$

*Proof.* By Section 3.4,

$$Y_C = \frac{O_C}{n^B} - \lambda.$$

Replace each term by its split, $O_C/n^B = (O_F/n^B)\,d_B$ and $\lambda = q\,\bar{d}$:

$$Y_C = \frac{O_F}{n^B}\,d_B - q\,\bar{d}.$$

Add and subtract $(O_F/n^B)\,\bar{d}$, which leaves $Y_C$ unchanged:

$$Y_C = \frac{O_F}{n^B}\,\bar{d} - q\,\bar{d} + \frac{O_F}{n^B}\,d_B - \frac{O_F}{n^B}\,\bar{d}.$$

The first two terms share the factor $\bar{d}$, and the last two share the factor $O_F/n^B$:

$$Y_C = \bar{d}\Big(\frac{O_F}{n^B} - q\Big) + \frac{O_F}{n^B}\,\big(d_B - \bar{d}\big).$$

If $O_F = 0$, then $O_C = 0$, the second line gives $Y_C = -q\,\bar{d}$, and so does the last line with the depth term equal to 0. □

**Reading the two terms.**

- *Breadth.* $O_F/n^B - q$ is the share of $B$ cells with an $A$ neighbour minus the share expected under random labelling. Multiplying by $\bar{d}$ turns it into $A$ neighbours per $B$ cell: each extra $B$ cell that touches $A$ brings the typical $\bar{d}$ $A$ neighbours with it. The term is 0 when the $B$ cells touch $A$ exactly as often as expected.

- *Depth.* $d_B - \bar{d}$ is how many more $A$ neighbours a $B$ cell that touches $A$ has than a typical candidate that touches $A$. It is multiplied by $O_F/n^B$ because only the $B$ cells that touch $A$ carry it. The term is 0 when the $B$ cells that touch $A$ sit in typical $A$ neighbourhoods, and positive when they sit where $A$ cells are more crowded.

In the example above, $Y_C = 5/3 - 0.8 \approx 0.87$. The breadth term is $2 \times (2/3 - 0.4) \approx 0.53$ and the depth term is $(2/3) \times (2.5 - 2) \approx 0.33$; together they give 0.87.

**The breadth term is the extra fraction.** Both are built from the excess $O_F - n^B q$, the number of extra $B$ cells with an $A$ neighbour ($E = n^B q$ by Corollary 1.2). The extra fraction divides it by the room, $Y_F = (O_F - n^B q)/R$, and the breadth term by $n^B$, so $O_F/n^B - q = (R/n^B)\, Y_F$.

**Corollary 2.1.** *For any informative image,*

$$Y_C = \bar{d}\,\frac{R}{n^B}\,Y_F + \frac{O_F}{n^B}\,\big(d_B - \bar{d}\big).$$

*Proof.* Substitute $O_F/n^B - q = (R/n^B)\, Y_F$ into the breadth term of Proposition 2. □

So the extra count is the extra fraction, multiplied by $\bar{d}\,R/n^B$, plus a depth term. In the example, the gain room is $R = n^B(1-q) = 1.8$, so $Y_F = (2 - 1.2)/1.8 \approx 0.44$ and $\bar{d}\,(R/n^B)\, Y_F = 2 \times 0.6 \times 0.44 \approx 0.53$, the breadth term found above.

Corollary 2.1 has three consequences for how the two effects are read.

1. *The extra count can change while the extra fraction does not.* Take $n^B = 100$ and $q = 0.3$, with the gain room $R = 70$, and let 51 $B$ cells have an $A$ neighbour, so $Y_F = (51 - 30)/70 = 0.3$. With $\bar{d} = 2$: if $d_B = 2$, then $Y_C = 2 \times 0.7 \times 0.3 = 0.42$; if $d_B = 3$, then $Y_C = 0.42 + 0.51 \times 1 = 0.93$. The extra fraction is 0.3 in both cases; only depth has changed. The extra count alone cannot say why it is high. A large extra count may mean that more $B$ cells touch $A$ (breadth), or that the $B$ cells touching $A$ sit where $A$ cells are more crowded (depth). The extra fraction measures breadth alone, so reporting it alongside the extra count tells the two apart.

2. *The extra count is not a rescaled extra fraction.* Even with no depth term, the extra count is the extra fraction multiplied by $\bar{d}\,R/n^B$. This factor differs between images: $R/n^B$ depends on $q$, and $\bar{d}$ on how densely the $A$ cells are packed. In the example above, if $\bar{d}$ were 4 instead of 2, the same $Y_F = 0.3$ with $d_B = \bar{d}$ would give $Y_C = 0.84$ instead of 0.42. $B$ cells placed in the same way can give different extra counts in tissues where the $A$ cells are packed differently. A difference in the extra count between groups may therefore come from the tissue rather than from the $B$ cells. Section 3.7 makes this precise under two models.

3. *The two terms can have opposite signs.* If the $B$ cells touch $A$ more often than expected but only at the edges of $A$ regions, the breadth term is positive and the depth term negative. The extra count can then be near 0 while the extra fraction is clearly positive. The two effects can disagree, and neither is wrong. They answer different questions, so choosing an effect means choosing the question. The disagreement is itself informative. It suggests that the $B$ cells sit at the edges of $A$ regions rather than inside them.

#### 3.7 Co-localisation or tissue? The assumptions behind each effect

Many methods call a pair of cell types differently co-localised whenever the score they compute differs between groups. But the tissues imaged can differ between groups, by chance or because of what distinguishes the groups: one group may have more of a cell type, denser tumour nests, more holes or tears, or smaller fields of view. Such differences can move a co-localisation score even when the cells behave the same way. For example, if the $A$ cells are packed more densely in one group, a $B$ cell next to them has more $A$ neighbours there, so the extra count rises even though no more $B$ cells are drawn to $A$ cells. Calling this pair changed would be a call confounded by the tissue. Section 3.6 already showed that the extra count can change because the tissue changes (consequence 2 of Corollary 2.1). This raises two questions: can the extra fraction also change because the tissue changes, and when an effect differs between groups, was the cause a change in how $B$ cells co-localise with $A$ cells or a change in the tissue?

**There is no single definition of co-localisation.** Loosely, two cell types co-localise when $B$ cells are found near $A$ cells more often than chance would place them. Turning this into a number takes three choices, and different methods make them differently.

1. *What chance means.* Here it is random labelling of the cells imaged (Section 3.2). Ripley's $K$ and its relatives instead take cells to be scattered evenly over the area imaged.

2. *What is summarised.* Whether a $B$ cell has any $A$ neighbour (breadth, the extra fraction), how many it has (breadth and depth, the extra count), or how surprising the count is (a $z$-score).

3. *What counts as a change in co-localisation.* This needs a model of how $B$ cells come to be near $A$ cells, which says what it means for $B$ cells to "behave the same way" in two tissues.

The data cannot make these choices, and methods that make them differently can disagree on the same images. So a statement that co-localisation differs between groups is a statement about co-localisation as defined by these choices. This section makes the third choice explicit.

The comparison of groups in Section 5 is a valid test of whether the effect differs, whatever the biology. Saying why it differs, through a change in how $B$ cells relate to $A$ cells or through a change in the tissue, needs the effect to stay the same when the $B$ cells behave the same way but the tissue differs. The tissue can differ, for example, in how much of it is near $A$ ($q$), or in how densely the $A$ cells are packed. Sections 3.7.1 and 3.7.2 give two models of the mechanism, label shift and hub recruitment. In both, a share $f$ of the $B$ cells is drawn to $A$, and $f$ is the behaviour: under these models, co-localisation is $f$. Under both, the extra fraction has expected value

$$\mathbb{E}(Y_F) = \frac{f(1-p)}{1-fp}, \qquad p := \frac{n^B}{M},$$

which depends only on $f$ and on the share $p$ of candidates that are $B$ cells, not on $q$ or on how the $A$ cells are packed (Propositions 3 and 4). The extra count depends on these quantities as well, through $\bar{d}$ and $q$ under label shift, and through the packing of $A$ cells around a hub under hub recruitment.

Propositions 3 and 4 hold only under their models. Under a different model of how $B$ cells come to be near $A$ cells, for example one in which being near $A$ raises the odds of a candidate being a $B$ cell, the extra fraction also changes with $q$. Neither effect is free of the tissue under every model, and the two models below are not the only possible ones. The same holds for any co-localisation measure, ours or another method's. Each summarises where $B$ cells sit, and reading a difference in it as a change in behaviour requires an assumption about how they came to be there. The two models are chosen because they are simple and describe attraction as it is usually pictured: some $B$ cells are where they are because of the $A$ cells, and the rest are not.

The next two subsections derive the expected extra fraction and extra count under each model. Both models describe attraction, a share of the $B$ cells drawn towards the $A$ cells, and both use the radius graph. Avoidance, a share of the $B$ cells kept away from the $A$ cells, is not modelled. Section 3.7.4 then uses them to choose between the effects. Throughout, we consider one image and one pair $A \to B$ with $A \neq B$, and use the notation of Section 3.6: $Y_F$ and $Y_C$ are the extra fraction and the extra count, $O_F$ and $O_C$ their observed totals, $c_\ell$ the number of $A$ cells within $r$ of candidate $\ell$, and $p = n^B/M$ the share of candidates that are $B$ cells. In both models a share $f$ of the $B$ cells is *drawn to $A$*, with $0 \leq f \leq 1$ and $fn^B$ a whole number. The extra fraction uses the gain room $R = n^B(1-q)$ (Section 3.3).

##### 3.7.1 Label shift

In label shift the cells and their positions are fixed, and only the labels move. So $q$, $\lambda$ and $\bar{d}$ are fixed.

**Definition 1** (Label shift)**.** Suppose at least $fn^B$ candidates have an $A$ neighbour ($fn^B \leq Mq$). The $B$ labels are placed in two steps.

(i) $fn^B$ of them go to a uniformly random set of $fn^B$ candidates with an $A$ neighbour;

(ii) the other $n^B - fn^B$ go to a uniformly random set of $n^B - fn^B$ of the $M - fn^B$ candidates not chosen in (i).

With $f = 0$ this is random labelling.

**Proposition 3** (Label shift)**.** *Under label shift,*

$$\mathbb{E}(Y_F) = \frac{f(1-p)}{1-fp}, \qquad \mathbb{E}(Y_C) = \frac{f(1-p)}{1-fp}\,\bar{d}\,(1-q).$$

*Proof. The $B$ cells drawn to $A$.* Each of the $fn^B$ cells chosen in (i) has an $A$ neighbour, so together they add $fn^B$ to $O_F$. Each is a uniformly random candidate with an $A$ neighbour, so its expected $c_\ell$ is the mean of $c_\ell$ over those candidates, which is $\bar{d}$ (Section 3.6). Together they add $fn^B\,\bar{d}$ to $O_C$ in expectation.

*What is left for the other $B$ cells.* Removing the $fn^B$ chosen candidates leaves $M - fn^B$ candidates, of which $Mq - fn^B$ have an $A$ neighbour. The $c_\ell$ of all candidates sum to $M\lambda$, and the chosen ones take $fn^B\,\bar{d}$ of that in expectation, so the remaining candidates' $c_\ell$ sum to $M\lambda - fn^B\,\bar{d}$ in expectation. Each of the other $n^B - fn^B$ $B$ cells sits at a uniformly random remaining candidate. So it has an $A$ neighbour with probability

$$\frac{Mq - fn^B}{M - fn^B} = \frac{q - fp}{1 - fp},$$

dividing the numerator and the denominator by $M$, and its expected $c_\ell$ is, in the same way,

$$\frac{M\lambda - fn^B\,\bar{d}}{M - fn^B} = \frac{\lambda - fp\,\bar{d}}{1 - fp}.$$

*The expected totals.* Adding the two groups of $B$ cells, and dividing by $n^B$,

$$\frac{\mathbb{E}(O_F)}{n^B} = f + (1-f)\,\frac{q - fp}{1 - fp}, \qquad \frac{\mathbb{E}(O_C)}{n^B} = f\,\bar{d} + (1-f)\,\frac{\lambda - fp\,\bar{d}}{1 - fp}.$$

*The extra fraction's excess.* Subtract $q$ and put everything over $1 - fp$:

$$\frac{\mathbb{E}(O_F)}{n^B} - q = \frac{f(1-fp) + (1-f)(q-fp) - q(1-fp)}{1-fp}.$$

Expanding the numerator,

$$f - f^2p + q - fp - fq + f^2p - q + fpq = f - fp - fq + fpq = f(1-p)(1-q).$$

So $\mathbb{E}(O_F) - n^Bq = n^B\,f(1-p)(1-q)/(1-fp)$.

*The extra fraction.* The room $R = n^B(1-q)$ is fixed, because $q$ is. Dividing the expected excess by $R$, the factor $n^B(1-q)$ cancels and

$$\mathbb{E}(Y_F) = \frac{\mathbb{E}(O_F) - n^Bq}{n^B(1-q)} = \frac{f(1-p)}{1-fp}.$$

*The extra count.* By Section 3.4, $\mathbb{E}(Y_C) = \mathbb{E}(O_C)/n^B - \lambda$. In the same way,

$$\frac{\mathbb{E}(O_C)}{n^B} - \lambda = \frac{f\,\bar{d}(1-fp) + (1-f)(\lambda - fp\,\bar{d}) - \lambda(1-fp)}{1-fp},$$

and the numerator is

$$f\bar{d} - f^2p\bar{d} + \lambda - fp\bar{d} - f\lambda + f^2p\bar{d} - \lambda + fp\lambda = f\bar{d} - fp\bar{d} - f\lambda + fp\lambda = f(1-p)(\bar{d} - \lambda).$$

Since $\lambda = q\,\bar{d}$ (Section 3.6), $\bar{d} - \lambda = \bar{d}\,(1-q)$, which gives the result. □

When $B$ cells are a small share of the candidates ($p$ near 0), $\mathbb{E}(Y_F)$ is close to $f$: the extra fraction estimates the share of $B$ cells drawn to $A$. The factor $(1-p)/(1-fp)$ arises because the $B$ cells that are not drawn to $A$ are placed among the candidates left over, which are slightly less often near $A$ than the candidates as a whole. The expected extra count is the expected extra fraction multiplied by $\bar{d}\,(1-q)$, which matches Corollary 2.1 with $R/n^B = 1 - q$ and no depth term: a $B$ cell drawn to $A$ goes to a typical candidate with an $A$ neighbour.

##### 3.7.2 Hub recruitment

In label shift, the $B$ cells drawn to $A$ take the places of candidates that were already near $A$. In hub recruitment they are instead moved: each is placed close to an $A$ cell, its *hub*. Cells move, so $q$ and $\lambda$ are computed after the move. We describe the image by the candidates that are not moved, whose positions are fixed: write $q_0$ for the share of them with an $A$ neighbour and $\lambda_0$ for their mean $c_\ell$.

**Definition 2** (Hub recruitment)**.** Fix a distance $\varrho \leq r$, how close to its hub a recruited cell lands. The image has $M$ candidates, $n^B$ of them $B$ cells.

(i) $fn^B$ of the $B$ cells are *recruited*. Each lies at a point drawn uniformly from the disc of radius $\varrho$ around its hub, an $A$ cell chosen uniformly at random, independently for each recruited cell. Every such disc lies inside the image.

(ii) The other $n^B - fn^B$ $B$ labels go to a uniformly random set of the $M - fn^B$ candidates that are not recruited.

Write $\lambda_0 + \rho_A$ for the expected number of $A$ cells other than the hub within $r$ of a recruited cell. The number $\rho_A$ defines how crowded the $A$ cells are around a hub. It is the number of $A$ cells, beyond the hub, near a recruited cell, minus the number near a typical candidate that is not moved. It is positive when $A$ cells cluster, as in tumour nests, and can be negative when they are spread out evenly.

**Proposition 4** (Hub recruitment)**.** *Under hub recruitment, with expectations over the recruitment and the labels,*

$$\mathbb{E}(Y_F) = \frac{f(1-p)}{1-fp}, \qquad \mathbb{E}(Y_C) = f(1-p)(1+\rho_A).$$

*Proof. Moving a $B$ cell changes no one else's $c_\ell$.* $c_\ell$ counts $A$ cells only, and a recruited cell is a $B$ cell, so placing it anywhere leaves every other candidate's $c_\ell$ unchanged.

*Recruited cells have an $A$ neighbour.* A recruited cell is within $\varrho$ of its hub, and $\varrho \leq r$, so it always has an $A$ neighbour. Its expected $c_\ell$ is its hub plus the others, $1 + \lambda_0 + \rho_A$.

*The share $q$ after recruitment.* Of the $M$ candidates, $M - fn^B$ are not moved, and a share $q_0$ of them has an $A$ neighbour; the $fn^B$ recruited cells all have one. So

$$q = \frac{(M - fn^B)\, q_0 + fn^B}{M} = q_0 - fp\, q_0 + fp = q_0 + fp(1-q_0),$$

and $1 - q = (1-q_0)(1-fp)$. This does not depend on where the recruited cells land, so $q$ and the room $R = n^B(1-q) = n^B(1-q_0)(1-fp)$ are fixed.

*The extra fraction's excess.* The $fn^B$ recruited cells add $fn^B$ to $O_F$. The other $n^B - fn^B$ $B$ cells are uniformly random candidates that are not moved, each with an $A$ neighbour with probability $q_0$. So

$$\mathbb{E}(O_F) = fn^B + n^B(1-f)\, q_0,$$

and, using $n^B q = n^B q_0 + n^B fp(1-q_0)$,

$$\begin{aligned}\mathbb{E}(O_F) - n^B q &= fn^B + n^B q_0 - fn^B q_0 - n^B q_0 - fn^B p(1-q_0) \\ &= fn^B(1-q_0) - fn^B p(1-q_0) = fn^B(1-p)(1-q_0).\end{aligned}$$

*The extra fraction.* Dividing by the fixed room,

$$\mathbb{E}(Y_F) = \frac{fn^B(1-p)(1-q_0)}{n^B(1-q_0)(1-fp)} = \frac{f(1-p)}{1-fp}.$$

*The extra count.* The recruited cells add $fn^B(1+\lambda_0+\rho_A)$ to $O_C$ in expectation, and the others $n^B(1-f)\lambda_0$, so

$$\frac{\mathbb{E}(O_C)}{n^B} = \lambda_0 + f(1+\rho_A).$$

Here $\lambda$ is not fixed, because it includes the recruited cells' $c_\ell$, so we take its expectation too. The candidates that are not moved contribute $(M - fn^B)\lambda_0$ to the sum of $c_\ell$, and the recruited ones $fn^B(1+\lambda_0+\rho_A)$ in expectation, so

$$\mathbb{E}(\lambda) = \frac{(M-fn^B)\lambda_0 + fn^B(1+\lambda_0+\rho_A)}{M} = \lambda_0 + fp(1+\rho_A).$$

Since $Y_C = O_C/n^B - \lambda$,

$$\mathbb{E}(Y_C) = \lambda_0 + f(1+\rho_A) - \lambda_0 - fp(1+\rho_A) = f(1-p)(1+\rho_A).$$

□

The expected extra fraction is the same as under label shift and does not involve $q_0$ or $\rho_A$. The expected extra count involves $\rho_A$, which depends on how the $A$ cells are arranged: in the same tissue with the same $f$, tumour nests that are more densely packed give a larger $\rho_A$ and a larger extra count.

**When $\varrho > r$.** Proposition 4 needs $\varrho \le r$. If $\varrho > r$, a recruited cell may have no $A$ cell within $r$, and the share $e$ of recruited cells that do depends on how the $A$ cells are packed. The same steps then give an expected excess of $fn^B(1-p)(e-q_0)$, so the extra fraction depends on the packing too. Neither proposition covers this case. So the extra fraction is free of the tissue only when the radius is at least the distance at which $B$ cells gather around $A$ cells, which is one more reason not to choose the radius too small.

##### 3.7.3 Summary

| model | $\mathbb{E}(Y_F)$ | $\mathbb{E}(Y_C)$ |
|---|---|---|
| label shift | $\frac{f(1-p)}{1-fp}$ | $\frac{f(1-p)}{1-fp}\,\bar{d}\,(1-q)$ |
| hub recruitment ($\varrho \le r$) | $\frac{f(1-p)}{1-fp}$ | $f(1-p)(1+\rho_A)$ |

Under both models the expected extra fraction depends only on the behaviour $f$ and on $p$. The factor $(1-p)/(1-fp)$ lies between $1-p$ and 1, so it matters only when $B$ cells are a large share of the candidates. If that share differs a lot between groups, the extra fraction differs somewhat too: with $f = 0.2$, it is 0.184 at $p = 0.1$ and 0.167 at $p = 0.2$. The expected extra count depends, in addition, on how the $A$ cells are laid out: through $q$ and $\bar{d}$ under label shift, and through $\rho_A$ under hub recruitment.

###### 3.7.4 Choosing an effect

We make the extra fraction the default of our method. For an attracting pair, its expected value under both models depends only on the share $f$ of $B$ cells drawn to $A$ (and on $p$), so a difference between groups can be read as a difference in $f$ even when the tissue differs. The extra count is the better choice when the question is how much $A$ surrounds the $B$ cells and the groups' $A$ tissue is comparable. The rest of this subsection sets out what each effect answers, how much the tissue moves each one under the two models, how to check whether the $A$ tissue is comparable, and how to read the two effects together.

**The question each effect answers.**

- The *extra fraction* answers: do $B$ cells touch $A$ cells more often than expected? Each $B$ cell contributes 1 if it has an $A$ neighbour and 0 otherwise, so the extra fraction measures breadth only.
- The *extra count* answers: how many more $A$ cells surround a $B$ cell than expected? Each $B$ cell contributes its number of $A$ neighbours, so the extra count measures breadth and depth together (Proposition 2).

Neither question is more correct than the other. The choice depends on which question is asked, and on whether its answer can be compared between groups whose tissues differ.

**What we want from an effect.** The comparison of groups asks whether the effect differs between them. We would like a difference to mean that the $B$ cells behave differently, here that the share $f$ of $B$ cells drawn to $A$ differs. That requires two images with the same $f$ to have the same expected effect, even when their tissues differ. Tissues can differ in two ways that matter here:

- in how much of the image is near $A$, measured by $q$;
- in how densely the $A$ cells are packed, measured by $\bar{d}$ (the typical number of $A$ neighbours of a candidate that has one) under label shift and by $\rho_A$ (how crowded the $A$ cells are around a hub) under hub recruitment.

**The extra fraction under both models.** The expected extra fraction, $f(1-p)/(1-fp)$, contains neither $q$ nor $\bar{d}$ nor $\rho_A$. With $f = 0.2$ it is $0.18/0.98 \approx 0.184$ at $p = 0.1$ and 0.167 at $p = 0.2$, whatever the tissue.

**The extra count under label shift.** By Proposition 3, the expected extra count is the expected extra fraction multiplied by $\bar{d}\,(1-q)$. With $f = 0.2$ and $p = 0.1$, as above:

| $q$ | $\bar{d}$ | $\mathbb{E}(Y_C)$ |
|---|---|---|
| 0.3 | 2 | $0.184 \times 2 \times 0.7 \approx 0.26$ |
| 0.3 | 4 | $0.184 \times 4 \times 0.7 \approx 0.51$ |
| 0.6 | 2 | $0.184 \times 2 \times 0.4 \approx 0.15$ |

The behaviour is the same in every row. Doubling the packing of the $A$ cells doubles the extra count, and raising $q$ from 0.3 to 0.6 nearly halves it. A comparison of the extra count between groups whose tissues differ like this would find a difference that the $B$ cells did not cause.

**The extra count under hub recruitment.** By Proposition 4, the expected extra count is $f(1-p)(1+\rho_A)$. It does not contain $q$, but it contains $\rho_A$. With $f = 0.2$ and $p = 0.1$, it is $0.18 \times 2 = 0.36$ when $\rho_A = 1$ and $0.18 \times 4 = 0.72$ when $\rho_A = 3$: denser nests of $A$ cells double the extra count with the same behaviour.

**A simulation of hub recruitment.** To see how large these tissue effects are in practice, we simulated hub recruitment with the same behaviour in two groups and different $A$ tissue.

- *Images.* Each image is $1000 \times 1000$ units with $r = 20$. The $A$ cells (a tumour) form nests: discs of radius 60 placed at random, each with about 150 cells. There are 3,400 candidates, of which 400 are $B$ cells.
- *Groups.* Ten patients per group, one image each. In the first group, images have about 15 nests. In the second, the tumour is either the same, or has more nests (30), fewer nests (3), or denser nests (15 nests of about 300 cells).
- *Behaviour.* In both groups, a share $f = 0.1$ or $f = 0.3$ of the $B$ cells is recruited, with $\varrho = r = 20$ (Definition 2). So the $B$ cells behave the same way, and any difference the test finds is caused by the tissue.
- *Analysis.* Each simulated data set is analysed with both effects, with an R implementation of the estimate and test of Sections 4 and 5 for one image per patient, without the rules of Section 5.8. This is repeated 500 times, and we record how often the test gives $p < 0.05$. A test that is not misled by the tissue does so about 5% of the time.

The results agree with Proposition 4.

- *Extra fraction.* The mean extra fraction was 0.089–0.090 for $f = 0.1$ and 0.274–0.275 for $f = 0.3$ in every tissue setting, against $f(1-p)/(1-fp) = 0.089$ and 0.274 with $p = 400/3400$. The test gave $p < 0.05$ in 3.6–7.0% of data sets, close to 5% in every setting.
- *Extra count.* The mean number of $A$ cells within $r$ of a point near a hub was 18.3 with 15 nests, 22.3 with 30 nests (more nests overlap), 15.2 with 3 nests and 35.4 with denser nests. So $\rho_A$ changed by about 20% when only the number of nests changed, and roughly doubled with denser nests. With $f = 0.3$, the extra count's test gave $p < 0.05$ in 14% of data sets with more or fewer nests, and in 100% with denser nests, although the $B$ cells behaved the same way in both groups. With $f = 0.1$ the rates were 8% and 7% with more or fewer nests, and 99% with denser nests.

So $\rho_A$ can differ between groups even when the $A$ tissue looks similar, and the extra count then differs with it.

**When the extra count is the right choice.** The extra count is the better effect when two things hold.

1. *Depth is the question.* If what matters is how much $A$ surrounds the $B$ cells, for example the number of tumour cells a T cell can contact, the extra count measures it and the extra fraction does not.

2. *The $A$ tissue is comparable between the groups.* Then $q$, $\bar{d}$ and $\rho_A$ are similar in the two groups, the tissue terms above are the same on both sides, and a difference in the extra count can be read as a difference in behaviour. Whether this holds can be checked from the data: $q$ and $\bar{d} = \lambda/q$ are computed for every image, so their spread in each group can be compared before the extra count is interpreted.

If the $A$ tissue differs between the groups, a difference in the extra count is still a valid finding that the extra count differs, but it cannot be attributed to the $B$ cells without further evidence, such as similar $q$ and $\bar{d}$ in the two groups.

**Reading the two effects together.** When both effects are tested, the four combinations of results read as follows, under the two models. Here "yes" means that the test found a significant difference, and "no" means that it did not.

| extra fraction differs | extra count differs | reading |
|---|---|---|
| yes | yes | $B$ cells touch $A$ more often in one group. The size of the extra count difference also reflects depth and the $A$ tissue. |
| no | yes | No evidence of a difference in how often $B$ cells touch $A$. The $B$ cells that touch $A$ sit among more $A$ cells in one group, either because they prefer crowded $A$ regions (depth, Proposition 2) or because the $A$ cells are packed more densely there (tissue). Comparing $q$ and $\bar{d}$ between the groups helps separate the two. |
| yes | no | $B$ cells touch $A$ more often in one group, but the extra contact is offset in the extra count, for example because it is at the edges of $A$ regions (consequence 3 in Section 3.6), or the extra count's test lacked the power to detect it. |
| no | no | No evidence of a difference at this radius. This is not evidence that there is no difference. |

A "no" in the table means only that no difference was found. It can mean that there is no difference, or that the test was underpowered: too few patients, a rare cell type that leaves many images uninformative, or a difference smaller than the test can detect (Section 6.5). The two effects can also differ in power for the same pair, so a "yes" for one and a "no" for the other does not by itself show that only one has changed.

#### 3.8 Self-pairs

**Why self-pairs differ.** For a pair $A \to B$, each candidate's number $x_\ell$ is set by the $A$ cells, and random labelling does not move them. That is why $O$ is a sum of fixed numbers over a random set (Corollary 1.1). For a self-pair $A \to A$, the $A$ labels are the ones being shuffled. So the same labels decide both which cells are counted and which cells are their neighbours, and shuffling moves both. Each $x_\ell$ is then itself a sum of labels. Proposition 1 does not apply, and the mean $E$ and variance $V$ have to be derived again.

Section 3.8.1 sets up the notation and one fact about labels that every proof uses. Sections 3.8.2 and 3.8.3 derive $E$ and $V$ for the extra fraction and the extra count, each with its proof, and Section 3.8.4 sums up what each image gives.

##### 3.8.1 Setup

Take one image and one self-pair $A \to A$. The image has $n$ cells and $n^A$ of them are $A$ cells.

- *Random labelling.* The positions of all cells are fixed, and the $n^A$ $A$ labels are placed on $n^A$ of the $n$ cells, with every set of $n^A$ cells equally likely. Every cell can carry an $A$ label, so all $n$ cells play the part of the candidates.
- *Labels.* $I_\ell = 1$ if cell $\ell$ is an $A$ cell and $I_\ell = 0$ otherwise.
- *Probabilities.* $\pi_k$ is the probability that $k$ given cells are all $A$ cells. By the same argument as (2),
$$\pi_k := \frac{n^A}{n} \cdot \frac{n^A - 1}{n-1} \cdots \frac{n^A - k + 1}{n - k + 1}.$$
- *Neighbours and links.* $\mathcal{N}(\ell)$ is the set of neighbours of cell $\ell$, not including $\ell$ itself. A *link* is an ordered pair of cells $(\ell, j)$ with $j \in \mathcal{N}(\ell)$: it starts at $\ell$ and ends at $j$. For each cell $\ell$, $d_\ell = |\mathcal{N}(\ell)|$ links start at $\ell$, $w_\ell$ links end at $\ell$ (the cells that have $\ell$ as a neighbour), and $t_\ell = d_\ell + w_\ell$ links touch $\ell$. In all there are $m_{\mathrm{L}} = \sum_\ell d_\ell$ links, which make up the set $\mathcal{L}$, and $b$ of them have their reverse as a link. With the radius graph every link goes both ways, so $w_\ell = d_\ell$ and $b = m_{\mathrm{L}}$; with the $k$-nearest-neighbour graph this need not hold.

The observed totals are as follows.

- extra fraction: $O$ is the number of $A$ cells with at least one other $A$ cell among their neighbours;
- extra count: $O$ is the total number of $A$ neighbours of the $A$ cells. A pair of $A$ cells that are each other's neighbours counts twice, once from each cell.

**The mean of a product of labels.** The proofs below need the means of products of labels, such as $I_\ell I_j$. One fact gives all of them. Here and below, a binomial coefficient $\binom{a}{c}$ with $a < c$ is 0.

**Lemma 1.** *Let $\ell_1, \dots, \ell_s$ be cells, not necessarily different, and let $k$ be the number of different cells among them. Under random labelling,*

$$\mathbb{E}\big(I_{\ell_1} I_{\ell_2} \cdots I_{\ell_s}\big) = \pi_k.$$

*Proof.* Each $I_\ell$ is 0 or 1, so $I_\ell^2 = I_\ell$, and a label that appears more than once in the product can be kept once. The product is then a product over $k$ different cells. It is 1 exactly when all $k$ cells are $A$ cells, and 0 otherwise. So its mean is the probability that $k$ given cells are all $A$ cells, which is $\pi_k$. □

So the mean of a product of labels depends only on how many different cells it involves. The proofs of the variances below come down to counting, for each term, how many different cells it involves.

##### 3.8.2 Extra fraction

Two probabilities are needed. Given that cell $\ell$ is an $A$ cell, the other $n^A - 1$ labels are on $n^A - 1$ of the other $n - 1$ cells, every set equally likely. The chance that none of them is among $s$ given cells is the number of ways to place them on the remaining $n - 1 - s$ cells, divided by the number of ways to place them on all $n - 1$:

$$h(s) := \binom{n-1-s}{n^A-1} \Big/ \binom{n-1}{n^A-1}.$$

In the same way, given that two cells are $A$ cells, the chance that none of $s$ given other cells is an $A$ cell is

$$h_2(s) := \binom{n-2-s}{n^A-2} \Big/ \binom{n-2}{n^A-2}.$$

For two different cells $\ell$ and $\ell'$, let $T_{\ell\ell'}$ be the probability that both have an $A$ neighbour, given that both are $A$ cells. With $a = |\mathcal{N}(\ell) \setminus \{\ell'\}|$, $a' = |\mathcal{N}(\ell') \setminus \{\ell\}|$ and $c = |\mathcal{N}(\ell) \cap \mathcal{N}(\ell')|$ (the proof below explains the four cases),

$$T_{\ell\ell'} = \begin{cases} 1 & \text{if } \ell' \in \mathcal{N}(\ell) \text{ and } \ell \in \mathcal{N}(\ell'), \\ 1 - h_2(a') & \text{if } \ell' \in \mathcal{N}(\ell) \text{ only}, \\ 1 - h_2(a) & \text{if } \ell \in \mathcal{N}(\ell') \text{ only}, \\ 1 - h_2(a) - h_2(a') + h_2(a + a' - c) & \text{otherwise.} \end{cases}$$

**Proposition 5** (Self-pair extra fraction)**.** *Under random labelling, the extra fraction's observed total $O$ has mean and variance*

$$E = \frac{n^A}{n} \sum_{\ell=1}^{n} \big\{1 - h(d_\ell)\big\}, \qquad V = \operatorname{Var}(O) = E - E^2 + \pi_2 \sum_{\ell \neq \ell'} T_{\ell\ell'}.$$

*Proof.* Let $Z_\ell = 1$ if at least one neighbour of $\ell$ is an $A$ cell. Then $O = \sum_\ell I_\ell Z_\ell$, and

$$\mathbb{E}(O) = \sum_\ell \Pr(I_\ell = 1 \text{ and } Z_\ell = 1) = \sum_\ell \Pr(I_\ell = 1) \Pr(Z_\ell = 1 \mid I_\ell = 1).$$

First, $\Pr(I_\ell = 1) = n^A/n$, by (1) with $n$ cells. Second, $Z_\ell = 0$ given $I_\ell = 1$ exactly when none of the $d_\ell$ neighbours of $\ell$ is an $A$ cell, which has probability $h(d_\ell)$. So $\Pr(Z_\ell = 1 \mid I_\ell = 1) = 1 - h(d_\ell)$, and summing over $\ell$ gives $E$.

*The variance.* The variance is $\mathbb{E}(O^2) - E^2$, so it remains to find the mean of $O^2$. Squaring $O$ gives one term for each cell and one for each ordered pair of different cells:

$$O^2 = \sum_{\ell} I_\ell^2 Z_\ell^2 + \sum_{\ell \neq \ell'} I_\ell Z_\ell \, I_{\ell'} Z_{\ell'}.$$

For one cell, $I_\ell$ and $Z_\ell$ are 0 or 1, so $I_\ell^2 Z_\ell^2 = I_\ell Z_\ell$. These terms add up to $O$, whose mean is $E$. For two different cells, the term is 1 exactly when both are $A$ cells and both have an $A$ neighbour, so

$$\mathbb{E}(I_\ell Z_\ell \, I_{\ell'} Z_{\ell'}) = \Pr(I_\ell = I_{\ell'} = 1) \, \Pr(Z_\ell = Z_{\ell'} = 1 \mid I_\ell = I_{\ell'} = 1) = \pi_2 \, T_{\ell\ell'},$$

where $T_{\ell\ell'}$ is the probability defined before the proposition. Adding the two kinds of term and subtracting $E^2$ gives

$$\mathrm{Var}(O) = E - E^2 + \pi_2 \sum_{\ell \neq \ell'} T_{\ell\ell'}.$$

*The four cases of $T_{\ell\ell'}$.* It remains to check the formula for $T_{\ell\ell'}$. Suppose $\ell$ and $\ell'$ are both $A$ cells. The other $n^A - 2$ labels are on $n^A - 2$ of the other $n - 2$ cells, every set equally likely, so the chance that none of them is among $s$ given cells, other than $\ell$ and $\ell'$, is $h_2(s)$. Let $U = \mathcal{N}(\ell) \setminus \{\ell'\}$ and $U' = \mathcal{N}(\ell') \setminus \{\ell\}$. These are the neighbours of each cell other than the other one, with $|U| = a$ and $|U'| = a'$. Neither contains $\ell$ or $\ell'$, since a cell is not its own neighbour. Their common cells are the common neighbours of $\ell$ and $\ell'$, so $|U \cap U'| = c$ and $|U \cup U'| = a + a' - c$.

Now $Z_\ell = 1$ exactly when $\ell' \in \mathcal{N}(\ell)$ (since $\ell'$ is an $A$ cell) or some cell of $U$ is an $A$ cell, and likewise for $Z_{\ell'}$. This gives four cases.

1. *Each is a neighbour of the other.* Then $Z_\ell = Z_{\ell'} = 1$ for certain, and $T_{\ell\ell'} = 1$.
2. *$\ell' \in \mathcal{N}(\ell)$ only.* Then $Z_\ell = 1$ for certain, and $Z_{\ell'} = 1$ unless no cell of $U'$ is an $A$ cell. So $T_{\ell\ell'} = 1 - h_2(a')$.
3. *$\ell \in \mathcal{N}(\ell')$ only.* In the same way, $T_{\ell\ell'} = 1 - h_2(a)$.
4. *Neither.* Then $Z_\ell = 0$ when no cell of $U$ is an $A$ cell, and $Z_{\ell'} = 0$ when no cell of $U'$ is. Both are 0 when no cell of $U \cup U'$ is. By inclusion–exclusion, given $I_\ell = I_{\ell'} = 1$,

   $$T_{\ell\ell'} = 1 - \Pr(Z_\ell = 0) - \Pr(Z_{\ell'} = 0) + \Pr(Z_\ell = Z_{\ell'} = 0) = 1 - h_2(a) - h_2(a') + h_2(a + a' - c).$$

□

$E$ depends only on the number of neighbours of each cell, not on which cells are $A$. So everything in Section 3.3 carries over: the two rooms $n^A - E$ and $E$, the choice of scale, the rule for images with $R < 1$ (Section 3.5), and the effect $Y = (O - E)/R$, with $\mathbb{E}(Y) = 0$ and $\mathrm{Var}(Y) = s^2 = V/R^2$.

With the radius graph every link goes both ways, so cases 2 and 3 of the proof do not occur: two cells are either neighbours of each other (case 1) or not (case 4).

Only pairs of cells that are linked or share a neighbour need their own $T_{\ell\ell'}$. Every other pair has $c = 0$, so its $T_{\ell\ell'}$ depends only on $a$ and $a'$, and these pairs can be added up from the numbers of cells with each number of neighbours. With a radius, the terms of pairs that share a neighbour are by default estimated from a sample of cells (Remark 2).

*Remark* 2 (Sampling the shared-neighbour terms). With the radius graph, $d_\ell$ is the number of neighbours of $\ell$, and the pairs that need their own term are of two kinds.

- *Linked pairs.* Their term is 1. There are $\sum_\ell d_\ell$ of them, one for each link, so they take time of order $n\kappa$, where $\kappa$ is the average number of neighbours.

- *Unlinked pairs that share $c \geq 1$ neighbours.* Their term differs from that of a pair with $c = 0$ only in its last part, $h_2(a + a' - c)$ in place of $h_2(a + a')$. Finding them for a cell $\ell$ means visiting each neighbour $v$ of $\ell$ and each neighbour of $v$, which takes time of order $\sum_{v \in \mathcal{N}(\ell)} d_v$, about $\kappa^2$. Over all cells this is $\sum_v d_v^2$, of order $n\kappa^2$.

The second kind dominates. The number of neighbours grows with the area of the disc, $\kappa \approx \rho\pi r^2$ for cells at density $\rho$, so the time grows as $n\kappa^2 \propto n\rho^2 r^4$: doubling the radius multiplies it by 16, and doubling the density, for the same number of cells, by 4. For large images at large radii this is most of the time of the whole analysis.

So by default the second kind is estimated from a sample. For each cell $\ell$, let

$$g_\ell := \sum_{\ell'} \big\{ h_2(a + a' - c) - h_2(a + a') \big\},$$

the sum over the cells $\ell'$ not linked to $\ell$ that share $c \geq 1$ neighbours with it, so that the exact sum of $T_{\ell\ell'}$ is that of the pairs with $c = 0$, corrected for the linked pairs, plus $\sum_\ell g_\ell$. Choose a sample $\mathcal{S}$ of 50 cells in the image, and replace $\sum_\ell g_\ell$ by

$$\frac{n}{|\mathcal{S}|} \sum_{\ell \in \mathcal{S}} g_\ell.$$

- *It is unbiased.* Over a random choice of $\mathcal{S}$, each $g_\ell$ is included with probability $|\mathcal{S}|/n$, so the scaled sum has mean $\sum_\ell g_\ell$. $V$ is linear in this sum, so the estimated $V$, and hence $s^2$, has the exact value as its mean.

- *Everything else is exact.* The terms of pairs with $c = 0$ (from the counts of cells with each number of neighbours) and of linked pairs are computed in full.

- *The time falls to order $|\mathcal{S}|\,\kappa^2$ for the second kind, which no longer grows with $n$; the whole calculation is then of order $n\kappa$.*

- *The sample is reproducible.* $\mathcal{S}$ is the 50 cells with the smallest value of a fixed pseudo-random function of their coordinates. It behaves like one such random draw, but the same data always give the same sample, whatever the order of the cells.

With $|\mathcal{S}| = n$ (an image of at most 50 cells) the result is exact, and the exact calculation remains available for any image. Only the extra fraction's self-pairs with a radius are affected. For them $s_{ij}^2$ in Section 4 is an unbiased estimate rather than the exact value.

##### 3.8.3 Extra count

For every cell $j$, let $x_j$ be its number of $A$ neighbours, and let

$$L \coloneqq \sum_{j=1}^{n} x_j, \qquad E \coloneqq \tilde{p}L, \qquad \tilde{p} \coloneqq \frac{n^A - 1}{n - 1}.$$

$L$ is the number of links that end at an $A$ cell (with the radius graph, the total number of neighbours of the $A$ cells), and $\tilde{p}$ is the chance that the other end of a link is an $A$ cell. So $E$ is the number of $A$ neighbours expected from how crowded the $A$ cells are. $L$ is the self-pair version of $M\lambda$ for $A \to B$, and $E = \tilde{p}L$ that of $E = n^B\lambda$. Unlike for $A \to B$, $E$ is not a fixed number: random labelling moves the $A$ cells, and with them $L$. However, the effect is built from the excess $O - E$, so the quantities needed are its mean, which should be 0, and its variance.

**Proposition 6** (Self-pair extra count)**.** *Under random labelling, the extra count's excess has mean and variance*

$$\mathbb{E}(O - E) = 0, \qquad V = \mathrm{Var}(O - E) = \mathbb{E}(O^2) - 2\tilde{p}\,\mathbb{E}(OL) + \tilde{p}^2\,\mathbb{E}(L^2),$$

*where, with* $S = \sum_\ell t_\ell(t_\ell - 1)$,

$$\begin{aligned}
\mathbb{E}(O^2) &= (m_{\mathrm{L}} + b)\,\pi_2 + (S - 2b)\,\pi_3 + (m_{\mathrm{L}}^2 - m_{\mathrm{L}} + b - S)\,\pi_4,\\
\mathbb{E}(OL) &= \pi_2 \sum_\ell w_\ell t_\ell + \pi_3\Big(m_{\mathrm{L}}^2 - \sum_\ell w_\ell t_\ell\Big),\\
\mathbb{E}(L^2) &= \pi_1 \sum_\ell w_\ell^2 + \pi_2\Big(m_{\mathrm{L}}^2 - \sum_\ell w_\ell^2\Big).
\end{aligned}$$

*Proof.* $O$ adds, for each $A$ cell $\ell$, the number of its neighbours that are $A$, so it has one term for each link:

$$O = \sum_\ell I_\ell \sum_{j \in \mathcal{N}(\ell)} I_j = \sum_{(\ell,j)} I_\ell I_j.$$

$I_\ell I_j$ is 1 exactly when both cells are $A$ cells, so its mean is $\pi_2$, and $\mathbb{E}(O) = \pi_2\, m_{\mathrm{L}}$.

$L$ adds, for each cell $\ell$, the number of its neighbours that are $A$, so it also has one term for each link:

$$L = \sum_\ell \sum_{j \in \mathcal{N}(\ell)} I_j = \sum_{(\ell,j)} I_j.$$

Each $I_j$ has mean $\pi_1 = n^A/n$, so $\mathbb{E}(L) = \pi_1\, m_{\mathrm{L}}$. Finally, $\tilde{p}\,\pi_1 = \frac{n^A-1}{n-1} \cdot \frac{n^A}{n} = \pi_2$, so $\mathbb{E}(E) = \mathbb{E}(\tilde{p}L) = \pi_2\, m_{\mathrm{L}} = \mathbb{E}(O)$, and the excess has mean 0.

*The variance.* Write $X \coloneqq O - \tilde{p}L$. Its mean is 0, so its variance is the mean of its square. Expanding the square,

$$\mathrm{Var}(X) = \mathbb{E}(O^2) - 2\tilde{p}\,\mathbb{E}(OL) + \tilde{p}^2\,\mathbb{E}(L^2),$$

so the three means are found in turn. Each is a sum of means of products of labels, and by Lemma 1 each of these depends only on how many different cells the product involves. So each mean comes down to counting terms by their number of different cells. For this, write $O$ and $L$ as sums over the links,

$$O = \sum_{e \in \mathcal{L}} I_{e_1} I_{e_2}, \qquad L = \sum_{e \in \mathcal{L}} I_{e_2} = \sum_{v} w_v I_v,$$

where a link $e = (e_1, e_2)$ starts at $e_1$ and ends at $e_2$. The second form of $L$ groups the links by the cell they end at: cell $v$ is the end of $w_v$ links.

*The mean of* $O^2$. Squaring $O$ gives one term for each ordered pair of links $(e, f)$:

$$O^2 = \sum_{e \in \mathcal{L}} \sum_{f \in \mathcal{L}} I_{e_1} I_{e_2} I_{f_1} I_{f_2}.$$

Each link has two different cells, so the two links together involve 2, 3 or 4 different cells, and the term has mean $\pi_2$, $\pi_3$ or $\pi_4$ accordingly. Let $S_2$, $S_3$ and $S_4$ be the numbers of ordered pairs of each kind, so that

$$\mathbb{E}(O^2) = S_2\, \pi_2 + S_3\, \pi_3 + S_4\, \pi_4.$$

- *Two different cells.* Then $f$ has the same two cells as $e$, so $f = e$, or $f$ is the reverse of $e$. The first gives $m_{\mathrm{L}}$ pairs, one for each link. The second gives $b$ pairs, one for each link whose reverse is a link. So $S_2 = m_{\mathrm{L}} + b$.

- *Three different cells.* Then $e$ and $f$ share exactly one cell. Fix a cell $v$. Of the links, $t_v$ touch $v$, so there are $t_v(t_v - 1)$ ordered pairs of different links that both touch $v$. Adding this over all cells gives $S = \sum_v t_v(t_v - 1)$. A pair that shares exactly one cell is counted once, at that cell. A pair of different links that share two cells is a link and its reverse. There are $b$ such ordered pairs, and each is counted twice, once at each of its cells. So $S = S_3 + 2b$, and $S_3 = S - 2b$.

- *Four different cells.* These are all the other pairs. There are $m_{\mathrm{L}}^2$ ordered pairs in all, so $S_4 = m_{\mathrm{L}}^2 - S_2 - S_3 = m_{\mathrm{L}}^2 - m_{\mathrm{L}} + b - S$.

This gives $\mathbb{E}(O^2)$ as stated.

*The mean of* $OL$. Multiplying the two sums,

$$OL = \sum_{e \in \mathcal{L}} \sum_{v} w_v\, I_{e_1} I_{e_2} I_v.$$

For a given link $e$, the term involves two different cells if $v$ is one of $e_1$ and $e_2$, and three otherwise. So

$$\mathbb{E}(OL) = \sum_{e \in \mathcal{L}} \Big\{ \pi_2\, (w_{e_1} + w_{e_2}) + \pi_3 \Big( \sum_{v} w_v - w_{e_1} - w_{e_2} \Big) \Big\}.$$

Two sums simplify this.

- Every link ends at exactly one cell, so $\sum_v w_v = m_{\mathrm{L}}$.

- In $\sum_e (w_{e_1} + w_{e_2})$, each cell $v$ contributes $w_v$ once for each link that touches it, so the sum is $\sum_v w_v t_v$.

This gives $\mathbb{E}(OL)$ as stated.

*The mean of $L^2$.* Squaring the second form of $L$,

$$L^2 = \sum_u \sum_v w_u w_v \, I_u I_v.$$

The term involves one cell if $u = v$, and two otherwise. Since $\sum_u \sum_v w_u w_v = m_{\mathrm{L}}^2$, this gives $\mathbb{E}(L^2)$ as stated. Putting the three means into the expansion of $\mathrm{Var}(X)$ gives $V$. □

The factor is $\tilde{p} = (n^A - 1)/(n-1)$ rather than $n^A/n$ because a cell is not its own neighbour: given that a cell is an $A$ cell, each other cell is an $A$ cell with probability $(n^A - 1)/(n-1)$. With $n^A/n$ the excess would have mean $-m_{\mathrm{L}}(\pi_1^2 - \pi_2)$, slightly below 0. The extra count is $Y = (O - E)/n^A$ as in Section 3.4, with $\mathbb{E}(Y) = 0$ and $\mathrm{Var}(Y) = s^2 = V/(n^A)^2$.

With the radius graph every link goes both ways, so $w_v = d_v$, $t_v = 2d_v$ and $b = m_{\mathrm{L}}$, with $m_{\mathrm{L}} = \sum_v d_v$. Then $S = \sum_v 2d_v(2d_v - 1)$, and the three means become

$$\begin{aligned}
\mathbb{E}(O^2) &= 2m_{\mathrm{L}}\,\pi_2 + 4\sum_v d_v(d_v - 1)\,\pi_3 + \Big(m_{\mathrm{L}}^2 + 2m_{\mathrm{L}} - 4\sum_v d_v^2\Big)\pi_4, \\
\mathbb{E}(OL) &= 2\pi_2 \sum_v d_v^2 + \pi_3\Big(m_{\mathrm{L}}^2 - 2\sum_v d_v^2\Big), \\
\mathbb{E}(L^2) &= \pi_1 \sum_v d_v^2 + \pi_2\Big(m_{\mathrm{L}}^2 - \sum_v d_v^2\Big).
\end{aligned}$$

So $V$ depends on the image only through $n$, $n^A$, $\sum_v d_v$ and $\sum_v d_v^2$, and it takes $O(n\kappa)$ time to compute.

##### 3.8.4 Summary

For a self-pair, each image gives the following.

| | Extra fraction | Extra count |
|---|---|---|
| Observed total $O$ | $A$ cells with another $A$ cell among their neighbours | $A$ neighbours of the $A$ cells, summed over the $A$ cells |
| Expected value $E$ | $\frac{n^A}{n}\sum_\ell \{1 - h(d_\ell)\}$, a fixed number (Proposition 5) | $\tilde{p}L$, which changes with the labelling; $\mathbb{E}(O - E) = 0$ (Proposition 6) |
| Variance $V$ | $\mathrm{Var}(O)$ (Proposition 5) | $\mathrm{Var}(O - E)$ (Proposition 6) |
| Scale | the room $R$: $n^A - E$ (gain) or $E$ (loss), by the pair's scale | $n^A$ |
| Effect $Y$ | $(O - E)/R$ | $(O - E)/n^A$ |
| Mean and variance of $Y$ | 0 and $s^2 = V/R^2$ | 0 and $s^2 = V/(n^A)^2$ |
| Not used when | $V = 0$ or $R < 1$ (Section 3.5) | $V = 0$ (Section 3.5) |

For a self-pair $A \to A$, $V = 0$ exactly when $O$ takes the same value under every labelling: for example, when $n^A \leq 1$, when every cell is an $A$ cell, or when no cell has a neighbour.

From here on, $Y$ and $s^2$ are used in the same way for self-pairs as for any other pair.

### 4 Combining images within patients

Section 3 gives each image $(i, j)$ an effect $Y_{ij}$ and its variance $s^2_{ij}$ under random labelling. This section combines the images of each group into one estimate of the group's mean effect, with patients as the units. It is done separately for each pair and each radius.

The images used here are those left after the steps of Section 3: images with no group are removed, images with no $B$ cell or with $V = 0$ are not used, the scale is chosen, and images whose chosen room is below one cell are not used (Section 3.5). A patient none of whose images is left takes no part.

#### 4.1 The two-level model

**Combining images needs weights.** A group's images are combined by a weighted average, in which each image counts in inverse proportion to the variance of its effect: an image whose effect is known precisely counts more than one whose effect is noisy. This is the usual way to average numbers measured with different precision. It needs the full variance of each image's effect, and $s^2_{ij}$ is only part of it.

**Why $s^2$ is not enough.** $s^2_{ij}$ is the variance of $Y_{ij}$ when only the labels of one image are shuffled, with its cells where they are. The effects of real images vary more than that, for three reasons.

1. *Cells that move together.* Random labelling places each $B$ label on its own cell. The variance of a total of independent pieces grows with the size of the pieces: 100 coins counted one by one give a total with variance $100 \cdot \frac{1}{4} = 25$, while 10 coins each worth 10 give a total with the same mean and variance $10 \cdot 10^2 \cdot \frac{1}{4} = 250$. Real $B$ cells often lie in clumps that sit near $A$ cells, or away from them, as a whole. Then $O$ is a total of clumps rather than of single cells, and its variance is larger than $V$, by roughly the typical clump size. So the effect of a real image strays from its expected value by more than $s^2_{ij}$ allows, even when the two types do not co-localise.

2. *Images of the same patient differ.* Each image shows a different patch of tissue, such as the tumour edge in one image and its core in another, and the mix and arrangement of cells change from patch to patch. So two images of the same patient need not have the same effect.

3. *Patients of the same group differ.* Patients in the same group still differ in their biology: the subtype and stage of their disease, their treatment, and their immune response. In one patient $T$ cells may enter the tumour, while in another of the same group they stay at its edge. So two patients of the same group need not have the same effect.

So $s_{ij}^2$ alone understates how much an image's effect varies. To weight the images well, the model below adds terms that capture the rest of the variance.

**The model.** The effect of image $j$ of patient $i$ is written as

$$Y_{ij} = \delta_{c(i)} + b_i + \eta_{ij} + \varepsilon_{ij},$$

where

- $\delta_c$ is the mean effect of group $c$, the quantity compared in Section 5;
- $b_i$ is patient $i$'s own departure from $\delta_{c(i)}$, with mean 0 and variance $\tau^2$, the variance between patients. It covers reason 3;
- $\eta_{ij}$ is image $j$'s own departure from its patient beyond the random-labelling noise, with mean 0 and variance $\omega^2$, the variance between images of the same patient. It covers reasons 1 and 2;
- $\varepsilon_{ij}$ is the random-labelling noise of Section 3, with mean 0 and variance $s_{ij}^2$, which is known.

Reasons 1 and 2 share one term because both act image by image, and the data cannot tell them apart: both show up only as images of one patient differing more than $s^2$ allows.

**Independence.** All the $b_i$, $\eta_{ij}$ and $\varepsilon_{ij}$ are taken to be independent of each other. This rests on three assumptions.

- different patients are different people, and their images are independent (Section 2);
- $\eta_{ij}$ is measured from the patient's own level $b_i$, so how far an image strays from its patient is taken not to depend on where that level is, and one image's departure is taken not to depend on another's;
- $\varepsilon_{ij}$ is the chance in which of an image's cells carry which label, which is taken not to depend on the patient or the patch.

The last two assumptions are simplifications. Two neighbouring patches of one patient, for instance, may resemble each other more than two distant ones.

Under these assumptions,

$$\mathrm{Var}(Y_{ij}) = s_{ij}^2 + \omega^2 + \tau^2, \qquad \mathrm{Cov}(Y_{ij}, Y_{ij'}) = \tau^2 \ \ (j \neq j'), \qquad \mathrm{Cov}(Y_{ij}, Y_{i'j'}) = 0 \ \ (i \neq i').$$

Two images of the same patient share $b_i$, so they are correlated; images of different patients are not.

**The working covariance.** Collect patient $i$'s effects in the vector $\mathbf{Y}_i = (Y_{i1}, \ldots, Y_{iJ_i})^\top$. Its covariance under the model is the $J_i \times J_i$ matrix

$$\mathbf{V}_i \coloneqq \operatorname{diag}\big(s_{i1}^2 + \omega^2, \ldots, s_{iJ_i}^2 + \omega^2\big) + \tau^2\, \mathbf{1}\mathbf{1}^\top,$$

where $\mathbf{1}$ is a vector of ones. For a patient with two images,

$$\mathbf{V}_i = \begin{pmatrix} s_{i1}^2 + \omega^2 + \tau^2 & \tau^2 \\ \tau^2 & s_{i2}^2 + \omega^2 + \tau^2 \end{pmatrix}.$$

**Fitting the model as a GEE.** The group means are estimated with a generalised estimating equation (GEE), with the two-level model as its working model. The estimates are the precision-weighted means that a random-effects meta-analysis would give, and their standard error does not rely on the model being right. With an identity link, three descriptions give the same estimate: (a) a GEE with the two-level model as its working model, (b) generalised least squares (GLS) under the two-level model, and (c) the pooled mean of a random-effects meta-analysis with the same weights. The main text calls this the two-level model, fitted as a GEE. It is not a mixed-effects model: the variance components $\omega^2$ and $\tau^2$ are used only to set the weights, and no random effects are estimated for patients or images. The standard error comes instead from a sandwich (CR2) variance, computed from the spread between patients. A GEE needs two things: a model for the mean, here $\mathbb{E}(Y_{ij}) = \delta_{c(i)}$, and a *working* covariance for each patient, here $\mathbf{V}_i$. The two are used in different ways.

- *The working covariance sets the weights.* The estimates of $\delta_1$ and $\delta_2$ solve an equation in which each patient's images are weighted by $\mathbf{V}_i^{-1}$ (Section 4.2). Images and patients with less noise get more weight.
- *The working covariance does not set the standard error.* The variance of the estimates is not read off $\mathbf{V}_i$. It is a sandwich variance, computed from how far each patient's effects actually fall from the estimate (Section 5).

So $\mathbf{V}_i$ does not have to be right. If the clumping, the patches or the patients behave differently from the model, or $\omega^2$ and $\tau^2$ are estimated poorly, the weights are less good and some power is lost, but the standard error still reflects the real spread between patients. This is why the model can be kept simple.

#### 4.2 Estimating the group means

This section estimates the group means $\delta_1$ and $\delta_2$ as if $\omega^2$ and $\tau^2$ were known. Section 4.3 then estimates $\omega^2$ and $\tau^2$, and their estimates are used in their place.

**The estimating equation.** Patient $i$ is in one group, so the mean of every entry of $\mathbf{Y}_i$ is $\delta_{c(i)}$, and the mean of $\mathbf{Y}_i$ is $\delta_{c(i)}\mathbf{1}$. The GEE estimate of $\delta_c$ solves

$$\sum_{i:\, c(i)=c} \mathbf{1}^\top \mathbf{V}_i^{-1}\big(\mathbf{Y}_i - \delta_c \mathbf{1}\big) = 0,$$

one equation for each group. Each patient's departures from the mean, $\mathbf{Y}_i - \delta_c \mathbf{1}$, are weighted by the inverse of their working covariance and added up, and $\delta_c$ is the value at which they balance. Solving for $\delta_c$,

$$\hat{\delta}_c = \frac{\sum_{i:\, c(i)=c} \mathbf{1}^\top \mathbf{V}_i^{-1} \mathbf{Y}_i}{\sum_{i:\, c(i)=c} \mathbf{1}^\top \mathbf{V}_i^{-1} \mathbf{1}}.$$

The next result shows that this is a weighted average in two stages, with no matrix to invert.

**Weights.** Two weights appear, one for images and one for patients.

- *Image weight.* Under the model, image $j$ of patient $i$ varies around its patient's level with variance $s_{ij}^2 + \omega^2$: its random-labelling noise plus its own departure $\eta_{ij}$. Its weight is one over that variance,

  $$\gamma_{ij} \coloneqq \frac{1}{s_{ij}^2 + \omega^2}.$$

  The patient's total image weight is $\Gamma_i = \sum_{j=1}^{J_i} \gamma_{ij}$, and their weighted mean effect is $\bar{Y}_i = \frac{1}{\Gamma_i} \sum_{j=1}^{J_i} \gamma_{ij} Y_{ij}$.

- *Patient weight.* $\bar{Y}_i$ varies around $\delta_{c(i)}$ with variance $1/\Gamma_i + \tau^2$: the noise left after averaging the patient's images, plus the patient's own departure $b_i$. Its weight is again one over that variance,

  $$u_i \coloneqq \frac{1}{1/\Gamma_i + \tau^2}.$$

**Proposition 7** (Closed-form estimate)**.** *For each patient,* $\mathbf{1}^\top \mathbf{V}_i^{-1} \mathbf{1} = u_i$ *and* $\mathbf{1}^\top \mathbf{V}_i^{-1} \mathbf{Y}_i = u_i \bar{Y}_i$*. Hence*

$$\hat{\delta}_c = \frac{\sum_{i:\, c(i)=c} u_i \bar{Y}_i}{\sum_{i:\, c(i)=c} u_i}.$$

*Proof.* Write $\mathbf{D}_i \coloneqq \operatorname{diag}(1/\gamma_{i1}, \ldots, 1/\gamma_{iJ_i})$, the diagonal matrix of $s_{ij}^2 + \omega^2$. Then $\mathbf{V}_i = \mathbf{D}_i + \tau^2 \mathbf{1}\mathbf{1}^\top$: a diagonal matrix plus a multiple of $\mathbf{1}\mathbf{1}^\top$.

The Sherman–Morrison formula states that, for an invertible matrix $\mathbf{A}$ and vectors $\mathbf{a}$, $\mathbf{b}$ with $1 + \mathbf{b}^\top \mathbf{A}^{-1} \mathbf{a} \neq 0$,

$$\left(\mathbf{A} + \mathbf{a}\mathbf{b}^\top\right)^{-1} = \mathbf{A}^{-1} - \frac{\mathbf{A}^{-1}\mathbf{a}\mathbf{b}^\top \mathbf{A}^{-1}}{1 + \mathbf{b}^\top \mathbf{A}^{-1} \mathbf{a}}.$$

With $\mathbf{A} = \mathbf{D}_i$, $\mathbf{a} = \tau^2 \mathbf{1}$ and $\mathbf{b} = \mathbf{1}$,

$$\mathbf{V}_i^{-1} = \mathbf{D}_i^{-1} - \frac{\tau^2\, \mathbf{D}_i^{-1} \mathbf{1}\mathbf{1}^\top \mathbf{D}_i^{-1}}{1 + \tau^2\, \mathbf{1}^\top \mathbf{D}_i^{-1} \mathbf{1}}.$$

Here $\mathbf{D}_i^{-1} = \operatorname{diag}(\gamma_{i1}, \ldots, \gamma_{iJ_i})$, so $\mathbf{1}^\top \mathbf{D}_i^{-1} = \boldsymbol{\gamma}_i^\top$, the row of image weights $(\gamma_{i1}, \ldots, \gamma_{iJ_i})$, and $\mathbf{1}^\top \mathbf{D}_i^{-1} \mathbf{1} = \Gamma_i$. This requires $1 + \tau^2 \Gamma_i > 0$, which holds.

Multiplying $\mathbf{V}_i^{-1}$ on the left by $\mathbf{1}^\top$, and using $\mathbf{1}^\top \mathbf{D}_i^{-1} \mathbf{1} = \Gamma_i$ in the second term,

$$\mathbf{1}^\top \mathbf{V}_i^{-1} = \boldsymbol{\gamma}_i^\top - \frac{\tau^2\, \Gamma_i\, \boldsymbol{\gamma}_i^\top}{1 + \tau^2 \Gamma_i}.$$

Writing the first term over the same denominator,

$$\mathbf{1}^\top \mathbf{V}_i^{-1} = \frac{\big(1+\tau^2\Gamma_i\big) - \tau^2\Gamma_i}{1+\tau^2\Gamma_i}\,\boldsymbol{\gamma}_i^\top = \frac{\boldsymbol{\gamma}_i^\top}{1+\tau^2\Gamma_i}.$$

Multiplying on the right by $\mathbf{1}$ and by $\mathbf{Y}_i$,

$$\mathbf{1}^\top \mathbf{V}_i^{-1}\mathbf{1} = \frac{\Gamma_i}{1+\tau^2\Gamma_i} = \frac{1}{1/\Gamma_i+\tau^2} = u_i, \qquad \mathbf{1}^\top \mathbf{V}_i^{-1}\mathbf{Y}_i = \frac{\sum_j \gamma_{ij} Y_{ij}}{1+\tau^2\Gamma_i} = \frac{\Gamma_i \bar{Y}_i}{1+\tau^2\Gamma_i} = u_i \bar{Y}_i.$$

Putting these into the expression for $\hat{\delta}_c$ gives the result. □

**Two stages.** So the estimate is built in two steps.

1. *Within each patient,* the images are averaged with weights $\gamma_{ij}$, giving $\bar{Y}_i$. These weights depend on $s^2$ and $\omega^2$ but not on $\tau^2$: every image of a patient shares the same $b_i$, so $\tau^2$ does not make one image more reliable than another.

2. *Across the patients of a group,* the patient means $\bar{Y}_i$ are averaged with weights $u_i$, giving $\hat{\delta}_c$.

The second step is the same as weighted least squares with one row per patient, $\bar{Y}_i$, and weight $u_i$.

*Remark* 3 (The caps). Since $\gamma_{ij} = 1/(s_{ij}^2 + \omega^2) \le 1/\omega^2$, no image can have weight above $1/\omega^2$; likewise $u_i = 1/(1/\Gamma_i + \tau^2) \le 1/\tau^2$, so no patient can have weight above $1/\tau^2$. An image with very little labelling noise, such as one with many cells, therefore cannot carry its whole patient, and a patient with many images cannot carry their whole group: the weight stops growing once the image's or patient's own noise is small next to $\omega^2$ or $\tau^2$. The caps apply only when $\omega^2 > 0$ or $\tau^2 > 0$. When an estimate is 0 (Section 4.3), that weight is uncapped: images are weighted by $1/s^2$ alone, or patients by $\Gamma_i$ alone. For example, if every image has weight 20, a patient with 10 images ($\Gamma_i = 200$) counts 10 times as much as a patient with 1 image ($\Gamma_i = 20$) when $\tau^2 = 0$, but only 1.8 times as much when $\tau^2 = 0.05$ ($u_i = 18.2$ against 10).

#### 4.3 Estimating the variance components

**What is estimated, and what for.** The variances $\omega^2$ and $\tau^2$ are estimated in turn: $\omega^2$ from how far images spread within their patients, then $\tau^2$ from how far patients spread within their groups, given $\omega^2$. The group means then follow (Section 4.2). The two variances only set the weights, so an error in either costs power but not validity (Section 5), as long as the weight is spread over enough patients (see *When an estimate is* 0 below).

**One value for both groups.** Each variance is estimated once for each pair and radius, from the patients of both groups together, as a two-sample $t$-test pools one variance for both groups. There are two reasons.

1. *Precision.* With 6 patients per group, a separate $\tau^2$ for each group rests on 5 degrees of freedom, and its estimate typically falls anywhere between about 0.17 and 2.6 times the true value (the central 95% range of a $\chi^2_5/5$ variable). A shared $\tau^2$ rests on 10, and the range narrows to about 0.32 to 2.0. The same holds for $\omega^2$, which often has only a few patients with two or more images.

2. *The same weighting in both groups.* Separate estimates would often be 0 in one group and positive in the other. One group's patients would then be weighted by $\Gamma_i$ alone, where one large patient can dominate, and the other's with caps (Remark 3).

The cost: if a group in truth makes tissue patchier (a larger $\omega^2$) or patients more varied (a larger $\tau^2$), the shared value fits neither group exactly, and some power is lost.

**The idea.** Neither variance is seen directly, but each predicts how far the units should spread. An image differs from its patient's mean by its labelling noise, of variance $s^2_{ij}$, and by its own departure, of variance $\omega^2$. So for a trial value $t$ of $\omega^2$, divide each image's squared distance from its patient's mean by $s^2_{ij} + t$. If $t$ is the true value, each of these is about 1 on average, and their total is about the number of images less one for each patient's mean (Proposition 8). If $t$ is too small the total is too large, and if $t$ is too large it is too small. The estimate is the value of $t$ at which the total comes out right. Patients within groups are treated in the same way, with $\tau^2$ in place of $\omega^2$.

Both estimates rest on one identity, used twice, in Proposition 8 and in showing that each estimate is unique.

**Lemma 2** (The weighted mean minimises the weighted spread)**.** *Let $x_1, \dots, x_K$ have positive weights $a_1, \dots, a_K$, let $A := \sum_k a_k$ and $\bar{x} = \frac{1}{A}\sum_k a_k x_k$, and for any value $\mu$ let $f(\mu) := \sum_k a_k (x_k - \mu)^2$. Then*

$$f(\mu) = f(\bar{x}) + A(\mu - \bar{x})^2.$$

*So $f(\mu) \geq f(\bar{x})$, with equality only at $\mu = \bar{x}$: no other centre gives a smaller weighted spread.*

*Proof.* Split each distance at $\bar{x}$: $x_k - \mu = (x_k - \bar{x}) + (\bar{x} - \mu)$. Squaring,

$$(x_k - \mu)^2 = (x_k - \bar{x})^2 + 2(x_k - \bar{x})(\bar{x} - \mu) + (\bar{x} - \mu)^2.$$

Weight by $a_k$ and add. The factor $\bar{x} - \mu$ is the same for every $k$, so it comes out of the sums:

$$f(\mu) = \sum_k a_k (x_k - \bar{x})^2 + 2(\bar{x} - \mu) \sum_k a_k (x_k - \bar{x}) + (\bar{x} - \mu)^2 \sum_k a_k.$$

The middle sum is $\sum_k a_k x_k - \bar{x}A = A\bar{x} - A\bar{x} = 0$, by the definition of $\bar{x}$: the weighted distances above and below the weighted mean cancel. The last sum is $A$. This leaves $f(\mu) = f(\bar{x}) + A(\mu - \bar{x})^2$. □

**Proposition 8** (Expected weighted spread)**.** *Let $x_1, \dots, x_K$ be independent, with a common mean $\mu$ and variances $1/a_1, \dots, 1/a_K$. Let $A = \sum_k a_k$ and $\bar{x} = \frac{1}{A}\sum_k a_k x_k$. Then*

$$\mathbb{E}\Big\{\sum_{k=1}^{K} a_k \big(x_k - \bar{x}\big)^2\Big\} = K - 1.$$

*Proof.* Write $x_k = \mu + e_k$, so the $e_k$ are independent with mean 0 and $\mathbb{E}(e_k^2) = 1/a_k$, and let $\bar{e} = \frac{1}{A}\sum_k a_k e_k = \bar{x} - \mu$. Lemma 2, at the true mean $\mu$, gives $\sum_k a_k(x_k - \mu)^2 = \sum_k a_k(x_k - \bar{x})^2 + A(\mu - \bar{x})^2$, that is,

$$\sum_k a_k\big(x_k - \bar{x}\big)^2 = \sum_k a_k e_k^2 - A\bar{e}^2.$$

For the first term, $\mathbb{E}\big(\sum_k a_k e_k^2\big) = \sum_k a_k \cdot \frac{1}{a_k} = K$. For the second, $\bar{e}$ has mean 0 and variance $\frac{1}{A^2}\sum_k a_k^2 \cdot \frac{1}{a_k} = \frac{1}{A}$, so $\mathbb{E}\big(A\bar{e}^2\big) = 1$. Subtracting gives $K - 1$. □

**$\omega^2$: the spread of images within patients.** For a trial value $t$ of $\omega^2$, weight image $j$ of patient $i$ by $\gamma_{ij}(t) = 1/(s_{ij}^2 + t)$, let $\Gamma_i(t) := \sum_j \gamma_{ij}(t)$, and let $\bar{Y}_i(t) = \frac{1}{\Gamma_i(t)}\sum_j \gamma_{ij}(t)Y_{ij}$ be the patient's mean under these weights, as in Section 4.2. The within-patient spread is

$$Q(t) := \sum_i \sum_{j=1}^{J_i} \gamma_{ij}(t)\big(Y_{ij} - \bar{Y}_i(t)\big)^2.$$

Each image is compared only with its own patient's mean, so $b_i$ and $\delta_{c(i)}$ drop out, and $Q$ measures only $\eta_{ij} + \varepsilon_{ij}$.

**The expected spread of images.** Fix patient $i$ and the departure $b_i$. The images $Y_{ij} = \delta_{c(i)} + b_i + \eta_{ij} + \varepsilon_{ij}$ are then independent, with the common mean $\delta_{c(i)} + b_i$ and variances $s_{ij}^2 + \omega^2 = 1/\gamma_{ij}(\omega^2)$. Proposition 8, with $K = J_i$ and $a_j = \gamma_{ij}(\omega^2)$, gives that the patient's part of $Q(\omega^2)$ has expected value $J_i - 1$. This does not depend on $b_i$, so it also holds without fixing it. Summing over patients,

$$\mathbb{E}\big\{Q(\omega^2)\big\} = d, \qquad d := \sum_i (J_i - 1).$$

A patient with one image adds 0 to both $Q$ and $d$.

**The estimate of $\omega^2$.** $\hat{\omega}^2$ is the value $t$ at which the images spread exactly as much as they should, $Q(t) = d$.

For example, take one patient with two images, $Y = 0.10$ and $Y = 0.30$, each with noise $s^2 = 0.01$. Their mean is 0.20, each image is 0.10 from it, and

$$Q(t) = \frac{0.10^2}{0.01 + t} + \frac{0.10^2}{0.01 + t} = \frac{0.02}{0.01 + t}, \qquad d = 2 - 1 = 1.$$

At $t = 0$, $Q = 2$: the images spread more than their noise alone explains. At $t = 0.01$, $Q = 1 = d$, so $\hat{\omega}^2 = 0.01$. At $t = 0.03$, $Q = 0.5$: that guess allows for more spread than there is.

**There is at most one such value.** $Q(t)$ decreases as $t$ grows. Each term has $t$ in its denominator, but each patient's mean $\bar{Y}_i(t)$ also changes with $t$, so this needs two steps. Take $t_1 < t_2$.

1. At $t_2$, keep the old means $\bar{Y}_i(t_1)$. Each squared distance is unchanged and its denominator is larger, so the total is smaller than $Q(t_1)$.

2. Now replace them by the means $\bar{Y}_i(t_2)$. A weighted mean is the value that makes its weighted total of squared distances smallest (Lemma 2), so this can only lower the total further, to $Q(t_2)$.

So $Q(t_2) \leq Q(t_1)$, $Q$ crosses $d$ at most once, and the crossing is found numerically.

**When $\hat{\omega}^2 = 0$.** If $Q(0) \leq d$, the images spread no more than their labelling noise allows. A larger $t$ only lowers $Q$, and a variance cannot be negative, so $\hat{\omega}^2 = 0$. In the example, images at 0.18 and 0.22 give $Q(0) = 2 \times 0.02^2/0.01 = 0.08 < 1$. A patient with one image is their own mean, so they add 0 to $Q$ and $J_i - 1 = 0$ to $d$. Only patients with two or more images inform $\omega^2$; if there are none, $d = 0$ and $\hat{\omega}^2 = 0$.

**$\tau^2$: the spread of patients within groups.** Given $\hat{\omega}^2$, each patient's mean $\bar{Y}_i$ and noise $1/\Gamma_i$ are known (Section 4.2). Under the model, $\bar{Y}_i$ varies around its group mean with variance $1/\Gamma_i + \tau^2 = 1/u_i$. For a trial value $t$ of $\tau^2$, let $u_i(t) = 1/(1/\Gamma_i + t)$, let $\hat{\delta}_c(t)$ be the weighted mean of Proposition 7 with these weights, and let

$$X(t) := \sum_i u_i(t)\big(\bar{Y}_i - \hat{\delta}_{c(i)}(t)\big)^2.$$

$X(t)$ is the spread of the patients around their own group's mean, each patient weighted by how precisely their mean would be known if $\tau^2$ were $t$.

**The expected spread of patients.** Fix a group $c$ with $m_c$ patients. At the true $\tau^2$, with $\hat{\omega}^2$ taken as $\omega^2$, the patients' means $\bar{Y}_i$ are independent, with the common mean $\delta_c$ and variances $1/u_i$, and $\hat{\delta}_c$ is their mean weighted by $u_i$. Proposition 8, with $K = m_c$ and $a_i = u_i$, gives that the group's part of $X(\tau^2)$ has expected value $m_c - 1$. Summing over the two groups,

$$\mathbb{E}\big\{X(\tau^2)\big\} = m - 2,$$

where $m$ is the number of patients and 2 is the number of group means estimated.

**The estimate of $\tau^2$.** $\hat{\tau}^2$ is the value $t$ at which $X(t) = m - 2$. By the same two steps as for $Q$, with group means in place of patient means, $X(t)$ decreases as $t$ grows, so there is at most one such value, and it is found numerically. If $X(0) \leq m - 2$, the patients spread no more than their noise allows, and $\hat{\tau}^2 = 0$.

**Why this estimator.** This is the estimator of Paule & Mandel [58], which they solved iteratively, with negative values replaced by 0. It has two features that suit this setting.

1. *The weights match.* The spread is weighted exactly as the means of Section 4.2 are, with the trial value included.

2. *No distribution is assumed.* Proposition 8 needs only means and variances. Restricted maximum likelihood, the usual alternative, assumes that every term of the model is normal, whereas the effects of Section 3 are bounded and often skewed.

**When an estimate is 0.** Either estimate can be exactly 0, and then its cap (Remark 3) is lost.

- $\hat{\omega}^2$ is often 0 when few patients have two or more images, and is 0 whenever their images spread no more than labelling noise allows. Images are then weighted by $1/s_{ij}^2$ alone, so a large image can carry most of its patient.

- $\hat{\tau}^2$ is often 0 when there are few patients, and is 0 whenever they spread no more than their noise allows. Patients are then weighted by $\Gamma_i$ alone, so a patient with many cells or images can carry most of their group. With too few patients carrying the weight, the standard error is no longer reliable, so the test of Section 5.7 is not run when the weights leave too few effective patients (Section 5.8).

### 5 Comparing groups

#### 5.1 What is compared

**The difference.** The two groups are compared through

$$D := \hat{\delta}_2 - \hat{\delta}_1,$$

which is positive when the effect is larger in group 2. The test is two-sided. Different groups have different patients, so $\hat{\delta}_1$ and $\hat{\delta}_2$ are independent and $\mathrm{Var}(D) = \mathrm{Var}(\hat{\delta}_1) + \mathrm{Var}(\hat{\delta}_2)$. The rest of this section estimates each group's variance in turn.

**Each estimate is a weighted sum of patients.** Write $f_i := \bar{Y}_i - \delta_{c(i)}$ for how far patient $i$'s mean is from their group's true mean, and $U_c = \sum_{i:\, c(i)=c} u_i$. By Proposition 7, $\hat{\delta}_c = \sum_i (u_i/U_c)\bar{Y}_i$, a weighted average whose weights $u_i/U_c$ add to 1. So $\delta_c$ can be written as the same weighted average of itself, $\delta_c = \sum_i (u_i/U_c)\delta_c$, and subtracting term by term,

$$\hat{\delta}_c - \delta_c = \sum_{i:\, c(i)=c} \frac{u_i}{U_c}\bar{Y}_i - \sum_{i:\, c(i)=c} \frac{u_i}{U_c}\delta_c = \sum_{i:\, c(i)=c} \frac{u_i}{U_c}\big(\bar{Y}_i - \delta_c\big) = \sum_{i:\, c(i)=c} \frac{u_i}{U_c} f_i.$$

The estimate's error is the weighted average of the patients' own errors, with the weights used to build it. The $f_i$ of different patients are independent with mean 0, so, with the weights held fixed,

$$\mathrm{Var}(\hat{\delta}_c) = \sum_{i:\, c(i)=c} \left(\frac{u_i}{U_c}\right)^2 \mathrm{Var}(f_i). \tag{3}$$

This needs only that patients are independent. It does not need the model's variances to be right.

**The model-based variance, and why it is not used.** If the model were right, $\mathrm{Var}(f_i)$ would be $1/u_i$ (Section 4.2), and (3) would give

$$\mathrm{Var}(\hat{\delta}_c) = \sum_i \frac{u_i^2}{U_c^2} \cdot \frac{1}{u_i} = \frac{1}{U_c}.$$

This is simple, but it is right only if every variance in the model is: $s_{ij}^2$, which understates real images (Section 4.1), and $\hat{\omega}^2$ and $\hat{\tau}^2$, which are estimates. If patients in truth spread more than the model says, $1/U_c$ is too small and the test finds differences that are not there. So the test uses (3) with $\mathrm{Var}(f_i)$ estimated from how far each patient actually falls from the estimate.

#### 5.2 The sandwich estimator

Equation (3) gives the variance of $\hat{\delta}_c$ once each patient's $\mathrm{Var}(f_i)$, the average squared distance of patient $i$'s mean from the true group mean, is known. The model supplies a value, $1/u_i$, but Section 5.1 explained why it is not trusted. The sandwich estimator takes the spread from the data instead. It looks at how far each patient actually fell from the estimate, squares that distance, and uses the square where (3) has $\mathrm{Var}(f_i)$. The model still chooses the weights $u_i/U_c$; only the variance comes from the data.

One patient gives one $f_i$, not a spread, so its variance cannot be read off that patient. But a variance is the average of a square: $f_i$ has mean 0, so $\mathrm{Var}(f_i) = \mathbb{E}(f_i^2)$. The square $f_i^2$ is too large for some patients and too small for others, but right on average, and so

$$\mathbb{E}\Big\{ \sum_{i:\, c(i)=c} \Big(\frac{u_i}{U_c}\Big)^2 f_i^2 \Big\} = \sum_{i:\, c(i)=c} \Big(\frac{u_i}{U_c}\Big)^2 \mathrm{Var}(f_i) = \mathrm{Var}(\hat{\delta}_c).$$

Patients are independent, so the errors of the single squares partly cancel in the sum, and the more patients there are, the closer the sum comes to $\mathrm{Var}(\hat{\delta}_c)$.

**Plugging in the residuals.** The sum cannot be computed yet, because $f_i = \bar{Y}_i - \delta_c$ needs the unknown $\delta_c$. The residual

$$\hat{e}_i := \bar{Y}_i - \hat{\delta}_{c(i)}$$

is $f_i$ with $\delta_c$ replaced by its estimate. Putting $\hat{e}_i$ for $f_i$ in the sum gives

$$V_c^{\mathrm{CR0}} := \sum_{i:\, c(i)=c} \Big(\frac{u_i}{U_c}\Big)^2 \hat{e}_i^2.$$

This step is not exact: $\hat{e}_i$ is measured from a mean that patient $i$ helped to fit, so $\hat{e}_i^2$ runs smaller than $f_i^2$ on average. Section 5.3 measures and removes this bias.

**Why it is called a sandwich.** In the matrix form of Section 4.2,

$$V_c^{\mathrm{CR0}} = \Big(\sum_i \mathbf{1}^\top \mathbf{V}_i^{-1}\mathbf{1}\Big)^{-1}\Big\{\sum_i \big(\mathbf{1}^\top \mathbf{V}_i^{-1}\hat{\mathbf{r}}_i\big)^2\Big\}\Big(\sum_i \mathbf{1}^\top \mathbf{V}_i^{-1}\mathbf{1}\Big)^{-1}, \qquad \hat{\mathbf{r}}_i = \mathbf{Y}_i - \hat{\delta}_c\mathbf{1},$$

two copies of the model's "bread", $\sum_i \mathbf{1}^\top \mathbf{V}_i^{-1}\mathbf{1} = U_c$, around the "meat" of observed residuals. By Proposition 7, $\mathbf{1}^\top \mathbf{V}_i^{-1}\hat{\mathbf{r}}_i = u_i\bar{Y}_i - u_i\hat{\delta}_c = u_i\hat{e}_i$, so the two forms agree.

**One variance for each group.** $V_c^{\mathrm{CR0}}$ is computed separately in each group, from that group's own residuals, and the variance of $D$ is estimated by the sum of the two groups' variances. As in Welch's form of the two-sample $t$-test, the groups are not assumed to spread equally. This is the cluster-robust variance of Liang & Zeger [59], with patients as the clusters; the name CR0 follows Cameron & Miller [60] and Pustejovsky & Tipton [61].

**Weights held fixed.** The weights $u_i$ depend on $\hat{\omega}^2$ and $\hat{\tau}^2$, which are estimated from the same data. As is usual for GEEs, they are treated as fixed. This is accurate when the estimates are stable, and the effective-patients rule of Section 5.8 excludes the cases where a few patients carry the weight.

#### 5.3 The CR2 criterion

**Residuals are too small.** Each residual is measured from a mean that the patient helped to fit, so it is pulled towards 0. Let $h_i := u_i/U_c$ be patient $i$'s share of their group's weight, its *leverage*; the shares of a group add to 1.

**Proposition 9** (Expected squared residual). *If the model's variances are right, so that* $\mathrm{Var}(f_i) = 1/u_i$, *then*

$$\mathbb{E}\big(\hat{e}_i^2\big) = \frac{1-h_i}{u_i}, \qquad \textit{and} \qquad \mathbb{E}\big(V_c^{\mathrm{CR0}}\big) = \frac{1}{U_c}\Big(1 - \sum_{i:\,c(i)=c} h_i^2\Big).$$

*Proof.* Subtracting and adding $\delta_c$, and using Section 5.1 for $\hat{\delta}_c - \delta_c$,

$$\hat{e}_i = \bar{Y}_i - \hat{\delta}_c = (\bar{Y}_i - \delta_c) - (\hat{\delta}_c - \delta_c) = f_i - \sum_k \frac{u_k}{U_c} f_k = f_i - \bar{f}_c, \qquad \bar{f}_c := \sum_k h_k f_k.$$

Here $\bar{f}_c$ is the weighted average of the errors $f_k$ of all the patients in group $c$, with weights $h_k = u_k/U_c$. So patient $i$'s residual is their own error minus this average error.

Squaring, $(f_i - \bar{f}_c)^2 = f_i^2 - 2f_i\bar{f}_c + \bar{f}_c^2$, and taking the average of each piece,

$$\mathbb{E}(\hat{e}_i^2) = \mathbb{E}(f_i^2) - 2\,\mathbb{E}(f_i\bar{f}_c) + \mathbb{E}(\bar{f}_c^2).$$

Every $f_k$ has mean 0, so $\bar{f}_c$ has mean 0 too. For quantities with mean 0, the average of a square is a variance and the average of a product is a covariance: $\mathbb{E}(f_i^2) = \mathrm{Var}(f_i)$, $\mathbb{E}(f_i\bar{f}_c) = \mathrm{Cov}(f_i, \bar{f}_c)$ and $\mathbb{E}(\bar{f}_c^2) = \mathrm{Var}(\bar{f}_c)$. So

$$\mathbb{E}(\hat{e}_i^2) = \mathrm{Var}(f_i) - 2\,\mathrm{Cov}(f_i, \bar{f}_c) + \mathrm{Var}(\bar{f}_c).$$

Each term in turn, using that the $f_k$ are independent:

- $\mathrm{Var}(f_i) = 1/u_i$;
- only the $k = i$ term of $\bar{f}_c$ involves $f_i$, so $\mathrm{Cov}(f_i, \bar{f}_c) = h_i \,\mathrm{Var}(f_i) = \frac{u_i}{U_c} \cdot \frac{1}{u_i} = \frac{1}{U_c}$;
- the variance of a weighted sum of independent terms is the sum of the squared weights times the variances, so $\mathrm{Var}(\bar{f}_c) = \sum_k h_k^2 \cdot \frac{1}{u_k} = \sum_k \frac{u_k}{U_c^2} = \frac{1}{U_c}$.

So $\mathbb{E}(\hat{e}_i^2) = \frac{1}{u_i} - \frac{2}{U_c} + \frac{1}{U_c} = \frac{1}{u_i} - \frac{1}{U_c} = \frac{1-h_i}{u_i}$, the last step because $1/U_c = h_i/u_i$. Then

$$\mathbb{E}\big(V_c^{\mathrm{CR0}}\big) = \sum_i h_i^2 \cdot \frac{1-h_i}{u_i} = \frac{1}{U_c} \sum_i h_i(1-h_i) = \frac{1}{U_c}\Big(1 - \sum_i h_i^2\Big),$$

using $h_i^2/u_i = h_i/U_c$ and $\sum_i h_i = 1$. □

Even when the model is right, CR0 falls short of the variance of $\hat{\delta}_c$, which is then $1/U_c$, by the factor $1 - \sum_i h_i^2$. The quantity

$$m_c^{\mathrm{eff}} := \frac{1}{\sum_{i:\,c(i)=c} h_i^2} = \frac{U_c^2}{\sum_{i:\,c(i)=c} u_i^2}$$

is the group's number of *effective patients*: it equals the number of patients when they are weighted equally, and falls towards 1 as one patient takes most of the weight. So the factor is $1 - 1/m_c^{\mathrm{eff}}$. With 6 equally weighted patients CR0 is 5/6 of the truth on average; with 2 effective patients it is half. The shortfall is largest exactly where there are fewest patients to learn the spread from.

Bell & McCaffrey [62] proposed adjusting the residuals so that the sandwich is exactly unbiased when the working model is right. This is called CR2.

#### 5.4 Closed-form CR2

**Proposition 10** (Closed-form CR2)**.** *The CR2 variance of $\hat{\delta}_c$ is*

$$V_c = \sum_{i:\,c(i)=c} \Big(\frac{u_i}{U_c}\Big)^2 \frac{\hat{e}_i^2}{1-h_i}.$$

*If the model's variances are right, $\mathbb{E}(V_c) = 1/U_c$, which is then the variance of $\hat{\delta}_c$.*

*Proof.* By Proposition 9, $\mathbb{E}\{\hat{e}_i^2/(1-h_i)\} = 1/u_i$, so each adjusted squared residual is unbiased for $\mathrm{Var}(f_i) = 1/u_i$. Taking the average of $V_c$ term by term, $\mathbb{E}(V_c) = \sum_i h_i^2/u_i = \sum_i h_i/U_c = 1/U_c$. □

So CR2 is CR0 with each patient's squared residual divided by $1 - h_i$: a patient with a small share of the weight is barely changed, and a patient with a large share, whose residual is pulled most towards 0, is scaled up most.

Section 5.5 shows that $V_c$ is the CR2 variance of Bell & McCaffrey [62] and Pustejovsky & Tipton [61], with images as the observations and patients as the clusters.

#### 5.5 CR2 in matrix form

This subsection shows that $V_c$ of Proposition 10 is a CR2 variance in the sense of Bell & McCaffrey [62] and Pustejovsky & Tipton [61], with the images as the observations and the patients as the clusters. Fix a group with patients $i = 1, \ldots, K$. Patient $i$'s images have the working covariance of Section 4.1,

$$\mathbf{V}_i = \operatorname{diag}\big(s_{i1}^2 + \omega^2, \ldots, s_{iJ_i}^2 + \omega^2\big) + \tau^2\,\mathbf{1}\mathbf{1}^\top,$$

whose off-diagonal entries $\tau^2$ are the covariance between two images of one patient, from their shared $b_i$. The residual vector is $\hat{\mathbf{r}}_i = \mathbf{Y}_i - \hat{\hat{\delta}}_c\mathbf{1}$.

**Whitening.** Multiply patient $i$'s data by $\mathbf{V}_i^{-1/2}$, the symmetric inverse square root of $\mathbf{V}_i$. Whitened data have working covariance $\mathbf{V}_i^{-1/2}\mathbf{V}_i\mathbf{V}_i^{-1/2} = \mathbf{I}$, so the fit becomes an ordinary least-squares fit of one mean, with design rows $\tilde{\mathbf{x}}_i = \mathbf{V}_i^{-1/2}\mathbf{1}$, residuals $\tilde{\mathbf{r}}_i = \mathbf{V}_i^{-1/2}\hat{\mathbf{r}}_i$ and bread $\sum_i \tilde{\mathbf{x}}_i^\top\tilde{\mathbf{x}}_i = U_c$. Patient $i$'s block of the hat matrix is $\mathbf{H}_{ii} = \tilde{\mathbf{x}}_i\tilde{\mathbf{x}}_i^\top/U_c$.

**The adjustment.** For an ordinary least-squares fit, Bell & McCaffrey [62] multiply each cluster's residuals by the symmetric root $\mathbf{A}_i = (\mathbf{I} - \mathbf{H}_{ii})^{-1/2}$. Applied to the whitened data, this gives

$$V_c^{\mathrm{CR2}} = \frac{1}{U_c^2}\sum_{i=1}^{K}\big(\tilde{\mathbf{x}}_i^\top\mathbf{A}_i\tilde{\mathbf{r}}_i\big)^2.$$

The criterion is that of Pustejovsky & Tipton [61, eqs. 5–6], and applying the symmetric root of Bell & McCaffrey [62] after whitening is our choice.

**Other solutions.** With several images per patient, the criterion does not fix $\mathbf{A}_i$ uniquely. Pustejovsky & Tipton [61] use a different solution (their eq. 7), built from a Cholesky factor of the working covariance. The two agree when $\mathbf{V}_i$ is a multiple of the identity, as with one image per patient, and both are unbiased under the working model. When a patient's images differ in their noise the two differ slightly, so their version computed on these data gives a slightly different variance.

**Proposition 11** (Matrix CR2 equals the closed form)**.** $V_c^{\mathrm{CR2}} = V_c$ *of Proposition 10.*

*Proof.* First, $\tilde{\mathbf{x}}_i^\top\tilde{\mathbf{x}}_i = \mathbf{1}^\top\mathbf{V}_i^{-1}\mathbf{1} = u_i$ (Proposition 7). So $\mathbf{H}_{ii} = h_i\mathbf{E}_i$, where $h_i = u_i/U_c$ and $\mathbf{E}_i = \tilde{\mathbf{x}}_i\tilde{\mathbf{x}}_i^\top/u_i$ is the projection onto the direction of $\tilde{\mathbf{x}}_i$: $\mathbf{E}_i^2 = \mathbf{E}_i$ and $\mathbf{E}_i\tilde{\mathbf{x}}_i = \tilde{\mathbf{x}}_i$.

Second, a function of $\mathbf{I} - h_i\mathbf{E}_i$ acts on the direction of $\tilde{\mathbf{x}}_i$ through its value at $1 - h_i$, and on the directions at right angles to it through its value at 1. So

$$\mathbf{A}_i = (\mathbf{I} - h_i\mathbf{E}_i)^{-1/2} = \mathbf{I} + \Big(\frac{1}{\sqrt{1-h_i}} - 1\Big)\mathbf{E}_i,$$

which can be checked by squaring: with $a = (1-h_i)^{-1/2} - 1$, $(\mathbf{I} + a\mathbf{E}_i)^2 = \mathbf{I} + (2a + a^2)\mathbf{E}_i$ and $2a + a^2 = (1+a)^2 - 1 = 1/(1-h_i) - 1 = h_i/(1-h_i)$, while $(\mathbf{I} - h_i\mathbf{E}_i)^{-1} = \mathbf{I} + \frac{h_i}{1-h_i}\mathbf{E}_i$.

Third, since $\tilde{\mathbf{x}}_i^\top \mathbf{E}_i = \tilde{\mathbf{x}}_i^\top$,

$$\tilde{\mathbf{x}}_i^\top \mathbf{A}_i = \tilde{\mathbf{x}}_i^\top + \Big(\frac{1}{\sqrt{1-h_i}} - 1\Big)\tilde{\mathbf{x}}_i^\top = \frac{\tilde{\mathbf{x}}_i^\top}{\sqrt{1-h_i}}.$$

Finally, $\tilde{\mathbf{x}}_i^\top \tilde{\mathbf{r}}_i = \mathbf{1}^\top \mathbf{V}_i^{-1} \hat{\mathbf{r}}_i = u_i \hat{e}_i$ (Section 5.2). Hence

$$V_c^{\mathrm{CR2}} = \frac{1}{U_c^2} \sum_i \frac{u_i^2 \hat{e}_i^2}{1-h_i} = \sum_i \Big(\frac{u_i}{U_c}\Big)^2 \frac{\hat{e}_i^2}{1-h_i} = V_c.$$

□

Only the direction of $\tilde{\mathbf{x}}_i$ matters because a patient's images enter $\hat{\delta}_c$ only through the single number $\mathbf{1}^\top \mathbf{V}_i^{-1} \mathbf{Y}_i = u_i \bar{Y}_i$. The parts of $\tilde{\mathbf{r}}_i$ at right angles to $\tilde{\mathbf{x}}_i$, the differences between a patient's images, are multiplied by $\tilde{\mathbf{x}}_i^\top$ and vanish.

#### 5.6 Satterthwaite degrees of freedom

$V_1 + V_2$ is itself estimated from few patients, so $D/\sqrt{V_1 + V_2}$ varies more than a standard normal variable. As in Welch's test, it is compared with a $t$ distribution whose degrees of freedom $\nu$ say how many patients' worth of information $V_1 + V_2$ rests on.

**The degrees of freedom.** Satterthwaite [63] chose $\nu$ so that $\nu(V_1 + V_2)/\mathbb{E}(V_1 + V_2)$ has the mean and variance of a $\chi^2_\nu$ variable, that is,

$$\nu = \frac{2\,\mathbb{E}(V_1 + V_2)^2}{\mathrm{Var}(V_1 + V_2)}.$$

The mean and variance are computed under the working model, with the patient departures $f_i$ taken to be normal for this step only. Applied to the CR2 variance in this way, these are the degrees of freedom proposed by Bell & McCaffrey [62] and recommended by Pustejovsky & Tipton [61].

**Proposition 12** (Satterthwaite degrees of freedom)**.** *Under the working model with normal $f_i$, $\mathbb{E}(V_c) = 1/U_c$ and $\mathrm{Var}(V_c) = 2T_c$, with*

$$T_c := \frac{1}{U_c^2}\Bigg\{ \sum_{i:\, c(i)=c} \frac{h_i^2(1-2h_i)}{(1-h_i)^2} + \Big( \sum_{i:\, c(i)=c} \frac{h_i^2}{1-h_i} \Big)^2 \Bigg\}.$$

*Hence*

$$\nu = \frac{\big(1/U_1 + 1/U_2\big)^2}{T_1 + T_2}.$$

*Proof.* Fix a group and drop the subscript $c$. Under the working model with normal $f_i$, each $f_i$ is normal with mean 0 and variance $1/u_i$, so its standard deviation is $1/\sqrt{u_i}$. Dividing by it gives a standard normal variable, so define

$$z_i := \sqrt{u_i}\, f_i, \qquad \text{that is,} \qquad f_i = \frac{z_i}{\sqrt{u_i}}.$$

The $z_i$ are independent standard normal, because the $f_i$ are independent. Since $u_i = Uh_i$, $f_i = z_i/\sqrt{Uh_i}$, and so $h_k f_k = h_k z_k/\sqrt{Uh_k} = \sqrt{h_k}\, z_k/\sqrt{U}$. Putting these into $\hat{e}_i = f_i - \sum_k h_k f_k$ (Proposition 9),

$$\hat{e}_i = \frac{z_i}{\sqrt{Uh_i}} - \frac{1}{\sqrt{U}} \sum_k \sqrt{h_k}\, z_k = \frac{1}{\sqrt{Uh_i}} \Big( z_i - \sqrt{h_i} \sum_k \sqrt{h_k}\, z_k \Big) = \frac{w_i}{\sqrt{Uh_i}},$$

where $w_i$ is the term in brackets. To write all the $w_i$ at once, let $\mathbf{z} = (z_1, \ldots, z_K)^\top$ and $\mathbf{q} = (\sqrt{h_1}, \ldots, \sqrt{h_K})^\top$.

- The sum in the bracket is the same for every $i$: it is the single number $\sum_k \sqrt{h_k}\, z_k = \mathbf{q}^\top \mathbf{z}$.
- So $w_i = z_i - q_i\,(\mathbf{q}^\top \mathbf{z})$, and stacking these for $i = 1, \ldots, K$,

$$\mathbf{w} = \mathbf{z} - \mathbf{q}\,(\mathbf{q}^\top \mathbf{z}) = \mathbf{z} - (\mathbf{q}\mathbf{q}^\top)\mathbf{z} = \mathbf{P}\mathbf{z}, \qquad \mathbf{P} := \mathbf{I} - \mathbf{q}\mathbf{q}^\top.$$

- $\mathbf{q}$ has length 1, because $\mathbf{q}^\top \mathbf{q} = \sum_k h_k = 1$. Hence $\mathbf{P}$ is symmetric and $\mathbf{P}^2 = \mathbf{I} - 2\mathbf{q}\mathbf{q}^\top + \mathbf{q}(\mathbf{q}^\top \mathbf{q})\mathbf{q}^\top = \mathbf{P}$. Therefore $\mathbf{P}$ is a projection, which removes from $\mathbf{z}$ its component along $\mathbf{q}$.

Now write the CR2 variance $V = V_c$ of Proposition 10 in terms of $\mathbf{z}$. Since $u_i/U = h_i$ and $\hat{e}_i^2 = w_i^2/(Uh_i)$, each term is

$$\Big(\frac{u_i}{U}\Big)^2 \frac{\hat{e}_i^2}{1-h_i} = \frac{h_i^2}{1-h_i} \cdot \frac{w_i^2}{Uh_i} = \frac{1}{U} \cdot \frac{h_i}{1-h_i}\, w_i^2.$$

So $V = \frac{1}{U} \sum_i d_i w_i^2$ with $d_i = h_i/(1-h_i)$. A weighted sum of squares is $\sum_i d_i w_i^2 = \mathbf{w}^\top \mathbf{D} \mathbf{w}$, where $\mathbf{D} = \operatorname{diag}(d_i)$, and $\mathbf{w} = \mathbf{P}\mathbf{z}$ with $\mathbf{P}$ symmetric, so

$$V = \frac{1}{U}\, \mathbf{z}^\top \mathbf{P}\mathbf{D}\mathbf{P}\mathbf{z} = \frac{1}{U}\, \mathbf{z}^\top \mathbf{M}\mathbf{z}, \qquad \mathbf{M} := \mathbf{P}\mathbf{D}\mathbf{P}.$$

For a standard normal vector $\mathbf{z}$ and a symmetric $\mathbf{M}$, $\mathbb{E}(\mathbf{z}^\top \mathbf{M}\mathbf{z}) = \operatorname{tr}\mathbf{M}$ and $\operatorname{Var}(\mathbf{z}^\top \mathbf{M}\mathbf{z}) = 2\operatorname{tr}(\mathbf{M}^2)$. The traces use $\operatorname{tr}(\mathbf{A}\mathbf{B}) = \operatorname{tr}(\mathbf{B}\mathbf{A})$, and $\mathbf{q}^\top \mathbf{D}\mathbf{q} = \sum_i q_i^2 d_i = \sum_i h_i d_i$.

*The mean.* Moving the first $\mathbf{P}$ of $\mathbf{M} = \mathbf{P}\mathbf{D}\mathbf{P}$ to the back, and using $\mathbf{P}^2 = \mathbf{P}$,

$$\operatorname{tr}\mathbf{M} = \operatorname{tr}(\mathbf{D}\mathbf{P}\mathbf{P}) = \operatorname{tr}(\mathbf{D}\mathbf{P}) = \operatorname{tr}(\mathbf{D}) - \operatorname{tr}(\mathbf{D}\mathbf{q}\mathbf{q}^\top) = \operatorname{tr}(\mathbf{D}) - \mathbf{q}^\top \mathbf{D}\mathbf{q}.$$

With $d_i = h_i/(1-h_i)$,

$$\operatorname{tr}\mathbf{M} = \sum_i \frac{h_i}{1-h_i} - \sum_i \frac{h_i^2}{1-h_i} = \sum_i \frac{h_i(1-h_i)}{1-h_i} = \sum_i h_i = 1,$$

so $\mathbb{E}(V) = 1/U$.

*The variance.* In the same way, $\mathbf{M}^2 = \mathbf{P}\mathbf{D}\mathbf{P}\mathbf{P}\mathbf{D}\mathbf{P} = \mathbf{P}\mathbf{D}\mathbf{P}\mathbf{D}\mathbf{P}$, and moving the first $\mathbf{P}$ to the back gives $\operatorname{tr}(\mathbf{M}^2) = \operatorname{tr}(\mathbf{D}\mathbf{P}\mathbf{D}\mathbf{P})$. Multiplying out $\mathbf{D}(\mathbf{I} - \mathbf{q}\mathbf{q}^\top)\mathbf{D}(\mathbf{I} - \mathbf{q}\mathbf{q}^\top)$ gives four terms,

$$\mathbf{D}^2 - \mathbf{D}^2\mathbf{q}\mathbf{q}^\top - \mathbf{D}\mathbf{q}\mathbf{q}^\top\mathbf{D} + \mathbf{D}\mathbf{q}\,(\mathbf{q}^\top\mathbf{D}\mathbf{q})\,\mathbf{q}^\top,$$

whose traces are $\operatorname{tr}(\mathbf{D}^2)$, $-\mathbf{q}^\top\mathbf{D}^2\mathbf{q}$, $-\mathbf{q}^\top\mathbf{D}^2\mathbf{q}$ and $(\mathbf{q}^\top\mathbf{D}\mathbf{q})^2$. So

$$\begin{aligned}\operatorname{tr}(\mathbf{M}^2) &= \operatorname{tr}(\mathbf{D}^2) - 2\,\mathbf{q}^\top\mathbf{D}^2\mathbf{q} + \big(\mathbf{q}^\top\mathbf{D}\mathbf{q}\big)^2 \\ &= \sum_i \frac{h_i^2}{(1-h_i)^2} - 2\sum_i \frac{h_i^3}{(1-h_i)^2} + \Big(\sum_i \frac{h_i^2}{1-h_i}\Big)^2,\end{aligned}$$

whose first two sums combine to $\sum_i h_i^2(1-2h_i)/(1-h_i)^2$. So $\operatorname{Var}(V) = 2\operatorname{tr}(\mathbf{M}^2)/U^2 = 2T$. The two groups have different patients, so $V_1$ and $V_2$ are independent, the means and variances add, and substituting into the definition of $\nu$ gives the result. □

**What $\nu$ measures.** With $K$ equally weighted patients in one group, $h_i = 1/K$, $\mathbf{D} = \mathbf{I}/(K-1)$ and $T = 1/\{U^2(K-1)\}$, so that group alone gives $\nu = K-1$, as for a sample variance. When one patient carries most of the weight, $\nu$ falls towards 1. With two groups, $\nu$ lies between the smaller of the two groups' values and their sum, as in Welch's test. Normality is assumed only to compute $\nu$; the estimate and its standard error do not use it.

#### 5.7 The test

**The statistic.** For each pair and radius,

$$t := \frac{D}{\sqrt{V_1 + V_2}},$$

and the two-sided $p$-value is $2\Pr(T_\nu > |t|)$, where $T_\nu$ has a $t$ distribution on the $\nu$ degrees of freedom of Proposition 12.

**Many pairs.** A pair is one ordered pair of cell types. With several radii, its $p$-value is the combined one of Section 6.4. The $p$-values of the pairs that are tested are adjusted together by the method of Benjamini & Hochberg [64], which is designed to control the false discovery rate among them. Pairs that are not tested (Section 5.8) have no $p$-value and do not count towards the adjustment.

#### 5.8 When a pair is not tested

A pair is not tested, and is reported with the reason, in five cases.

1. *Fewer than two patients in a group.* A group needs at least two patients with an effect for its spread, and so its variance, to be estimated at all.
2. *No variation.* The effects show no variation at all, so that $V_1 + V_2 = 0$: for example, no $B$ cell has an $A$ cell within $r$ in any image. Then $t$ is $0/0$.
3. *No informative patient (the extra fraction).* In one group, every image of every patient is at a bound of the extra fraction, $Y = 1$ on the gain scale or $Y = -1$ on the loss scale (Section 3.3). Such effects are fixed by the bound rather than measured, so that group says nothing about how far its patients spread; its $V_c$ is then 0 or close to it, and the standard error would rest on the other group alone.

4. *Too few effective patients.* In one group, if the effective patients $m_c^{\text{eff}}$ of Section 5.3 fall below $\min(3, m_c/2)$, where $m_c$ is the number of that group's patients with an effect, then the pair is not tested. The weights $u_i$ use $\hat{\omega}^2$ and $\hat{\tau}^2$. This happens mostly when $\hat{\tau}^2 = 0$ and one patient is much larger than the rest (Section 4.3).

5. *Too few degrees of freedom.* If $\nu < 1$, the $t$ distribution is too heavy-tailed for the test to be calibrated.

**Why the effective-patients rule.** When a few patients carry the weight, the steps of this section fail together. CR0 loses a large part of the variance (Section 5.3). CR2 restores it on average by dividing by $1 - h_i$, which is close to 0 for the dominant patient, so $V_c$ rests almost entirely on that one patient's residual. And $\nu$ falls towards 1, where the $t$ distribution is very sensitive to how well the working model holds. In simulations without this rule, null tests in which one patient carried more than 90% of a group's weight gave $p < 0.05$ in 12 to 21% of cases. Floors on $\hat{\tau}^2$, caps on each patient's weight and other variance estimators were compared with the rule, and none was both calibrated and more powerful. The rule costs little power, because the pairs it removes are almost all pairs with no true difference, most of them involving rare cell types.

### 6 Extensions

#### 6.1 Adding covariates

The groups may differ in more than their condition. The patients of one group may be older, for example, or their images may hold more $A$ cells by chance. If such a variable also moves the effect, the groups' effects can differ because of it, and that difference would be mistaken for a change in co-localisation. Suppose that in every patient $B$ cells sit nearer $A$ cells in younger tissue, and that the patients of group 2 are younger, with mean age 50 against 65 in group 1. Then $\hat{\delta}_2 - \hat{\delta}_1$ is positive even if, at any given age, the two groups do not differ. A variable added to the model so that the groups are compared at the same value of it is a *covariate.* This section adds covariates to the model of Section 4.1: Section 6.1.1 gives the model, Section 6.1.2 treats covariates that differ between a patient's images, Section 6.1.3 estimates $\omega^2$ and $\tau^2$, and Section 6.1.4 gives the test. Section 6.1.5 applies the method to the abundance of the $A$ cells.

##### 6.1.1 The model with covariates

Each image $(i, j)$ has a vector $\mathbf{x}_{ij}$ of covariate values, and its effect is modelled as

$$Y_{ij} = \delta_{c(i)} + \mathbf{x}_{ij}^\top \boldsymbol{\beta} + b_i + \eta_{ij} + \varepsilon_{ij},$$

with $b_i$, $\eta_{ij}$ and $\varepsilon_{ij}$ as in Section 4.1. Each entry of $\boldsymbol{\beta}$ is the change in the effect for a change of one unit in its covariate, the others held fixed. Section 6.1.2 adds one more term for covariates that differ between a patient's images.

A covariate is of one of two kinds.

- *Patient-level:* it has the same value in every image of a patient, such as age, sex or treatment.
- *Image-level:* it can differ between the images of one patient, such as the region imaged (the tumour core or its edge), the number of cells, or the share of $A$ cells. A covariate is image-level if its value differs between two images of at least one patient.

Each covariate is centred: its mean over the images used, from both groups together, is subtracted from every value, so that a value of 0 is the average. For example, if the images used have a mean age of 58, an image from a patient aged 50 has the value $-8$. Then $\delta_c$ is the mean effect of group $c$ at the average value of every covariate. Centring does not change the comparison: it moves $\delta_1$ and $\delta_2$ by the same amount, so their difference is the same as without centring.

An image with a missing value of any covariate is removed for the pair. For the extra fraction, the scale (gain or loss) has already been chosen from every image with a group (Section 3.3), so removing images does not change it. The images left are used in every later step: the estimates of $\omega^2$ and $\tau^2$, the fit and the test. A missing patient-level covariate removes every image of that patient, and, as in Section 4, a patient none of whose images is left takes no part.

##### 6.1.2 Patient-level and image-level covariates

A patient-level covariate enters the model with one slope, as above. An image-level covariate can be related to the effect in two ways.

- *Within a patient:* images of the same patient with different values of the covariate have different effects.
- *Between patients:* patients whose images have higher values on average have different effects.

The two need not agree. Within a patient, the slope compares patches of the same tissue, so everything about the patient is held fixed and only the patch changes. Between patients, the slope compares different patients, so it also carries whatever else about the patients goes with their average value of the covariate. A single slope $\boldsymbol{\beta}$ mixes the two relations, each weighted by how much information the data hold on it.

The mixed slope can move the comparison of the groups. Suppose that the effect rises by 0.02 per unit of the covariate within a patient and by 0.05 per unit between patients, and that the patients of group 2 are 1 unit higher on average. The covariate then explains 0.05 of the difference between the groups. A single slope lies somewhere between 0.02 and 0.05; at 0.03, it removes only 0.03, and the other 0.02 is reported as a difference between the groups although the covariate explains it. The more images patients have, the more the within-patient relation counts in the single slope, and the larger this leftover becomes.

To separate the two relations, each image-level covariate gets a second column: its plain average over the patient's images. Write $\mathbf{x}^{\mathrm{I}}_{ij}$ for the image-level covariates of image $(i, j)$ and

$$\bar{\mathbf{x}}^{\mathrm{I}}_i := \frac{1}{J_i}\sum_{j=1}^{J_i} \mathbf{x}^{\mathrm{I}}_{ij}$$

for their averages over patient $i$'s images. Unlike $\bar{Y}_i$ (Section 4.2), this average is not weighted, so it is fixed before $\omega^2$ or $\tau^2$ is estimated. The model becomes

$$Y_{ij} = \delta_{c(i)} + \mathbf{x}_{ij}^\top \boldsymbol{\beta} + \left(\bar{\mathbf{x}}_i^{\mathrm{I}}\right)^\top \boldsymbol{\lambda} + b_i + \eta_{ij} + \varepsilon_{ij}.$$

For an image-level covariate, its two terms can be rewritten as

$$\beta\, x_{ij} + \lambda\, \bar{x}_i = \beta\, (x_{ij} - \bar{x}_i) + (\beta + \lambda)\, \bar{x}_i.$$

The first term compares the images of one patient, which share $\bar{x}_i$, so $\beta$ is the *within-patient slope*. The second compares patients through their averages, so $\beta + \lambda$ is the *between-patient slope*. The groups are then compared at equal patient averages of the covariate. In the example above, $\beta = 0.02$ and $\beta + \lambda = 0.05$, and the whole 0.05 is removed. For a patient with two images, with values 10% and 30%, the average 20% places the patient among the patients, and the 10% and 30% inform the slope within the patient. A patient with one image has $x_{i1} = \bar{x}_i$, so they inform only the between-patient slope.

The reported effect of an image-level covariate is $\beta$, its within-patient slope. A patient-level covariate gets no second column: its average over the patient's images is the covariate itself, so the column would repeat it.

Even with the patient average included, adjustment can still leave part of an image-level covariate's effect in the difference between the groups. The patient average $\bar{x}_i$ is itself an estimate: it averages only $J_i$ patches, so by chance it is off from the patient's typical value of the covariate. When the effect depends on that typical value, this chance part spreads the averages without moving the effect, and the fitted between-patient slope comes out flatter than the true one. Adjustment then removes only part of the difference between the groups that the covariate causes, and the rest stays in $\delta_2 - \delta_1$. The part left is larger when images of the same patient differ a lot on the covariate, when patients have few images, and when the groups differ in their typical value of the covariate. This is a known limitation of adjusting with observed patient averages [65]. The within-patient slope $\boldsymbol{\beta}$ is unaffected.

Write $\mathbf{x}_i^{\mathrm{P}}$ for the patient-level covariates of patient $i$, with slopes $\boldsymbol{\beta}^{\mathrm{P}}$, and $\boldsymbol{\beta}^{\mathrm{I}}$ for the slopes of the image-level covariates, so that $\mathbf{x}_{ij}^\top \boldsymbol{\beta} = (\mathbf{x}_i^{\mathrm{P}})^\top \boldsymbol{\beta}^{\mathrm{P}} + (\mathbf{x}_{ij}^{\mathrm{I}})^\top \boldsymbol{\beta}^{\mathrm{I}}$. Four terms of the model are then the same for all of patient $i$'s images. Together they form the patient's level

$$\alpha_i := \delta_{c(i)} + \left(\mathbf{x}_i^{\mathrm{P}}\right)^\top \boldsymbol{\beta}^{\mathrm{P}} + \left(\bar{\mathbf{x}}_i^{\mathrm{I}}\right)^\top \boldsymbol{\lambda} + b_i,$$

and the model reads

$$Y_{ij} = \alpha_i + \left(\mathbf{x}_{ij}^{\mathrm{I}}\right)^\top \boldsymbol{\beta}^{\mathrm{I}} + \eta_{ij} + \varepsilon_{ij}.$$

Within a patient, only the image-level covariates, $\eta_{ij}$ and $\varepsilon_{ij}$ change. Section 6.1.3 uses this to estimate $\omega^2$.

##### 6.1.3 Estimating the variance components

As in Section 4.3, $\omega^2$ is estimated from how far images spread within their patients, then $\tau^2$ from how far patients spread within their groups, each by the method of Paule & Mandel [58]. Covariates change both spreads. An image-level covariate makes the images of one patient differ for a reason the model already includes; if $\omega^2$ were estimated as in Section 4.3, the spread the covariate explains would be counted as $\omega^2$. The same holds for $\tau^2$ and the spread of patients. So each spread is measured after the covariates are fitted. Both estimates rest on one result, which extends Proposition 8 from one common mean to a mean that follows a set of columns.

**Proposition 13** (Expected weighted spread about a fit)**.** *Let $y_1, \ldots, y_K$ be independent, with means $\mathbf{w}_1^\top\boldsymbol{\theta}, \ldots, \mathbf{w}_K^\top\boldsymbol{\theta}$ and variances $1/a_1, \ldots, 1/a_K$, where the $\mathbf{w}_k$ are known and $\boldsymbol{\theta}$ is not. Let $\mathbf{W}$ be the matrix with rows $\mathbf{w}_k^\top$, and*

$$S := \min_{\boldsymbol{\theta}} \sum_{k=1}^{K} a_k\big(y_k - \mathbf{w}_k^\top\boldsymbol{\theta}\big)^2.$$

*Then $\mathbb{E}(S) = K - \operatorname{rank}(\mathbf{W})$.*

*Proof.* Multiply each $y_k$ and $\mathbf{w}_k$ by $\sqrt{a_k}$: let $\tilde{y}_k := \sqrt{a_k}\, y_k$ and $\tilde{\mathbf{w}}_k = \sqrt{a_k}\,\mathbf{w}_k$, and let $\tilde{\mathbf{y}}$ and $\tilde{\mathbf{W}}$ collect them. Then $a_k(y_k - \mathbf{w}_k^\top\boldsymbol{\theta})^2 = (\tilde{y}_k - \tilde{\mathbf{w}}_k^\top\boldsymbol{\theta})^2$, so

$$S = \min_{\boldsymbol{\theta}} \big\|\tilde{\mathbf{y}} - \tilde{\mathbf{W}}\boldsymbol{\theta}\big\|^2.$$

The vectors $\tilde{\mathbf{W}}\boldsymbol{\theta}$ make up the column space of $\tilde{\mathbf{W}}$, and the closest of them to $\tilde{\mathbf{y}}$ is its projection $\mathbf{H}\tilde{\mathbf{y}}$ onto that space. So $S = \|(\mathbf{I} - \mathbf{H})\tilde{\mathbf{y}}\|^2$. Multiplying rows by positive numbers does not change the rank, so the column space has dimension $\rho = \operatorname{rank}(\mathbf{W})$.

Write $\tilde{\mathbf{y}} = \tilde{\mathbf{W}}\boldsymbol{\theta} + \mathbf{e}$. The $e_k = \sqrt{a_k}(y_k - \mathbf{w}_k^\top\boldsymbol{\theta})$ are independent, with mean 0 and variance $a_k \cdot (1/a_k) = 1$. The true mean $\tilde{\mathbf{W}}\boldsymbol{\theta}$ lies in the column space, so $(\mathbf{I} - \mathbf{H})\tilde{\mathbf{W}}\boldsymbol{\theta} = \mathbf{0}$, and

$$S = \big\|(\mathbf{I} - \mathbf{H})\mathbf{e}\big\|^2 = \mathbf{e}^\top(\mathbf{I} - \mathbf{H})\mathbf{e},$$

since $\mathbf{I} - \mathbf{H}$ is symmetric and $(\mathbf{I} - \mathbf{H})^2 = \mathbf{I} - \mathbf{H}$. For any matrix $\mathbf{M}$, $\mathbf{e}^\top\mathbf{M}\mathbf{e} = \sum_k M_{kk}e_k^2 + \sum_{k \neq k'} M_{kk'}e_k e_{k'}$. Taking expectations, $\mathbb{E}(e_k^2) = 1$ and $\mathbb{E}(e_k e_{k'}) = 0$ for $k \neq k'$, so $\mathbb{E}(\mathbf{e}^\top\mathbf{M}\mathbf{e}) = \operatorname{tr}\mathbf{M}$. Hence $\mathbb{E}(S) = K - \operatorname{tr}\mathbf{H}$. A projection onto a space of dimension $\rho$ has $\rho$ eigenvalues equal to 1 and the rest equal to 0, so $\operatorname{tr}\mathbf{H} = \rho$. □

With one column of ones, $\mathbf{w}_k = 1$, the minimum is at the weighted mean (Lemma 2), $\operatorname{rank}(\mathbf{W}) = 1$, and this is Proposition 8. Each column of $\mathbf{W}$ costs one from the expected spread, unless it is a combination of the other columns.

For $\omega^2$, take a trial value $t$, weight image $j$ of patient $i$ by $\gamma_{ij}(t) = 1/(s_{ij}^2 + t)$ as in Section 4.3, and let

$$Q_{\mathrm{W}}(t) := \min_{\alpha_i,\, \boldsymbol{\beta}^{\mathrm{I}}} \sum_{i:\, J_i \geq 2} \sum_{j=1}^{J_i} \gamma_{ij}(t)\Big(Y_{ij} - \alpha_i - \big(\mathbf{x}_{ij}^{\mathrm{I}}\big)^\top \boldsymbol{\beta}^{\mathrm{I}}\Big)^2,$$

over the patients with two or more images. Here $\alpha_i$ is the patient's level of Section 6.1.2, fitted as a separate unknown for each patient without splitting it into its parts, so only the image-level covariates remain. With no image-level covariate, the minimum over $\alpha_i$ is at $\bar{Y}_i(t)$ and $Q_{\mathrm{W}}(t)$ is the $Q(t)$ of Section 4.3.

Fix every $b_i$. Then image $j$ of patient $i$ has mean $\alpha_i + (\mathbf{x}_{ij}^{\mathrm{I}})^\top \boldsymbol{\beta}^{\mathrm{I}}$ and variance $1/\gamma_{ij}(\omega^2)$, and different images are independent. This is the setting of Proposition 13 at $t = \omega^2$, so $\mathbb{E}\{Q_{\mathrm{W}}(\omega^2)\}$ is the number of observations minus the rank of the design. We count each in turn.

The observations are the images of the patients with two or more images, $\sum_{i:\, J_i \geq 2} J_i$ of them.

The design has one column for each of these patients, marking that patient's images, and one column for each image-level covariate. Each patient's column adds one to the rank. An image-level covariate adds one more only if it still varies once each patient's average is subtracted, and is not a combination of the other image-level covariates. Write $r_{\mathrm{W}}$ for the number that do; with one image-level covariate that differs between a patient's images, $r_{\mathrm{W}} = 1$.

Subtracting, each patient with two or more images contributes $J_i - 1$, so

$$\mathbb{E}\big\{Q_{\mathrm{W}}(\omega^2)\big\} = d - r_{\mathrm{W}}, \qquad d = \sum_i (J_i - 1),$$

where the sum for $d$ can run over all patients, because a patient with one image has $J_i - 1 = 0$. $\hat{\omega}^2$ is the value $t$ at which $Q_{\mathrm{W}}(t) = d - r_{\mathrm{W}}$, and $\hat{\omega}^2 = 0$ if $Q_{\mathrm{W}}(0) \leq d - r_{\mathrm{W}}$. This target needs only that, within a patient, the effect changes linearly with the image-level covariates. How patients' levels relate to the covariates does not enter, because each $\alpha_i$ is free.

For $\tau^2$, take the patient summaries at $\hat{\omega}^2$: $\Gamma_i$ and $\bar{Y}_i$ as in Section 4.2, and the same weighted average of the rows of the design,

$$\bar{\mathbf{z}}_i := \frac{1}{\Gamma_i} \sum_{j=1}^{J_i} \gamma_{ij} \mathbf{z}_{ij},$$

where $\mathbf{z}_{ij}$ is image $(i, j)$'s row of the design,

$$\mathbf{z}_{ij} := \Big(1\{c(i) = 1\},\ 1\{c(i) = 2\},\ \mathbf{x}_{ij}^\top,\ \big(\bar{\mathbf{x}}_i^{\mathrm{I}}\big)^\top\Big)^\top,$$

and $1\{c(i) = c\}$ is 1 if patient $i$ is in group $c$ and 0 otherwise. With the coefficients in $\boldsymbol{\theta} = (\delta_1, \delta_2, \boldsymbol{\beta}^\top, \boldsymbol{\lambda}^\top)^\top$, $\mathbf{z}_{ij}^\top \boldsymbol{\theta} = \delta_{c(i)} + \mathbf{x}_{ij}^\top \boldsymbol{\beta} + (\bar{\mathbf{x}}_i^{\mathrm{I}})^\top \boldsymbol{\lambda}$, which is $\mathbb{E}(Y_{ij})$. For a trial value $t$ of $\tau^2$, with $u_i(t) = 1/(1/\Gamma_i + t)$, let

$$X_{\mathrm{B}}(t) := \min_{\boldsymbol{\theta}} \sum_{i=1}^{m} u_i(t) \big(\bar{Y}_i - \bar{\mathbf{z}}_i^\top \boldsymbol{\theta}\big)^2.$$

Under the model, $\bar{Y}_i$ has mean $\bar{\mathbf{z}}_i^\top \boldsymbol{\theta}$ and variance $1/\Gamma_i + \tau^2 = 1/u_i$, independently across patients. Proposition 13, with $K = m$ and $\mathbf{W}$ the matrix $\bar{\mathbf{Z}}$ of rows $\bar{\mathbf{z}}_i^\top$, gives

$$\mathbb{E}\big\{X_{\mathrm{B}}(\tau^2)\big\} = m - \operatorname{rank}(\bar{\mathbf{Z}}).$$

$\hat{\tau}^2$ is the value $t$ at which $X_{\mathrm{B}}(t) = m - \operatorname{rank}(\bar{\mathbf{Z}})$, and $\hat{\tau}^2 = 0$ if $X_{\mathrm{B}}(0) \leq m - \operatorname{rank}(\bar{\mathbf{Z}})$. The rank is usually the number of columns, and smaller when a column is a combination of the others. With no covariates, $\bar{\mathbf{z}}_i$ is patient $i$'s group indicator, the rank is 2, the fit is each group's weighted mean, and this is the $X(t) = m - 2$ of Section 4.3.

Both $Q_{\mathrm{W}}(t)$ and $X_{\mathrm{B}}(t)$ decrease in $t$ by the argument of Section 4.3, so each crosses its target at most once, and the crossing is found numerically.

##### 6.1.4 The test

Collect patient $i$'s rows $\mathbf{z}_{ij}^\top$ in the matrix $\mathbf{Z}_i$. With $\mathbf{V}_i$ of Section 4.1 at $\hat\omega^2$ and $\hat\tau^2$, the estimate is the generalised least-squares fit

$$\hat{\boldsymbol\theta} := \Big(\sum_{i=1}^m \mathbf{Z}_i^\top \mathbf{V}_i^{-1}\mathbf{Z}_i\Big)^{-1}\sum_{i=1}^m \mathbf{Z}_i^\top \mathbf{V}_i^{-1}\mathbf{Y}_i.$$

With only the group columns, $\mathbf{Z}_i$ is a column of ones in its group's place, and this is Proposition 7 in each group. The difference between the groups is $D = \hat\delta_2 - \hat\delta_1 = \mathbf{c}^\top\hat{\boldsymbol\theta}$, where $\mathbf{c}$ has $-1$ in the place of $\delta_1$, 1 in the place of $\delta_2$ and 0 elsewhere.

The slopes are shared by the two groups, so both $\hat\delta_1$ and $\hat\delta_2$ depend on every patient. They are no longer independent, and the sum $V_1 + V_2$ of Section 5 does not apply. The variance of $D$ is instead the CR2 variance of the whole fit, in the matrix form of Section 5.5. Whiten each patient's data: let $\mathbf{X}_i := \mathbf{V}_i^{-1/2}\mathbf{Z}_i$ and $\tilde{\mathbf{r}}_i = \mathbf{V}_i^{-1/2}(\mathbf{Y}_i - \mathbf{Z}_i\hat{\boldsymbol\theta})$, let $\mathbf{B} = (\sum_i \mathbf{X}_i^\top\mathbf{X}_i)^{-1}$ be the bread, and let $\mathbf{A}_i := (\mathbf{I} - \mathbf{X}_i\mathbf{B}\mathbf{X}_i^\top)^{-1/2}$ be the adjustment of Bell & McCaffrey [62]. Then

$$V^{\mathrm{CR2}} = \sum_{i=1}^m \big(\mathbf{a}_i^\top\tilde{\mathbf{r}}_i\big)^2, \qquad \mathbf{a}_i := \mathbf{A}_i\mathbf{X}_i\mathbf{B}\mathbf{c}.$$

With only the group columns, $\mathbf{B}$ is diagonal with entries $1/U_1$ and $1/U_2$, a patient's terms involve only their own group, and $V^{\mathrm{CR2}} = V_1 + V_2$ by Proposition 11.

**Proposition 14** (Satterthwaite degrees of freedom of the whole fit)**.** *Let $\boldsymbol\Omega$ be the $m\times m$ matrix with entries*

$$\Omega_{ik} := \mathbf{a}_i^\top\big(1\{i=k\}\,\mathbf{I} - \mathbf{X}_i\mathbf{B}\mathbf{X}_k^\top\big)\mathbf{a}_k.$$

*Under the working model with normal errors,* $\mathbb{E}(V^{\mathrm{CR2}}) = \operatorname{tr}\boldsymbol\Omega$ *and* $\operatorname{Var}(V^{\mathrm{CR2}}) = 2\operatorname{tr}(\boldsymbol\Omega^2)$*, so*

$$\nu = \frac{\big(\operatorname{tr}\boldsymbol\Omega\big)^2}{\operatorname{tr}\big(\boldsymbol\Omega^2\big)}.$$

*Proof.* Stack the whitened data of all patients into $\tilde{\mathbf{y}}$, $\mathbf{X}$ and $\tilde{\mathbf{r}}$. Under the working model, $\tilde{\mathbf{y}} = \mathbf{X}\boldsymbol\theta + \mathbf{e}$ with $\mathbf{e}$ standard normal. The fit is the least-squares fit of the whitened data, so $\tilde{\mathbf{r}} = (\mathbf{I}-\mathbf{H})\mathbf{e}$, where $\mathbf{H} = \mathbf{X}\mathbf{B}\mathbf{X}^\top$ is the hat matrix, with blocks $\mathbf{H}_{ik} = \mathbf{X}_i\mathbf{B}\mathbf{X}_k^\top$; as in the proof of Proposition 13, $(\mathbf{I}-\mathbf{H})\mathbf{X} = \mathbf{0}$ removes $\boldsymbol\theta$.

Let $\mathbf{g}_i$ be the stacked vector with $\mathbf{a}_i$ in patient $i$'s block and $\mathbf{0}$ elsewhere, and $\mathbf{v}_i = (\mathbf{I}-\mathbf{H})\mathbf{g}_i$. Since $\mathbf{I}-\mathbf{H}$ is symmetric, $\mathbf{a}_i^\top\tilde{\mathbf{r}}_i = \mathbf{g}_i^\top(\mathbf{I}-\mathbf{H})\mathbf{e} = \mathbf{v}_i^\top\mathbf{e}$, and

$$V^{\mathrm{CR2}} = \sum_i \big(\mathbf{v}_i^\top\mathbf{e}\big)^2 = \mathbf{e}^\top\mathbf{M}\mathbf{e}, \qquad \mathbf{M} := \sum_i \mathbf{v}_i\mathbf{v}_i^\top.$$

Since $(\mathbf{I}-\mathbf{H})^2 = \mathbf{I}-\mathbf{H}$, $\mathbf{v}_i^\top\mathbf{v}_k = \mathbf{g}_i^\top(\mathbf{I}-\mathbf{H})\mathbf{g}_k = \mathbf{a}_i^\top(1\{i=k\}\,\mathbf{I} - \mathbf{H}_{ik})\mathbf{a}_k = \Omega_{ik}$.

For a standard normal vector $\mathbf{e}$ and a symmetric $\mathbf{M}$, $\mathbb{E}(\mathbf{e}^\top\mathbf{M}\mathbf{e}) = \operatorname{tr}\mathbf{M}$ and $\operatorname{Var}(\mathbf{e}^\top\mathbf{M}\mathbf{e}) = 2\operatorname{tr}(\mathbf{M}^2)$ (Section 5.6). Here $\operatorname{tr}\mathbf{M} = \sum_i \mathbf{v}_i^\top\mathbf{v}_i = \operatorname{tr}\boldsymbol\Omega$, and, by the cyclic property,

$$\operatorname{tr}(\mathbf{M}^2) = \sum_{i,k}\operatorname{tr}\big(\mathbf{v}_i\mathbf{v}_i^\top\mathbf{v}_k\mathbf{v}_k^\top\big) = \sum_{i,k}\big(\mathbf{v}_i^\top\mathbf{v}_k\big)^2 = \sum_{i,k}\Omega_{ik}^2 = \operatorname{tr}\big(\boldsymbol\Omega^2\big),$$

since $\boldsymbol\Omega$ is symmetric. Putting the mean and variance into the definition of $\nu$ of Section 5.6 gives the result. □

With only the group columns, $\Omega_{ik} = 0$ for patients in different groups, $\mathbf{\Omega}$ splits into one block for each group, and $\nu$ is that of Proposition 12. These are the degrees of freedom of Bell & McCaffrey [62] for a general design, as recommended by Pustejovsky & Tipton [61].

The test is $t = D/\sqrt{V^{\mathrm{CR2}}}$, with the two-sided $p$-value $2\Pr(T_\nu > |t|)$. The effect of each covariate is tested in the same way, with $\mathbf{c}$ picking out its entry of $\boldsymbol{\beta}$; for an image-level covariate this is its within-patient slope (Section 6.1.2). The entries of $\boldsymbol{\lambda}$ are not reported.

The five cases of Section 5.8 are checked first, on the fit without covariates. If one holds, the pair is not tested. The covariates add one more case.

6. *The design cannot be fitted.* If there are no more patients than columns of the design, or one column is a combination of the others, $\boldsymbol{\theta}$ cannot be estimated. An example is a covariate that takes one value in every patient of group 1 and another in every patient of group 2: it cannot be told apart from the group. The pair is then tested without the covariates, as in Section 5, and reported with the reason.

##### 6.1.5 Example: abundance adjustment

How common the $A$ cells are can differ between groups: a disease may bring more of them, for example. Under label shift and hub recruitment, the expected extra fraction does not depend on how much of the tissue is near $A$ cells (Section 3.7), and so not on how common they are. However, under other models, such as one in which being near $A$ raises the odds of a candidate being a $B$ cell, the expected extra fraction would change. Then a difference in the extra fraction between groups can come from a difference in how common the $A$ cells are. Adding their share as a covariate can account for it and asks whether the difference remains when patients with the same share of $A$ cells are compared.

The covariate of image $(i, j)$ is the log share of $A$ cells among all its cells,

$$\log \frac{n_{ij}^A}{n_{ij}}.$$

The share is on the log scale because shares of a cell type often differ several-fold between images, and on this scale a doubling is the same step, whether from 1% to 2% or from 10% to 20%. Every image used has at least one $A$ cell: an image with none is not used for the pair (Section 3.5, case 3), so the logarithm is finite. The covariate belongs to the pair, through $A$, and is image-level, so it gets the patient-average column of Section 6.1.2; its reported effect is how the effect changes between a patient's images with the share of $A$ cells. It is not added when the share is the same in every image used. The adjustment is an option and is not used by default.

For example, suppose the images of group 2 have twice the share of $A$ cells of those of group 1, a difference of $\log 2 \approx 0.69$ on the log scale, and that the effect rises by 0.05 per unit of the log share, both within and between patients. The share then explains $0.69 \times 0.05 \approx 0.035$ of the difference between the groups: a difference of 0.10 without adjustment becomes 0.065 with it.

The adjustment has two limits.

- The share counts the $A$ cells of the whole image, not how they are laid out. An image whose $A$ cells are packed into nests and one whose $A$ cells are spread out can have the same share, so the adjustment does not separate more $A$ cells from more tightly packed ones.

- When more $A$ cells is itself part of the difference being studied, adjusting removes that part. If a disease brings both $A$ cells and $B$ cells to them, the adjusted test asks whether $B$ cells sit nearer $A$ cells at the same share, which is a different question.

#### 6.2 More than two groups

Some studies compare more than two groups of patients, such as three stages of a disease. With $C$ groups, the model of Section 4.1 has a mean $\delta_c$ for each group $c = 1, \dots, C$, and the question for a pair is whether all $C$ means are equal. Comparing every two groups would give $C(C-1)/2$ $p$-values per pair, three for three groups. Controlling the false discovery rate over pairs needs one $p$-value per pair, so a rule would be needed to choose among them. Instead, each pair gets one overall test of

$$H_0: \ \delta_1 = \delta_2 = \cdots = \delta_C,$$

which enters the Benjamini–Hochberg adjustment as the pair's one $p$-value. Only pairs called by the overall test are followed up with comparisons of two groups.

##### 6.2.1 Fitting the model

The fit is that of Section 6.1 with one indicator per group, $\mathbf{z}_{ij} = \big(1\{c(i) = 1\}, \dots, 1\{c(i) = C\}\big)^\top$, followed by any covariates. $\omega^2$ is estimated as before, and $\tau^2$ by Paule–Mandel on the patient summaries (Section 6.1.3), whose target is now $m - C$: one less for each group mean fitted, as the target is $m - 2$ for two groups (Section 4.3). Without covariates the groups share only $\omega^2$ and $\tau^2$, so each $\hat{\delta}_c$ is its group's weighted mean (Proposition 7) with CR2 variance $V_c$ (Proposition 10), and the estimates of different groups are independent.

##### 6.2.2 The overall test

Write $H_0$ as $\mathbf{L}\boldsymbol{\theta} = \mathbf{0}$, where each of the $g = C - 1$ rows of $\mathbf{L}$ compares one group with group 1, and has 0 in the columns of any covariates. For three groups,

$$\mathbf{L}\boldsymbol{\theta} = \begin{pmatrix} \delta_2 - \delta_1 \\ \delta_3 - \delta_1 \end{pmatrix}.$$

The CR2 variance of Section 6.1.4 carries over from one contrast to $g$ of them. Patient $i$'s term is the vector $\mathbf{s}_i = \mathbf{L}\mathbf{B}\mathbf{X}_i^\top \mathbf{A}_i \tilde{\mathbf{r}}_i$, and

$$\mathbf{V}^{\mathrm{CR2}} := \sum_{i=1}^{m} \mathbf{s}_i \mathbf{s}_i^\top$$

is the $g \times g$ CR2 covariance of $\mathbf{L}\hat{\boldsymbol{\theta}}$. With one row it is the $V^{\mathrm{CR2}}$ of Section 6.1.4; without covariates it is $\mathbf{L}\,\mathrm{diag}(V_1, \dots, V_C)\mathbf{L}^\top$. The Wald statistic

$$Q_{\mathrm{F}} := \big(\mathbf{L}\hat{\boldsymbol{\theta}}\big)^\top \big(\mathbf{V}^{\mathrm{CR2}}\big)^{-1} \mathbf{L}\hat{\boldsymbol{\theta}}$$

measures how far the group means are from equal, each direction scaled by its uncertainty; with $g = 1$, $Q_{\mathrm{F}} = t^2$. $Q_{\mathrm{F}}$ does not depend on which group is the reference: another choice of rows giving the same set of differences is $\mathbf{ML}$ for an invertible $\mathbf{M}$, and $\mathbf{M}$ cancels in $Q_{\mathrm{F}}$.

For example, take three groups with

$$\hat{\boldsymbol{\delta}} = (0.10,\ 0.15,\ 0.02), \qquad (V_1, V_2, V_3) = (0.0004,\ 0.0009,\ 0.0004).$$

The two-group statistics are $t = 1.39$ for groups 1 and 2, $-2.83$ for groups 1 and 3, and $-3.61$ for groups 2 and 3. With $\mathbf{L}$ as above, $\mathbf{L}\hat{\boldsymbol{\delta}} = (0.05,\ -0.08)$,

$$\mathbf{V}^{\mathrm{CR2}} = \begin{pmatrix} V_1 + V_2 & V_1 \\ V_1 & V_1 + V_3 \end{pmatrix} = \begin{pmatrix} 0.0013 & 0.0004 \\ 0.0004 & 0.0008 \end{pmatrix},$$

and $Q_{\mathrm{F}} = 15.4$. Comparing group 2 with group 1 and group 3 with group 2 instead gives the same 15.4.

If $\mathbf{V}^{\mathrm{CR2}}$ were the true covariance, $Q_{\mathrm{F}}$ would follow a chi-squared distribution on $g$ degrees of freedom. It is estimated from $m$ patients, so, as $t$ is referred to Satterthwaite's degrees of freedom rather than the normal, $Q_{\mathrm{F}}$ is referred to the approximate Hotelling's $T^2$ test of Tipton [66] and Pustejovsky & Tipton [61]. It treats $\mathbf{V}^{\mathrm{CR2}}$ as a Wishart matrix on $\nu_{\mathrm{F}}$ degrees of freedom and, as for Hotelling's $T^2$, refers

$$F := \frac{\nu_{\mathrm{F}} - g + 1}{\nu_{\mathrm{F}}\, g}\, Q_{\mathrm{F}}$$

to the $F$ distribution on $g$ and $\nu_{\mathrm{F}} - g + 1$ degrees of freedom. $\nu_{\mathrm{F}}$ is chosen so that $\mathbf{V}^{\mathrm{CR2}}$ varies as much as such a Wishart matrix would.

To match $\mathbf{V}^{\mathrm{CR2}}$ with a Wishart matrix, first rescale it so that its expected value is the identity. Let $\mathbf{W} = (\mathbf{LBL}^{\top})^{-1/2}$ and $\mathbf{s}_i^* = \mathbf{W}\mathbf{s}_i$, so that $\mathbf{W}\mathbf{V}^{\mathrm{CR2}}\mathbf{W} = \sum_i \mathbf{s}_i^* \mathbf{s}_i^{*\top}$; the proposition below shows its mean is $\mathbf{I}$. Patient $i$'s rescaled term is $\mathbf{s}_i^* = \mathbf{P}_i \tilde{\mathbf{r}}_i$, with $\mathbf{P}_i = \mathbf{WLBX}_i^{\top}\mathbf{A}_i$. Let

$$\boldsymbol{\Sigma}_{ik} := \mathbf{P}_i \big( 1\{i = k\}\, \mathbf{I} - \mathbf{X}_i \mathbf{B} \mathbf{X}_k^{\top} \big) \mathbf{P}_k^{\top};$$

the proof below shows that, under the working model, this is the $g \times g$ covariance of the terms of patients $i$ and $k$, $\mathrm{Cov}(\mathbf{s}_i^*, \mathbf{s}_k^*)$. Each of the $g^2$ entries of $\mathbf{W}\mathbf{V}^{\mathrm{CR2}}\mathbf{W}$ has its own variance, and one $\nu_{\mathrm{F}}$ cannot in general match them all. Following Tipton [66], $\nu_{\mathrm{F}}$ matches their sum, called the summed variance.

**Proposition 15** (Degrees of freedom of the overall test)**.** *Under the working model with normal errors,* $\mathbf{W}\mathbf{V}^{\mathrm{CR2}}\mathbf{W}$ *has mean* $\mathbf{I}$ *and summed variance* $\sum_{i,k} \big\{ (\operatorname{tr} \boldsymbol{\Sigma}_{ik})^2 + \operatorname{tr}(\boldsymbol{\Sigma}_{ik}\boldsymbol{\Sigma}_{ik}) \big\}$*. A Wishart matrix with mean* $\mathbf{I}$ *on* $\nu_{\mathrm{F}}$ *degrees of freedom has summed variance* $g(g+1)/\nu_{\mathrm{F}}$*, so matching the two gives*

$$\nu_{\mathrm{F}} = \frac{g(g+1)}{\sum_{i,k} \big\{ (\operatorname{tr} \boldsymbol{\Sigma}_{ik})^2 + \operatorname{tr}(\boldsymbol{\Sigma}_{ik}\boldsymbol{\Sigma}_{ik}) \big\}}.$$

*With* $g = 1$*,* $\nu_{\mathrm{F}}$ *is the* $\nu$ *of Proposition 14 and* $F = t^2$*.*

*Proof.* The proof finds the mean and the summed variance of $\mathbf{W}\mathbf{V}^{\mathrm{CR2}}\mathbf{W} = \sum_i \mathbf{s}_i^* \mathbf{s}_i^{*\top}$, finds the summed variance of a Wishart matrix, and sets the two equal. The mean and the variance both depend on how the patients' terms $\mathbf{s}_i^*$ vary together, so the first step is their covariance. As in the proof of Proposition 14, under the working model the stacked whitened residuals are $\tilde{\mathbf{r}} = (\mathbf{I} - \mathbf{H})\mathbf{e}$ with $\mathbf{e}$ standard normal. Since $(\mathbf{I} - \mathbf{H})^2 = \mathbf{I} - \mathbf{H}$, $\mathrm{Cov}(\tilde{\mathbf{r}}_i, \tilde{\mathbf{r}}_k)$ is the $(i,k)$ block of $\mathbf{I} - \mathbf{H}$, that is $1\{i = k\}\, \mathbf{I} - \mathbf{X}_i \mathbf{B} \mathbf{X}_k^\top$. So $\mathrm{Cov}(\mathbf{s}_i^*, \mathbf{s}_k^*) = \mathbf{P}_i \,\mathrm{Cov}(\tilde{\mathbf{r}}_i, \tilde{\mathbf{r}}_k) \mathbf{P}_k^\top = \boldsymbol{\Sigma}_{ik}$, and the $\mathbf{s}_i^*$ are normal with mean $\mathbf{0}$.

Since $\mathbf{s}_i^*$ has mean $\mathbf{0}$, the mean of $\mathbf{s}_i^* \mathbf{s}_i^{*\top}$ is its covariance with itself, $\boldsymbol{\Sigma}_{ii}$, so the mean of $\sum_i \mathbf{s}_i^* \mathbf{s}_i^{*\top}$ is $\sum_i \boldsymbol{\Sigma}_{ii}$. Since $\mathbf{A}_i = (\mathbf{I} - \mathbf{X}_i \mathbf{B} \mathbf{X}_i^\top)^{-1/2}$ is symmetric, $\mathbf{A}_i(\mathbf{I} - \mathbf{X}_i \mathbf{B} \mathbf{X}_i^\top)\mathbf{A}_i = \mathbf{I}$, so $\boldsymbol{\Sigma}_{ii} = \mathbf{W}\mathbf{L}\mathbf{B}\mathbf{X}_i^\top \mathbf{X}_i \mathbf{B}\mathbf{L}^\top \mathbf{W}$. Summing $\boldsymbol{\Sigma}_{ii}$ over $i$, and using $\sum_i \mathbf{X}_i^\top \mathbf{X}_i = \mathbf{B}^{-1}$,

$$\sum_i \boldsymbol{\Sigma}_{ii} = \mathbf{W}\mathbf{L}\mathbf{B}\Big(\sum_i \mathbf{X}_i^\top \mathbf{X}_i\Big)\mathbf{B}\mathbf{L}^\top \mathbf{W} = \mathbf{W}\mathbf{L}\mathbf{B}\mathbf{L}^\top \mathbf{W} = \mathbf{I},$$

by the choice of $\mathbf{W}$.

The summed variance adds up the variances of the $g^2$ entries, so take one entry at a time. Entry $(a,b)$ of $\mathbf{W}\mathbf{V}^{\mathrm{CR2}}\mathbf{W}$ is $\sum_i s_{ia}^* s_{ib}^*$, so its variance is $\sum_{i,k} \mathrm{Cov}(s_{ia}^* s_{ib}^*,\, s_{ka}^* s_{kb}^*)$. For normal variables $x_1, \ldots, x_4$ with mean 0, Isserlis' theorem gives $\mathrm{Cov}(x_1 x_2,\, x_3 x_4) = \mathrm{Cov}(x_1, x_3)\,\mathrm{Cov}(x_2, x_4) + \mathrm{Cov}(x_1, x_4)\,\mathrm{Cov}(x_2, x_3)$. Writing $\Sigma_{ik,ab}$ for entry $(a,b)$ of $\boldsymbol{\Sigma}_{ik}$, which is $\mathrm{Cov}(s_{ia}^*, s_{kb}^*)$,

$$\mathrm{Cov}(s_{ia}^* s_{ib}^*,\, s_{ka}^* s_{kb}^*) = \Sigma_{ik,aa}\Sigma_{ik,bb} + \Sigma_{ik,ab}\Sigma_{ik,ba}.$$

Summed over $a$ and $b$, the first term gives $\sum_a \Sigma_{ik,aa} \sum_b \Sigma_{ik,bb} = (\mathrm{tr}\, \boldsymbol{\Sigma}_{ik})^2$ and the second gives $\sum_a (\boldsymbol{\Sigma}_{ik}\boldsymbol{\Sigma}_{ik})_{aa} = \mathrm{tr}(\boldsymbol{\Sigma}_{ik}\boldsymbol{\Sigma}_{ik})$. Summing over $i$ and $k$ gives the stated summed variance.

This is the summed variance to be matched. For the benchmark, the same quantity is needed for a Wishart matrix. A Wishart matrix with mean $\mathbf{I}$ on $\nu_{\mathrm{F}}$ degrees of freedom is $\nu_{\mathrm{F}}^{-1} \sum_{j=1}^{\nu_{\mathrm{F}}} \boldsymbol{\xi}_j \boldsymbol{\xi}_j^\top$, with $\boldsymbol{\xi}_j$ independent standard normal vectors of length $g$. Entry $(a,b)$ is the average of $\nu_{\mathrm{F}}$ independent copies of $\xi_a \xi_b$, whose variance is 2 when $a = b$ and 1 otherwise, so the entry's variance is $(1 + 1\{a = b\})/\nu_{\mathrm{F}}$. The $g$ diagonal entries contribute $2g/\nu_{\mathrm{F}}$ and the other $g^2 - g$ contribute $(g^2 - g)/\nu_{\mathrm{F}}$, a total of $g(g+1)/\nu_{\mathrm{F}}$. Setting $g(g+1)/\nu_{\mathrm{F}}$ equal to the summed variance of $\mathbf{W}\mathbf{V}^{\mathrm{CR2}}\mathbf{W}$ and solving for $\nu_{\mathrm{F}}$ gives the stated formula.

It remains to check that, with two groups, this is the test of Proposition 14. With $g = 1$, $\mathbf{L} = \mathbf{c}^\top$, $\mathbf{W} = (\mathbf{c}^\top \mathbf{B} \mathbf{c})^{-1/2}$ is a number and $\mathbf{P}_i = \mathbf{W}\mathbf{a}_i^\top$, with $\mathbf{a}_i$ of Section 6.1.4. So $\boldsymbol{\Sigma}_{ik}$ is the number $\Omega_{ik}/(\mathbf{c}^\top \mathbf{B}\mathbf{c})$, with $\Omega_{ik}$ of Proposition 14, and the mean above gives $\mathbf{c}^\top \mathbf{B}\mathbf{c} = \sum_i \Omega_{ii} = \mathrm{tr}\, \boldsymbol{\Omega}$. Both terms of the summed variance are then $\Omega_{ik}^2/(\mathrm{tr}\, \boldsymbol{\Omega})^2$, so it equals $2\,\mathrm{tr}(\boldsymbol{\Omega}^2)/(\mathrm{tr}\, \boldsymbol{\Omega})^2$ and $\nu_{\mathrm{F}} = (\mathrm{tr}\, \boldsymbol{\Omega})^2/\mathrm{tr}(\boldsymbol{\Omega}^2) = \nu$. Finally $F = Q_{\mathrm{F}} = t^2$ on 1 and $\nu$ degrees of freedom, which is the test of $t$ on $\nu$. □

In the example, if $\nu_{\mathrm{F}} = 20$, then $F = (19/40) \times 15.4 = 7.3$ on 2 and 19 degrees of freedom, and $p = 0.0045$. Treated as chi-squared, $Q_{\mathrm{F}} = 15.4$ would give $p = 0.00046$, ten times smaller: with few patients the correction matters as much as it does for a single difference.

The pair is tested only when every group passes the rules of Section 5.8, and not when $\nu_{\mathrm{F}} - g + 1 < 1$.

##### 6.2.3 Comparing two groups

For a pair called by the overall test, the next question is which groups differ. Any two groups $a$ and $b$ are compared by the two-group test of Section 5 on the patients of those two groups only, with $\omega^2$ and $\tau^2$ estimated again from them, so the comparison is the analysis those patients would get on their own. The scale stays the one chosen from all images, as for the overall test. A comparison is tested when its two groups pass the rules of Section 5.8, so a sparse third group does not block it.

Choosing which pairs to follow up uses the data, so the comparisons within the called pairs need a stricter threshold than $\alpha$. We use the procedure of Benjamini & Bogomolov [67] for testing within selected families of hypotheses, with each pair as a family and its comparisons as the hypotheses. Say $m_{\mathrm{p}}$ pairs have an overall test and $R$ of them are called at false discovery rate $\alpha$. Within each called pair, the comparisons are tested at level $\alpha R/m_{\mathrm{p}}$ with a Bonferroni correction. So, with $K$ comparisons tested in the pair, a comparison is significant when

$$p \leq \frac{\alpha R}{m_{\mathrm{p}} K}.$$

For example, with 300 pairs, 30 called at $\alpha = 0.05$ and three groups, the threshold is $0.05 \times 30/(300 \times 3) = 0.0017$.

Benjamini & Bogomolov [67] show that this procedure keeps the expected share of called pairs with at least one false comparison at or below $\alpha$, under two conditions. First, the pairs are selected using the $p$-values of their own comparisons. Second, the $p$-values of different pairs are independent or positively dependent. Neither holds exactly here. The pairs are selected by the overall test, which is a separate test, and different pairs use the same patients. So the guarantee is approximate.

#### 6.3 Time-to-event outcomes

For a time-to-event outcome, such as overall survival, patient $i$ has a follow-up time $T_i$ and an event indicator $d_i$: 1 if the event (for example, death) was seen and 0 if follow-up ended first, for example because the patient was still alive when the study closed, so that the patient is censored. The question for a pair is whether its co-localisation goes with how soon the event comes. The time cannot simply take the place of the group in the model of Section 4.1: a censored time is only a lower bound, so a patient censored at one year and one who died at one year are not alike. Hence, the Cox model is the standard model in this situation, and it allows for censoring. From a Cox model, each patient gets one number, the martingale residual, which says whether they had the event sooner or later than expected. This number takes the place of the group in the model, so the question becomes whether co-localisation rises or falls with it.

##### 6.3.1 The martingale residual

A Cox model is fitted once, with the patients' covariates $\mathbf{w}_i$ but not the effect (extra fraction or extra count) of any pair, and the same model is used for every pair:

$$h_i(t) = h_0(t) \exp\big(\mathbf{w}_i^\top \boldsymbol{\zeta}\big).$$

Here $h_i(t)$ is patient $i$'s hazard, their rate of the event at time $t$ given no event before $t$. $\boldsymbol{\zeta}$ is the vector of the covariates' coefficients. $h_0(t)$ is the baseline hazard, the hazard of a patient whose covariates are all 0. The fit gives the estimate $\hat{\boldsymbol{\zeta}}$, i.e. the estimated log hazard ratios. Then $\hat{\boldsymbol{\zeta}}$ gives the number of events $\hat{H}_i$ that patient $i$ is expected to have up to their last follow-up,

$$\hat{H}_i := e^{\mathbf{w}_i^\top \hat{\boldsymbol{\zeta}}} \sum_{k:\, t_k \le T_i} \frac{1}{\sum_{j \in \mathcal{R}(t_k)} e^{\mathbf{w}_j^\top \hat{\boldsymbol{\zeta}}}},$$

where $t_k$ are the event times and $\mathcal{R}(t_k)$ are the patients still followed at $t_k$ [68]. Tied event times use Efron's version [69]. The martingale residual [70] is

$$M_i := d_i - \hat{H}_i.$$

A patient who died at two years when $\hat{H}_i = 0.4$ has $M_i = 0.6$: they had the event sooner than expected. A patient censored at five years with $\hat{H}_i = 0.7$ has $M_i = -0.7$: they went longer without it than expected. The residuals sum to 0 over the patients, and $\sum_i w_{ia} M_i = 0$ for each covariate $a$ (Proposition 16), so the $M_i$ carry no trace of the covariates. An image-level covariate enters the Cox model as its average over the patient's images. The abundance share of Section 6.1.5 does not enter the Cox model: it is the share of the pair's $A$ cells, so it differs between pairs, while the Cox model is fitted once for every pair. It enters only the model of the effect described below.

##### 6.3.2 The model and the test

With groups, the model of Section 6.1 has one mean $\delta_c$ for each group. Here there are no groups, so the group indicators are replaced by an intercept and $M_i$:

$$Y_{ij} = \theta_0 + \theta_1 M_i + \mathbf{x}_{ij}^\top \boldsymbol{\beta} + \left(\bar{\mathbf{x}}_i^{\mathrm{I}}\right)^\top \boldsymbol{\lambda} + b_i + \eta_{ij} + \varepsilon_{ij}.$$

The intercept $\theta_0$ is the mean effect of a patient with $M_i = 0$, who had the event when expected, and the slope $\theta_1$ is how much the mean effect changes per unit of $M_i$. The other terms are those of Section 6.1.1, with the patient averages only for image-level covariates (Section 6.1.2). A positive $\theta_1$ means that patients who had the event sooner than expected have higher co-localisation, and a negative $\theta_1$ that they have lower co-localisation. Therefore, the test of the pair is of the null hypothesis

$$H_0 : \ \theta_1 = 0,$$

that co-localisation does not change with $M_i$, and so does not go with how soon the event comes.

The model is fitted as in Section 6.1: $\omega^2$ within patients, and $\tau^2$ by Paule–Mandel on the patient summaries. $\theta_1 = 0$ is tested with the CR2 variance of the whole fit on Satterthwaite's degrees of freedom (Section 6.1.4), with the contrast $\mathbf{c}$ picking out $\theta_1$.

**Why the covariates enter both models.** The covariates enter both the Cox model and the model of the effect, and each does a different job. In the Cox model, they make $M_i$ the part of the outcome that the covariates do not explain. In the model of the effect, they remove the part of $Y_{ij}$ that the covariates explain. Putting them in the Cox model alone would be valid but would lose power. A covariate could then not cause a false positive. Since $\sum_i w_{ia} M_i = 0$, $M_i$ is uncorrelated with every covariate, so a covariate that shifts the effect cannot make it appear to change with $M_i$. However, if the covariates were not added to the model of the effect, the covariate's effect on $Y_{ij}$ would not be removed, so it would stay in the residuals. Larger residuals give a larger CR2 standard error of $\hat{\theta}_1$, but $\hat{\theta}_1$ itself is hardly more variable, since $M_i$ is uncorrelated with the covariate. The standard error is then too large, so the test is too cautious and has less power.

**The link with the Cox model.** The obvious way to ask whether the pair's effect goes with survival is to add each patient's mean effect $\bar{Y}_i$ to the Cox model as a covariate and test whether its coefficient is 0. We do not do this, for two reasons. First, the Cox model treats each $\bar{Y}_i$ as exact, so a patient with a handful of cells, whose mean effect is very noisy, counts as much as a patient with thousands. Second, its standard error comes from the Cox model and is right only if that model is. The test of $\theta_1$ is still close to this obvious test. The proposition below shows that, in the simplest case, $\hat{\theta}_1$ has the same numerator as Cox's score statistic for $\bar{Y}_i$, the usual statistic for testing that coefficient. The test of $\theta_1$ is therefore the obvious test with both problems fixed: patients are weighted by how precisely their effect is measured, and the standard error is the CR2 one.

**Proposition 16** (The slope and Cox's score)**.** *Suppose there are no tied event times.*

*(i) For any number $x_i$ per patient, Cox's score statistic for adding $x_i$ to the Cox model of Section 6.3, at $\hat{\boldsymbol{\zeta}}$ and a coefficient of* 0, *is $\sum_i x_i M_i$. In particular, $\sum_i M_i = 0$ and $\sum_i w_{ia} M_i = 0$ for each covariate $a$.*

*(ii) Without covariates in the model of the effect and with equal patient weights $u_i = u$, the numerator of $\hat{\theta}_1$ is $u \sum_i M_i \bar{Y}_i$, which is $u$ times Cox's score statistic for adding each patient's mean effect $\bar{Y}_i$.*

*Proof.* Part (i) links Cox's score to the martingale residuals, and part (ii) links $\hat{\theta}_1$ to the score. For part (i), first write out the score. Let $s_k := \exp(\mathbf{w}_k^\top \hat{\boldsymbol{\zeta}})$ be patient $k$'s risk score, and $S(t) = \sum_{k \in \mathcal{R}(t)} s_k$ the total risk score of the patients still followed at $t$. Differentiating the log partial likelihood [71] with respect to the new coefficient, at 0, gives Cox's score

$$U = \sum_i d_i \big\{ x_i - \bar{x}(T_i) \big\}, \qquad \bar{x}(t) := \frac{\sum_{k \in \mathcal{R}(t)} x_k s_k}{S(t)},$$

where $\bar{x}(t)$ is the average of $x$ over the patients still followed at $t$, weighted by their risk scores. Each event adds the gap between the $x_i$ of the patient who had it and this average.

Part (i) says that this score equals $\sum_k x_k M_k$. Since $M_k = d_k - \hat{H}_k$, $\sum_k x_k M_k = \sum_k x_k d_k - \sum_k x_k \hat{H}_k$, and the first sum is the first part of $U$. It remains to show that $\sum_k x_k \hat{H}_k = \sum_i d_i \, \bar{x}(T_i)$. With no ties, the event times are the $T_i$ with $d_i = 1$, so the formula for $\hat{H}_k$ above reads $\hat{H}_k = s_k \sum_{i:\, d_i = 1,\, T_i \le T_k} 1/S(T_i)$. Swapping the order of the sums, and noting that the patients with $T_k \ge T_i$ are those still followed at $T_i$,

$$\sum_k x_k \hat{H}_k = \sum_{i:\, d_i = 1} \frac{1}{S(T_i)} \sum_{k \in \mathcal{R}(T_i)} x_k s_k = \sum_i d_i \, \bar{x}(T_i).$$

So $\sum_k x_k M_k = U$.

The two sums in part (i) follow by choosing $x$. With $x_k = 1$, every $\bar{x}(t)$ is 1, so $U = 0$ and $\sum_k M_k = 0$. With $x_k = w_{ka}$, $U$ is the score of covariate $a$ at $\hat{\boldsymbol{\zeta}}$, which is 0 since $\hat{\boldsymbol{\zeta}}$ maximises the partial likelihood; so $\sum_k w_{ka} M_k = 0$.

For part (ii), write $\hat{\theta}_1$ as a sum over patients and compare it with the score. Without covariates, both columns of the model, 1 and $M_i$, are the same in every image of a patient, so the fit is a weighted regression of the patient means $\bar{Y}_i$ on $M_i$ with weights $u_i$ (Section 4.2). Its slope is

$$\hat{\theta}_1 = \frac{\sum_i u_i (M_i - \bar{M}_u) \bar{Y}_i}{\sum_i u_i (M_i - \bar{M}_u)^2},$$

with $\bar{M}_u$ the $u$-weighted mean of the $M_i$. With equal weights, $\bar{M}_u$ is the plain mean of the $M_i$, which is 0 by part (i), so the numerator is $u \sum_i M_i \bar{Y}_i$. By part (i) with $x_i = \bar{Y}_i$, this is $u$ times Cox's score for adding $\bar{Y}_i$. □

##### 6.3.3 When a pair is tested

Patients with no follow-up time or event status, or with a covariate missing in every image, are removed before anything else. This includes the choice between the gain and loss scales (Section 3.3), so the result is exactly that of the data without them.

A time-to-event outcome is tested at one radius only, since the combination of several radii (Section 6.4) is for two groups.

A pair's informative patients are those with an outcome and an effect for the pair. The pair is tested only when these patients carry enough information:

1. There is at least one event and at least one censored patient.

2. There are at least two more patients than the model has columns: 2 for $\theta_0$ and $\theta_1$, one for each covariate, and one more for each image-level covariate's patient average. That is 4 patients without covariates, 5 with one patient-level covariate and 6 with the abundance share. The two spare patients leave information for the variance once the model is fitted. With fewer, the model is nearly saturated: it has almost as many columns as patients, so it fits them almost exactly and the CR2 variance has little to go on. In calibration with covariates and 3 to 5 informative patients, 8.1% of null tests were rejected at $p < 0.05$.

As for every test, the pair is not tested when $\nu < 1$. The number of informative events is reported with each pair, since with fewer than 3 the test has little power.

##### 6.3.4 The hazard ratio

The test gives a $p$-value, but $\theta_1$ is hard to interpret: it is the change in the mean effect per unit of $M_i$. The usual effect size for a time-to-event outcome is a hazard ratio, so one is also reported, from a Cox model with the pair's effect as a covariate. As explained above, this model is not used for the test; here it only gives an effect size, and the first two of the three steps below reduce its problems.

**1. The model of the effect without $M_i$.** The model of the effect above is fitted again with $\theta_1 M_i$ left out:

$$Y_{ij} = \theta_0 + \mathbf{x}_{ij}^\top \boldsymbol{\beta} + \left(\bar{\mathbf{x}}_i^{\mathrm{I}}\right)^\top \boldsymbol{\lambda} + b_i + \eta_{ij} + \varepsilon_{ij}.$$

The patients' effects from this fit become the predictor in the Cox model. If $M_i$ were in the fit, the outcome would be part of its own predictor. Each patient's mean effect, taken at the average covariates, is

$$y_i := \frac{1}{\Gamma_i} \sum_{j=1}^{J_i} \gamma_{ij} \Big( Y_{ij} - \mathbf{x}_{ij}^\top \hat{\boldsymbol{\beta}} - \left(\bar{\mathbf{x}}_i^{\mathrm{I}}\right)^\top \hat{\boldsymbol{\lambda}} \Big),$$

with the image weights $\gamma_{ij} = 1/(s_{ij}^2 + \omega^2)$ and their sum $\Gamma_i$ (Section 4.2).

**2. Shrinking noisy patients.** A patient with few cells has a noisy $y_i$, and noise in a predictor pulls its hazard ratio towards 1 [72]. Under the model just fitted, $y_i$ is the patient's own level plus noise:

$$y_i = \theta_0 + b_i + e_i, \qquad \mathrm{Var}(b_i) = \tau^2, \qquad \mathrm{Var}(e_i) = 1/\Gamma_i,$$

where $b_i$ is the patient's departure (Section 4.1) and $e_i$ is the noise left after averaging the patient's images. The Cox model should be given the level $\theta_0 + b_i$, not $y_i$. The level is not observed, so it is replaced by its best linear prediction from $y_i$:

$$\tilde{Y}_i := \theta_0 + \frac{\mathrm{Cov}(b_i, y_i)}{\mathrm{Var}(y_i)} (y_i - \theta_0) = \theta_0 + \frac{\tau^2}{\tau^2 + 1/\Gamma_i} (y_i - \theta_0),$$

since $\mathrm{Cov}(b_i, y_i) = \mathrm{Var}(b_i) = \tau^2$ and $\mathrm{Var}(y_i) = \tau^2 + 1/\Gamma_i$. The estimates $\hat{\theta}_0$ and $\hat{\tau}^2$ are put in place of $\theta_0$ and $\tau^2$. If $b_i$ and $e_i$ are normal, $\tilde{Y}_i$ is the expected level given $y_i$. Replacing a noisy predictor by this expectation is known as regression calibration, and it removes most of the pull towards 1 [72]. The factor $\hat{\tau}^2/(\hat{\tau}^2 + 1/\Gamma_i)$ is the share of the spread of $y_i$ that is real variation between patients. It is near 1 for a patient with many cells, whose $y_i$ is kept, and near 0 for a patient with few, whose $y_i$ is pulled to the mean.

**3. The Cox model.** The shrunken effects are standardised across patients,

$$S_i := \frac{\tilde{Y}_i - \mathrm{mean}(\tilde{Y})}{\mathrm{SD}(\tilde{Y})},$$

and entered into a Cox model with the covariates $\mathbf{w}_i$:

$$h_i(t) = h_0(t) \exp\big(\phi S_i + \mathbf{w}_i^\top \boldsymbol{\zeta}\big).$$

Here $\boldsymbol{\zeta}$ is fitted again, together with $\phi$. The reported hazard ratio is $e^{\hat{\phi}}$: a patient whose shrunken effect is one standard deviation higher, with the same covariates, has $e^{\hat{\phi}}$ times the hazard. For example, a hazard ratio of 1.5 means 1.5 times the hazard.

The hazard ratio is per standard deviation, not per unit of the effect, since a unit has no useful size here. One unit of the extra fraction is 100 percentage points, far more than any two patients differ by. A standard deviation is instead the size of a typical difference between patients for that pair. Hazard ratios per standard deviation are therefore on the same scale for every pair, and for both the extra fraction and the extra count.

**One test per pair.** The hazard ratio has no $p$-value of its own, and this is deliberate. The $p$-value of a pair is that of the test of $\theta_1$ alone, so each pair has one test and one $p$-value to adjust for multiple testing. The hazard ratio only states the size of the association in familiar units. Its standard error is reported, but it treats $\tilde{Y}_i$ as known, although $\hat{\theta}_0$, $\hat{\tau}^2$ and the fitted covariate terms behind it are estimated. So the standard error is approximate and is not turned into a $p$-value.

When $\hat{\tau}^2$ is so small that every patient is shrunk almost to the mean (every factor $\hat{\tau}^2/(\hat{\tau}^2 + 1/\Gamma_i)$ below 0.01), the shrunken effects barely differ and their standard deviation is close to 0, so no hazard ratio is reported.

#### 6.4 Several radii

The radius sets how close two cells must be to count as neighbours, and the right distance is often not known in advance. Testing the pair at several radii and keeping the smallest $p$-value would give too many false positives, since each radius is another chance of a small $p$-value by luck. With four unrelated radii, at least one would reach $p < 0.05$ in $1 - 0.95^4 = 18.5\%$ of pairs with no difference. Multiplying the smallest $p$-value by the number of radii $K$ (Bonferroni) prevents this, but is too strict here. Nearby radii count many of the same neighbours, so their statistics move together, and $K$ such radii are fewer than $K$ separate chances.

This section gives a test over several radii that allows for how much they overlap. Section 6.4.1 puts the radii's statistics on one scale and estimates how much they move together. Three $p$-values can then be computed from them:

1. the joint $p$-value (Section 6.4.2), the probability itself under a normal approximation, which takes milliseconds to seconds per pair;
2. Davies' bound (Section 6.4.3), the default, a closed form slightly above the joint $p$-value, which takes microseconds per pair;
3. the Cauchy combination (Section 6.4.4), which needs no correlations but can dilute a strong difference at one radius.

Whichever is used, the combined $p$-values of the pairs that are tested are adjusted together by the method of Benjamini & Hochberg [64], as for a single radius. Several radii are combined only for a comparison of two groups.

When the radius of interest is known, testing that radius alone is more powerful, since it pays no penalty for other radii. When it is not, about four radii spanning the range of interest are recommended, none below about one cell diameter. More radii, closely spaced, add little information, since adjacent radii are then highly correlated, and their correlations are hard to estimate with few patients.

##### 6.4.1 Putting the radii on one scale

Let the pair be tested at radii $r_1 < \cdots < r_K$, with statistic $t_k$ on $\nu_k$ degrees of freedom at radius $r_k$ (Section 5.7). Radii at which the pair is not tested (Section 5.8) are left out, and if only one radius is left, its own $p$-value is reported.

The statistics cannot be compared as they are, since the degrees of freedom differ between radii, and the same $t$ is stronger evidence on more degrees of freedom: $t = 2.5$ gives $p = 0.037$ on 8 degrees of freedom but $p = 0.017$ on 40. So each statistic is first converted to the normal value with the same two-sided $p$-value,

$$z_k := \Phi^{-1}\{1 - \Pr(T_{\nu_k} > |t_k|)\},$$

where $\Phi$ is the standard normal distribution function, so that $\Phi(x)$ is the chance that a standard normal variable is below $x$, and $\Phi^{-1}$ is its inverse. Then $2\Phi(-z_k)$ is radius $k$'s two-sided $p$-value, and the two examples give $z = 2.09$ and $z = 2.40$. The value $z_k$ is never negative: it measures the size of the difference at radius $k$, whatever its sign.

The cell type pair's co-localisation is then summarised by the largest of these, $z_{\max} = \max_k z_k$, and the radius attaining it is reported as the radius with the strongest evidence. The $p$-value of the pair is the probability that the largest would be at least this large if the groups did not differ at any radius:

$$p := \Pr\Big(\max_k |Z_k| \geq z_{\max}\Big), \qquad (Z_1, \ldots, Z_K) \sim N(\mathbf{0}, \mathbf{R}).$$

Here $Z_1, \ldots, Z_K$ stand for the radii's normal values when the groups do not differ. Each is standard normal, and $\mathbf{R}$ is their correlation matrix, which says how much the radii overlap. Two extremes show its role. If all radii were identical, $p$ would be one radius's $p$-value, $2\Phi(-z_{\max})$. If they were unrelated, $p = 1 - \{1 - 2\Phi(-z_{\max})\}^K$, close to Bonferroni's $K \cdot 2\Phi(-z_{\max})$. Real radii lie in between. This $p$ is a normal approximation, not an exact probability, since the $z_k$ are only approximately standard normal and jointly normal, and $\mathbf{R}$ is estimated from the data.

**The correlation between radii.** The same patients are compared at every radius, so a patient whose effect is high at one radius tends to be high at the next. The correlation is estimated from each patient's share of the standard error. At radius $r_k$, the CR2 variance of the difference $D$ is a sum of one squared term per patient, $\sum_i \psi_{ik}^2 = V_1 + V_2$, where patient $i$'s signed term is

$$\psi_{ik} := s_i \, \frac{u_{ik}}{U_{c(i)k}} \, \frac{\hat{e}_{ik}}{\sqrt{1 - h_{ik}}}.$$

Here $s_i = -1$ for a patient of group 1 and $+1$ for group 2, since $D = \hat{\delta}_2 - \hat{\delta}_1$, and $u_{ik}$, $U_{ck}$, $\hat{e}_{ik}$ and $h_{ik}$ are those of Proposition 10 at radius $r_k$. With covariates, $\psi_{ik}$ is patient $i$'s term of the CR2 variance of the adjusted difference (Section 6.1). The variance at one radius sums each patient's squared term, and in the same way the covariance between two radii sums the products of each patient's two terms. Dividing by the two standard errors gives the correlation of the statistics at radii $k$ and $l$,

$$\hat{\rho}_{kl} := \frac{\sum_i \psi_{ik}\psi_{il}}{\sqrt{\sum_i \psi_{ik}^2 \sum_i \psi_{il}^2}},$$

the cluster-robust estimate with patients as the clusters. Like $V_c$, it makes no assumption about how the neighbours counted at the two radii overlap.

##### 6.4.2 The joint $p$-value

With $\mathbf{R} = (\hat{\rho}_{kl})$, $p$ is the probability that a $K$-dimensional normal vector falls outside the box $[-z_{\max}, z_{\max}]^K$. It has no closed form and is computed numerically. To keep it accurate when it is small, it is split by the first radius, in order of radius, whose $|Z_k|$ reaches $z_{\max}$:

$$p = 2\Phi(-z_{\max}) + \sum_{k=2}^{K} 2\Pr\big(Z_k \geq z_{\max},\ |Z_j| < z_{\max} \text{ for } j < k\big).$$

The first term is the chance that radius 1 already reaches $z_{\max}$. The $k$-th term is the chance that radius $k$ is the first to reach it. It is twice the chance for $Z_k \geq z_{\max}$, since $\mathbf{Z}$ and $-\mathbf{Z}$ have the same distribution.

Each term is computed by the quasi-Monte Carlo method of Genz [73] [see also 74] to a relative error of about $10^{-3}$. Since each term is held to 0.1% of itself, the error of $p$ is at most about 0.1% of $p$, however small $p$ is. Computing $p$ as one minus the chance of falling inside the box would not keep this accuracy: that chance is close to 1, and an error of $10^{-3}$ in it would swamp a $p$ of $10^{-4}$. With two radii, $p$ is computed directly by numerical integration, which is more precise still. This is the accuracy of the computation only: $p$ itself is still the normal approximation above.

A radius whose correlation with a smaller radius already kept exceeds 0.995 is left out. Such a radius adds almost no information, and it makes $\mathbf{R}$ nearly singular, so the computation becomes unstable.

##### 6.4.3 Davies' bound (the default)

The joint $p$-value takes milliseconds per pair with 4 radii and seconds with 25. Davies' bound replaces it by a closed form, an upper bound on $p$ that takes microseconds per pair for any number of radii. It is the default for this reason.

**Proposition 17** (Davies' bound). *With* $\Lambda := \sum_{k=2}^{K} \arccos \hat{\rho}_{k-1,k}$*,*

$$p \leq p_{\mathrm{D}} := \min\Big\{1,\ 2\Phi(-z_{\max}) + \frac{\Lambda}{\pi}\, e^{-z_{\max}^2/2}\Big\}.$$

*Proof.* The idea is to join the radii by a continuous path, and to bound the chance that the path ever reaches $z_{\max}$. This chance is at least $p$, and unlike $p$ it can be bounded in closed form.

First, the radii are placed on a sphere. Two normal variables with variance 1 and correlation $\rho$ can be pictured as two unit vectors at angle $\arccos \rho$: the more correlated they are, the closer they are. So $Z_1, \ldots, Z_K$ are points on a unit sphere, with $Z_{k-1}$ and $Z_k$ at angle $\arccos \hat{\rho}_{k-1,k}$.

Next, these points are joined in order of radius by arcs of great circles, the shortest paths on the sphere. Each point $\theta$ on the arcs is again a normal variable $Z(\theta)$ with variance 1. This gives a normal process that passes through every $Z_k$ and has total length $\Lambda$. Since it passes through every $Z_k$, its largest $|Z(\theta)|$ is at least $\max_k |Z_k|$, so

$$p \leq \Pr\Big(\sup_{\theta} |Z(\theta)| \geq z_{\max}\Big).$$

It remains to bound the right-hand side. The process can reach $z_{\max}$ in only two ways. Either $|Z(\theta)|$ is already at least $z_{\max}$ at the start, which has probability $2\Phi(-z_{\max})$, or it crosses $z_{\max}$ upwards somewhere along the path. The chance of at least one upcrossing is at most the expected number of upcrossings. By Rice's formula, a normal process with variance 1 moving at unit speed, as along a great circle, crosses a level $c$ upwards on average $e^{-c^2/2}/(2\pi)$ times per unit length. Both $Z$ and $-Z$ count, so the expected number of upcrossings of $z_{\max}$ by $|Z(\theta)|$ over the length $\Lambda$ is $\Lambda e^{-z_{\max}^2/2}/\pi$ [75]. Adding the two ways gives the bound, and a probability is at most 1. □

The bound follows Davies [75, 76], who studied tests in which a parameter is present only under the alternative: here the radius, which plays no part if the groups do not differ at any radius. Davies bounds the largest value of a normal process by counting its upcrossings. The proposition doubles his bound, since large differences of either sign count, and applies it to the path through the radii built in the proof.

$\Lambda$ measures how different the radii are. Each pair of adjacent radii adds the angle $\arccos \hat{\rho}$, which is 0 for identical radii and $\pi/2$ for unrelated ones, so radii that nearly repeat each other add little to $p_{\mathrm{D}}$. For example, with $K = 4$ radii and $z_{\max} = 2.5$, one radius alone gives $2\Phi(-2.5) = 0.012$. With adjacent correlations of 0.99, 0.9 and 0, $p_{\mathrm{D}} = 0.018$, 0.031 and 0.078, while Bonferroni gives 0.050 in each case. This is why several close radii keep much more power than a Bonferroni correction, though less than a single well-chosen radius, which pays no penalty.

The bound needs only the $K-1$ correlations between adjacent radii, which are better estimated than all $K(K-1)/2$ when there are few patients, and it stays at microseconds per pair whatever $K$. It is never smaller than the joint $p$-value, so the test it gives is conservative relative to the joint $p$-value. In practice $p_{\mathrm{D}}$ is about 1.1 times the joint $p$-value with 4 radii and 1.3 to 1.4 times with 8 to 16. Its excess over the joint $p$-value comes from counting crossings rather than exceedances, and from the path between the radii; both shrink as $z_{\max}$ grows, so the bound is tightest at small $p$-values, where discoveries are decided, and loosest at large ones. A radius nearly identical to its neighbour adds an angle near 0 and changes $p_{\mathrm{D}}$ little, so no radius needs to be left out.

When adjacent radii are nearly unrelated, as in the last case above, $p_{\mathrm{D}}$ can exceed Bonferroni's $K \cdot 2\Phi(-z_{\max})$. So the bound suits close radii, and for widely spaced radii the joint $p$-value is better.

##### 6.4.4 The Cauchy combination

As a third option, the radii's own $p$-values $p_1, \ldots, p_K$ are combined without any correlations. Each is turned into a value of the standard Cauchy distribution, and these are averaged:

$$T := \frac{1}{K}\sum_{k=1}^{K} \tan\{(\tfrac{1}{2} - p_k)\pi\}, \qquad p := \frac{1}{2} - \frac{\arctan T}{\pi}.$$

The value $\tan\{(\frac{1}{2} - p_k)\pi\}$ is the point that a standard Cauchy variable exceeds with probability $p_k$, so a small $p_k$ gives a large value: $p_k = 0.01$ gives 31.8. If the radii were independent, $T$ would again be standard Cauchy, and the second formula is the probability that it exceeds the observed $T$. Liu & Xie [77] show that this probability stays approximately correct under any dependence between the radii when it is small, since the Cauchy distribution's tails are so heavy that the dependence between terms matters little there. So the Cauchy combination needs no correlations. The radius with the smallest $p_k$ is reported as the radius with the strongest evidence.

Unlike the maximum, it averages over the radii, so radii with large $p$-values can dilute a strong difference at another. For example, $p$-values of 0.01, 0.9, 0.9 and 0.9 combine to $p = 0.056$, larger than Bonferroni's $4 \times 0.01 = 0.04$.

#### 6.5 The number of patients needed

Spatial omics cohorts are expensive to image, and most hold tens of patients: the 74 analyses of the main text had a median of 35 patients in all. With few patients, a real but small difference in co-localisation between two groups often gives $p \geq \alpha$, so the study reports no change. The number of patients is therefore best worked out before the tissue is collected. For co-localisation this is harder than usual. A patient's information about a pair depends on two things: how many cells of the two types their images hold, which varies many-fold between patients, and how much patients differ from one another. The fitted model estimates these separately, the first through each patient's precision $\Gamma_i$ and the second through the variance between patients $\tau^2$, which together give the patient's weight $u_i$ (Section 4). A model fitted to a pilot cohort therefore gives what the calculation needs. The pilot need not be new. A published cohort of similar tissue, imaged with a similar panel, serves as well, even if its groups differ from those of the planned study, because these quantities describe the tissue and the imaging rather than the comparison. This section derives the number of patients per group, $m^*$, for a two-sided test at level $\alpha$ with 80% power, with the same number of patients in each group.

**A change in co-localisation.** A change is a difference $\Delta = \delta_2 - \delta_1$ between the two groups' mean extra fractions (Section 4.1). Its meaning follows from the room (Section 3.3). On the gain scale, the extra fraction is the share of the $B$ cells that would lack an $A$ neighbour by chance but have one, so $\Delta > 0$ means this share is larger in group 2. On the loss scale, the extra fraction is negative, and its size is the share of the $B$ cells that would have an $A$ neighbour by chance but do not, so $\Delta < 0$ means this share is larger in group 2. $\Delta$ is the smallest change the study should be able to detect, chosen in advance; it is not the difference observed in the pilot, which is too uncertain to plan from.

**What comes from the pilot.** The calculation concerns a new study whose patients have not yet been imaged. Symbols without a hat, such as $\varphi$ and $\bar{u}_c$, are the true values for that study and are unknown; symbols with a hat, such as $\hat{\varphi}$ and $\hat{\bar{u}}_c$, are their estimates from the pilot. $\Delta$ is neither: it is chosen.

**The correction to CR2.** The estimate of $\Delta$ is $D = \hat{\delta}_2 - \hat{\delta}_1$. If the model is accurate, $\hat{\delta}_c$ has variance $1/U_c$, where $U_c = \sum_{i:\, c(i)=c} u_i$ is the total weight of group $c$'s patients (Proposition 10), so the model gives $D$ the variance $1/U_1 + 1/U_2$. The model can be wrong, for example when cells of a type cluster, and patients then vary more than it allows. Write $\varphi$ for the ratio of the true variance of $D$ to the variance the model gives, so $\varphi = 1$ when the model is right. CR2 estimates the variance of $D$ from how far the patients actually spread, $V_1 + V_2$ (Section 5.4), so the pilot estimates $\varphi$ by

$$\hat{\varphi} := \frac{V_1 + V_2}{1/U_1 + 1/U_2},$$

with $V_c$ and $U_c$ from the pilot's patients. With few pilot patients, $\hat{\varphi}$ can be well above or below $\varphi$.

**The mean weights.** Let $\bar{u}_c$ be the mean weight $u_i$ of the patients of group $c$ in the new study, and let

$$\bar{u} := \frac{2}{1/\bar{u}_1 + 1/\bar{u}_2}$$

be the harmonic mean of $\bar{u}_1$ and $\bar{u}_2$. The pilot estimates $\bar{u}_c$ by $\hat{\bar{u}}_c$, the mean weight of its own patients in group $c$.

**Proposition 18** (Number of patients). *Suppose a new study has $m_c$ patients in group $c$, and that*

*(i) the $m_c$ patients of group $c$ have mean weight $\bar{u}_c$;*

*(ii) the variance of $D$ is $\varphi$ times the variance the model gives,* $\mathrm{Var}(D) = \varphi\,(1/U_1 + 1/U_2)$, *with the same $\varphi$ as in the pilot; and*

*(iii) $D$ is normal with mean $\Delta$.*

*Let*

$$\sigma^2 := \varphi\Big(\frac{1}{m_1\bar{u}_1} + \frac{1}{m_2\bar{u}_2}\Big),$$

*which the proof shows is the variance of $D$, and let $\Phi$ be the standard normal distribution function (Section 6.4). Then the power of the two-sided test at level $\alpha$, the chance that the test gives $p < \alpha$ with $D$ of the same sign as $\Delta$, is*

$$\Phi\Big(\frac{|\Delta|}{\sigma} - \Phi^{-1}(1-\alpha/2)\Big).$$

*With the same number of patients in each group, the power is at least 80% exactly when $m_1 = m_2 \geq m^*$, where*

$$m^* := \frac{2\varphi\,\big\{\Phi^{-1}(1-\alpha/2) + \Phi^{-1}(0.8)\big\}^2}{\bar{u}\,\Delta^2}.$$

*Proof.* The two groups have no patients in common, so, with the weights treated as fixed, $\hat{\delta}_1$ and $\hat{\delta}_2$ are independent and

$$\mathrm{Var}(D) = \mathrm{Var}(\hat{\delta}_1) + \mathrm{Var}(\hat{\delta}_2).$$

Under the model, $\mathrm{Var}(\hat{\delta}_c) = 1/U_c$, where $U_c = \sum_{i:\,c(i)=c} u_i$ (Proposition 10). Group $c$ has $m_c$ patients, and by (i) their mean weight is $\bar{u}_c$, so their total weight is $U_c = m_c\bar{u}_c$. The model's variance of $D$ is therefore

$$\frac{1}{m_1\bar{u}_1} + \frac{1}{m_2\bar{u}_2}.$$

By (ii), this is multiplied by $\varphi$, which gives $\sigma^2$, the variance of $D$.

Write $a \coloneqq \Phi^{-1}(1-\alpha/2)$. With no change, $\Delta = 0$ and $D/\sigma$ is standard normal, so the test gives $p < \alpha$ when $|D/\sigma| > a$. By (iii), $D = \Delta + \sigma Z$ with $Z$ standard normal, so $D/\sigma = \Delta/\sigma + Z$. For $\Delta > 0$, the test gives $p < \alpha$ with $D$ of the same sign when $\Delta/\sigma + Z > a$, that is, $Z > a - \Delta/\sigma$, which has chance $1 - \Phi(a - \Delta/\sigma) = \Phi(\Delta/\sigma - a)$ by the symmetry of the normal. For $\Delta < 0$, it does so when $\Delta/\sigma + Z < -a$, that is, $Z < -a - \Delta/\sigma$, which has chance $\Phi(-\Delta/\sigma - a)$. Both are $\Phi(|\Delta|/\sigma - a)$, the power stated. The test can also give $p < \alpha$ with $D$ of the opposite sign to $\Delta$, but it would then report the change the wrong way round, so this does not count as detecting it. Its chance is $\Phi(-|\Delta|/\sigma - a)$. At 80% power, $|\Delta|/\sigma = a + \Phi^{-1}(0.8)$, so this chance of detecting a change in the opposite direction is $\Phi(-2a - \Phi^{-1}(0.8))$, which is below $10^{-6}$ for any $\alpha \le 0.05$. Therefore, counting it would add less than $10^{-6}$ to the power, which makes no practical difference to $m^*$, so leaving it out loses nothing.

Now let $m_1 = m_2 = m$. Because $\bar{u}$ is the harmonic mean of $\bar{u}_1$ and $\bar{u}_2$, $1/\bar{u}_1 + 1/\bar{u}_2 = 2/\bar{u}$, so

$$\sigma^2 = \frac{2\varphi}{m\bar{u}}.$$

This falls as $m$ grows, so the power rises with $m$. It is 80% when $|\Delta|/\sigma - a = \Phi^{-1}(0.8)$, that is, when

$$\Delta^2 = \left\{a + \Phi^{-1}(0.8)\right\}^2 \frac{2\varphi}{m\bar{u}}.$$

Solving for $m$ gives $m = m^*$, so the power is at least 80% exactly when $m \ge m^*$. □

In practice, $\bar{u}_c$ and $\varphi$ are unknown, so $m^*$ is computed with the pilot's $\hat{\bar{u}}_c$ and $\hat{\varphi}$ in their place, and rounded up.

**Why the harmonic mean.** $\bar{u}$ is the harmonic mean of $\bar{u}_1$ and $\bar{u}_2$, not the mean weight of all patients, because $\mathrm{Var}(D)$ adds the variances of the two group means. The two differ when one group has less information per patient, for example fewer $B$ cells per image. That group's variance is the larger of the two terms, and the harmonic mean follows it.

*Remark* 4 (More cells or more patients?). A patient's weight is $u_i = 1/(1/\Gamma_i + \tau^2)$ (Section 4.2), where $\Gamma_i$ grows with the cells in the patient's images. However many cells are imaged, $u_i$ stays below $1/\tau^2$. Once $1/\Gamma_i$ is small next to $\tau^2$, more cells per patient add little, and only more patients reduce $\mathrm{Var}(D)$.

**What the number assumes.** $m^*$ is a planning figure, not a guarantee: it is only as good as the pilot's estimates. Assumption (i) holds if new patients are imaged as the pilot's were, with similar numbers of images and cells, which the design of the study controls. Assumption (ii) holds if new patients vary about the model as the pilot's do, which the design does not control. Assumption (iii) replaces the $t$ distribution on Satterthwaite degrees of freedom that the test uses (Section 5.6) by the normal, so when $m^*$ is small it is slightly too small. The pair must also keep the same scale, gain or loss, so that $\Delta$ has the same meaning.

The weakest links are $\hat{\varphi}$ and $\hat{\tau}^2$, which sets the weights $u_i$. Both measure how much patients vary, and a pilot with few patients estimates them poorly. An estimate that is too small, of either, makes $m^*$ too small: a smaller $\hat{\tau}^2$ gives larger weights and so a larger $\hat{\bar{u}}$. Planning with a larger $\hat{\varphi}$, or recomputing $m^*$ once the first patients of the study are in, as in an internal pilot [78], guards against this.

**The smallest change a cohort can detect.** By the same argument with $\mathrm{Var}(D) = V_1 + V_2$, a cohort that has already been collected can detect, with 80% power at level $\alpha$, a change of

$$\Delta_{\min} := \left\{\Phi^{-1}(1-\alpha/2) + \Phi^{-1}(0.8)\right\}\sqrt{V_1 + V_2},$$

which uses its own CR2 variance and needs no assumption about new patients.

### 7 Algorithm and cost

This section puts the steps of Sections 2–5 together as one algorithm (Algorithm 1) and gives the time each step takes. The algorithm is written for two groups without covariates. Then every estimate and variance has a closed form (Propositions 7, 10 and 12), so no general GEE solver and no matrix inversion is needed, only sums over images and patients. The only numerical steps are finding $\hat{\omega}^2$ and $\hat{\tau}^2$, and each is the single crossing of a decreasing function (Section 4.3).

With several radii, lines 2–29 are run at each radius, the radii are combined into one $p$-value per pair (Section 6.4), and line 30 adjusts these.

**Notation for the cost.** Write $G$ for the number of cell types, so that there are $G^2$ pairs, including the $G$ self-pairs. An image has $n$ cells, and $\kappa$ is the average number of neighbours of a cell: $\kappa \approx \rho\pi r^2$ for cells at density $\rho$ with the radius graph, and $\kappa = k$ with the $k$-nearest-neighbour graph. $J = \sum_i J_i$ is the total number of images, and $N$ the total number of cells over all images.

Times are given in $O$ notation. A time of $O(n)$ grows at most in proportion to $n$, up to a constant that does not depend on the data: twice as many cells, at most about twice the time. Constants are left out, so $O(\cdot)$ says how the time grows with the data, not how many seconds it takes.

**Two kinds of work.** The steps of Algorithm 1 are of two kinds, and they are costed separately.

1. *Work on the cells* (lines 2–5 and 7–10). These steps read every cell. Lines 2–5 find the neighbours and count them, once per image. Lines 7–10 sit inside the loop over pairs: for each pair, they pass over the cells of each image once more and reduce the image to four numbers, $n^B$, $O$, $E$ and $V$.

**Algorithm 1** The method at one radius $r$.

**Require:** for every cell, its location, cell type, image and patient; the group of every patient; the radius $r$ (or $k$); the effect (extra fraction or extra count).

Remove the images with no group. $\triangleright$ Section 2

**for** each image **do**

Find the neighbours of every cell.

For every cell and every cell type $A$, count the $A$ cells among its neighbours.

**end for**

**for** each pair $A \to B$ **do**

**for** each image **do**

Set the $x_\ell$ of the candidates, and compute $O$, $E$ and $V$. $\triangleright$ Sections 3.1–3.8

If $n^B = 0$ or $V = 0$, do not use the image. $\triangleright$ Section 3.5

**end for**

**if** the effect is the extra fraction **then**

Choose the scale from the sign of $\sum(O - E)$ over all images. $\triangleright$ Section 3.3

Give each image its room $R$; do not use images with $R < 1$. $\triangleright$ Section 3.5

$Y \leftarrow (O - E)/R$ and $s^2 \leftarrow V/R^2$ for each image.

**else**

$Y \leftarrow (O - E)/n^B$ and $s^2 \leftarrow V/(n^B)^2$ for each image. $\triangleright$ Section 3.4

**end if**

Leave out the patients none of whose images is used. $\triangleright$ Section 4

Find $\hat{\omega}^2$ from $Q(t) = d$, or set it to 0 if $Q(0) \le d$. $\triangleright$ Section 4.3

Compute the image weights $\gamma_{ij}$ and the patient means $\bar{Y}_i$. $\triangleright$ Section 4.2

Find $\hat{\tau}^2$ from $X(t) = m - 2$, or set it to 0 if $X(0) \le m - 2$. $\triangleright$ Section 4.3

Compute $u_i$, $\hat{\delta}_1$, $\hat{\delta}_2$ and $D = \hat{\delta}_2 - \hat{\delta}_1$. $\triangleright$ Proposition 7

Compute $\hat{e}_i$, $h_i$, $V_1$, $V_2$, $m_1^{\text{eff}}$, $m_2^{\text{eff}}$ and $\nu$. $\triangleright$ Sections 5.3–5.6

**if** one of the five cases of Section 5.8 holds **then**

Report the pair as not tested, with the reason.

**else**

$t \leftarrow D/\sqrt{V_1 + V_2}$, and $p \leftarrow 2\Pr(T_\nu > |t|)$. $\triangleright$ Section 5.7

**end if**

**end for**

Adjust the $p$-values of the tested pairs. $\triangleright$ Section 5.7

2. *Work on the image summaries* (lines 11–28). These steps use only the four numbers of each image and never look at the cells again. So their time depends on the numbers of images and patients, not of cells.

Lines 7–10 are therefore costed with the cells, although they sit in the loop over pairs.

**Work on the cells.** For an image of $n$ cells:

- *Neighbours (line 3).* With a spatial index such as a $k$-d tree, finding the neighbours of every cell takes $O(n \log n + n\kappa)$ time.
- *Counts (line 4).* One pass over the $n\kappa$ links gives every cell's number of neighbours of each type, for all $G$ types at once, in $O(n\kappa)$ time.
- *A pair $A \to B$ (line 8).* With the counts in hand, $x_\ell$, $\bar{x}$, $\sigma_x^2$ and $O$ take one pass over the $n$ cells, in $O(n)$ time, so $O(G^2 n)$ for all pairs.
- *A self-pair (line 8, $A = B$).* The extra count needs only the numbers of links at each cell (with the radius graph, $\sum_\ell d_\ell$ and $\sum_\ell d_\ell^2$), so $O(n\kappa)$ time. The extra fraction also needs the pairs of cells that share a neighbour. Found exactly, they take $O(n\kappa^2)$ time. By default they are estimated from a sample of cells, and the time falls to $O(n\kappa)$ (Remark 2). With $G$ self-pairs, this is $O(Gn\kappa)$.

Adding these, one image takes $O(n \log n + Gn\kappa + G^2 n)$ time, and all images together $O(N \log n_{\max} + GN\kappa + G^2 N)$, where $n_{\max}$ is the number of cells of the largest image.

Since the null moments are exact (Corollary 1.1), this is done once. A permutation test would instead repeat line 8 for every permutation of every image, so with 1,000 permutations the work on the cells would be about 1,000 times larger.

**Work on the image summaries.** For one pair, with $J$ images:

- *The effects (lines 11–18)* take $O(J)$ time.
- *The variance components (lines 19 and 21).* Each value of $Q(t)$ or $X(t)$ takes $O(J)$ time. Both functions decrease, so a bracketing search (Brent's method) finds the crossing to a fixed precision in a number of steps $I$ that does not grow with the numbers of images or patients. Each component then takes $O(IJ)$ time.
- *The estimate and the test (lines 20 and 22–28)* take $O(J)$ time.

So one pair takes $O(IJ)$ time, and all pairs $O(G^2 IJ)$. The adjustment (line 30) sorts the $p$-values, which takes $O(P \log P)$ time for $P$ tested pairs.

**An example.** Take 100 patients with one image each, $n = 10{,}000$ cells per image, $G = 20$ cell types (400 pairs) and $\kappa = 20$ neighbours per cell. The work on the cells is about $Gn\kappa + G^2 n = 4 \times 10^6 + 4 \times 10^6$ per image, or $8 \times 10^8$ over the 100 images. The work on the summaries, with $I = 50$ steps for illustration, is about $G^2 IJ = 400 \times 50 \times 100 = 2 \times 10^6$ for each variance component. So the cells account for almost all of the time, and the comparison of groups is cheap however many pairs are tested.

**Memory.** An image's neighbours take $O(n\kappa)$ memory, and the images can be processed one at a time, so the largest image sets the memory needed. After line 10 each image keeps only a few numbers for each pair ($n^B$, $O$, $E$, $V$ and, for the extra fraction, $q$ or the room), $O(G^2 J)$ memory in all. In the example above, this is $400 \times 100$ sets of a few numbers.

**Summary.** The time is dominated by the work on the cells and grows in proportion to the total number of cells, apart from the $\log n$ of the neighbour search. The comparison of groups grows only with the numbers of images and patients, and the method needs no permutations.

### 8 Notation

The table lists the symbols used in more than one place, in the order they appear, with the section where each is introduced. Symbols used only inside one proof are defined there and are not listed. Bold symbols are vectors or matrices; all others are numbers, sets or functions. Within one image, the subscript $(i, j)$ is dropped.

| Symbol | Meaning | Section |
|---|---|---|
| *Data* | | |
| $m$, $i$ | number of patients; patient index | 2 |
| $c(i)$, $m_c$ | group of patient $i$ (1 or 2); number of patients in group $c$ | 2 |
| $J_i$, $j$ | number of images of patient $i$; image index | 2 |
| $n_{ij}$ | number of cells in image $(i, j)$ | 2 |
| $A \to B$ | pair: $A$ the anchor type, $B$ the tested type | 2 |
| $n_{ij}^A$, $n_{ij}^B$ | numbers of $A$ and $B$ cells in image $(i, j)$ | 2 |
| $r$, $k$ | radius of the radius graph; number of neighbours in the $k$-nearest-neighbour graph | 2 |
| *The effect in one image* | | |
| $M$, $\ell$ | number of candidates (cells that are not $A$), $M = n - n^A$; candidate index | 3.1 |
| $p$ | share of candidates that are $B$, $p = n^B/M$ | 3.1 |
| $x_\ell$ | number of candidate $\ell$: 1 or 0 for the extra fraction, its number of $A$ neighbours for the extra count | 3.1 |
| $O$ | observed total, $\sum_{\ell \in B} x_\ell$ | 3.1 |
| $\bar{x}$, $\sigma_x^2$ | mean and variance of the $x_\ell$ over the candidates | 3.2 |
| $E$, $V$ | mean and variance of $O$ under random labelling | 3.2 |
| $q$ | share of candidates with an $A$ neighbour (the extra fraction's $\bar{x}$) | 3.3 |
| $R$ | room: $n^B(1-q)$ on the gain scale, $n^B q$ on the loss scale | 3.3 |

| Symbol | Meaning | Section |
|---|---|---|
| $Y$ | effect of an image: $(O-E)/R$ (extra fraction) or $(O-E)/n^B$ (extra count) | 3.3, 3.4 |
| $s^2$ | variance of $Y$ under random labelling: $V/R^2$ or $V/(n^B)^2$ | 3.3, 3.4 |
| $\lambda$ | mean number of $A$ neighbours of a candidate (the extra count's $\bar{x}$) | 3.4 |
| $Y_F$, $Y_C$; $O_F$, $O_C$ | extra fraction and extra count; their observed totals | 3.6 |
| $c_\ell$ | number of $A$ cells within $r$ of candidate $\ell$ | 3.6 |
| $\bar{d}$ | depth: mean number of $A$ neighbours of a candidate that has one, $\lambda/q$ | 3.6 |
| $d_B$ | depth of the $B$ cells, $O_C/O_F$ | 3.6 |
| *Models of co-localisation (Section 3.7)* | | |
| $f$ | share of $B$ cells drawn to $A$ (the behaviour) | 3.7 |
| $\varrho$ | largest distance from a recruited cell to its hub | 3.7.2 |
| $q_0$, $\lambda_0$ | $q$ and $\lambda$ among the candidates that are not moved | 3.7.2 |
| $\rho_A$ | crowding of $A$ cells around a hub | 3.7.2 |
| *Self-pairs (Section 3.8)* | | |
| $I_\ell$ | 1 if cell $\ell$ is an $A$ cell, 0 otherwise | 3.8 |
| $\pi_k$ | probability that $k$ given cells are all $A$ cells | 3.8 |
| $\mathcal{N}(\ell)$ | neighbours of cell $\ell$, not including $\ell$ | 3.8 |
| $d_\ell$, $w_\ell$, $t_\ell$ | numbers of links that start at, end at, and touch cell $\ell$ | 3.8 |
| $\mathcal{L}$, $m_{\mathrm{L}}$, $b$ | set of links; their number; number of links whose reverse is a link | 3.8 |
| $h(s)$, $h_2(s)$ | chance that none of $s$ given cells is an $A$ cell, given one or two $A$ cells | 3.8 |
| $T_{\ell\ell'}$ | chance that cells $\ell$ and $\ell'$ both have an $A$ neighbour, given both are $A$ | 3.8 |
| $L$, $\tilde{p}$ | self-pair extra count: number of links that end at an $A$ cell; $\tilde{p} = (n^A - 1)/(n-1)$ | 3.8 |
| $S$ | $\sum_\ell t_\ell(t_\ell - 1)$ | 3.8 |
| $\kappa$ | average number of neighbours of a cell | 3.8.2 |
| *Combining images within patients* | | |
| $Y_{ij}$, $s_{ij}^2$ | effect of image $(i,j)$ and its random-labelling variance | 4 |
| $\delta_c$ | mean effect of group $c$ | 4.1 |
| $b_i$, $\tau^2$ | departure of patient $i$; variance between patients | 4.1 |
| $\eta_{ij}$, $\omega^2$ | departure of image $(i,j)$ from its patient; variance between images of a patient | 4.1 |
| $\varepsilon_{ij}$ | random-labelling noise of image $(i,j)$, with variance $s_{ij}^2$ | 4.1 |

| Symbol | Meaning | Section |
|---|---|---|
| $\mathbf{Y}_i$ | vector of patient $i$'s effects ($J_i \times 1$) | 4.1 |
| $\mathbf{V}_i$ | working covariance of $\mathbf{Y}_i$ ($J_i \times J_i$) | 4.1 |
| $\mathbf{1}$ | vector of ones | 4.1 |
| $\gamma_{ij}$, $\Gamma_i$ | image weight $1/(s_{ij}^2 + \omega^2)$; patient $i$'s total image weight | 4.2 |
| $\bar{Y}_i$ | patient $i$'s weighted mean effect | 4.2 |
| $u_i$ | patient weight, $1/(1/\Gamma_i + \tau^2)$ | 4.2 |
| $\hat{\delta}_c$ | estimate of $\delta_c$ | 4.2 |
| $Q(t)$, $d$ | spread of images within patients at trial value $t$; its expected value $\sum_i (J_i - 1)$ | 4.3 |
| $X(t)$ | spread of patients within groups at trial value $t$ | 4.3 |
| $\hat{\omega}^2$, $\hat{\tau}^2$ | estimates of $\omega^2$ and $\tau^2$ | 4.3 |
| *Comparing groups* | | |
| $D$ | difference $\hat{\delta}_2 - \hat{\delta}_1$ | 5.1 |
| $f_i$ | error of patient $i$'s mean, $\bar{Y}_i - \delta_{c(i)}$ | 5.1 |
| $U_c$ | total patient weight of group $c$, $\sum_{i:\, c(i)=c} u_i$ | 5.1 |
| $\hat{e}_i$ | residual of patient $i$, $\bar{Y}_i - \hat{\delta}_{c(i)}$ | 5.2 |
| $V_c^{\text{CR0}}$ | CR0 variance of $\hat{\delta}_c$ | 5.2 |
| $\hat{\mathbf{r}}_i$ | residual vector of patient $i$, $\mathbf{Y}_i - \hat{\delta}_c \mathbf{1}$ | 5.2 |
| $h_i$ | leverage of patient $i$, $u_i/U_c$ | 5.3 |
| $m_c^{\text{eff}}$ | effective number of patients of group $c$, $1/\sum_i h_i^2$ | 5.3 |
| $V_c$ | CR2 variance of $\hat{\delta}_c$ | 5.4 |
| $\mathbf{V}_i^{-1/2}$ | symmetric inverse square root of $\mathbf{V}_i$ | 5.5 |
| $\tilde{\mathbf{x}}_i$, $\tilde{\mathbf{r}}_i$ | whitened design and residuals of patient $i$ | 5.5 |
| $\mathbf{A}_i$ | CR2 adjustment matrix of patient $i$ | 5.5 |
| $V_c^{\text{CR2}}$ | CR2 variance of $\hat{\delta}_c$ in matrix form; equals $V_c$ | 5.5 |
| $\mathbf{H}_{ii}$ | patient $i$'s block of the hat matrix | 5.5 |
| $\nu$, $T_c$ | Satterthwaite degrees of freedom; group $c$'s part of its denominator | 5.6 |
| $t$ | test statistic, $D/\sqrt{V_1 + V_2}$ | 5.7 |
| $T_\nu$ | a variable with a $t$ distribution on $\nu$ degrees of freedom | 5.7 |
| *Extensions* | | |
| $\mathbf{x}_{ij}$, $\boldsymbol{\beta}$ | covariates of image $(i, j)$; their slopes (within-patient for image-level covariates) | 6.1.1 |
| $\bar{\mathbf{x}}_i^{\text{I}}$, $\boldsymbol{\lambda}$ | patient averages of the image-level covariates; their added slopes | 6.1.2 |
| $\mathbf{x}_{ij}^{\text{I}}$, $\mathbf{x}_i^{\text{P}}$; $\boldsymbol{\beta}^{\text{I}}$, $\boldsymbol{\beta}^{\text{P}}$ | image-level and patient-level covariates; their slopes | 6.1.2 |

| Symbol | Meaning | Section |
|---|---|---|
| $\mathbf{z}_{ij}$, $\boldsymbol{\theta}$ | row of the design for image $(i, j)$; its coefficients | 6.1.3 |
| $\mathbf{Z}_i$, $\mathbf{X}_i$ | patient $i$'s rows of the design; the same rows whitened | 6.1.4 |
| $\mathbf{B}$, $\mathbf{c}$ | bread $(\sum_i \mathbf{X}_i^\top \mathbf{X}_i)^{-1}$; contrast picking out the tested entry of $\boldsymbol{\theta}$ | 6.1.4 |
| $\boldsymbol{\Omega}$, $\Omega_{ik}$ | matrix of the whole fit whose traces give $\nu$; its entry for patients $i$ and $k$ | 6.1.4 |
| $C$, $\mathbf{L}$, $g$ | number of groups; contrasts of the overall test; their number, $C - 1$ | 6.2 |
| $Q_{\mathrm{F}}$ | Wald statistic of the overall test | 6.2 |
| $\nu_{\mathrm{F}}$ | degrees of freedom of the overall test | 6.2 |
| $\mathbf{W}$, $\boldsymbol{\Sigma}_{ik}$ | scaling that gives $\mathbf{V}^{\mathrm{CR2}}$ of the overall test mean $\mathbf{I}$; covariance of patients $i$'s and $k$'s scaled terms | 6.2 |
| $\alpha$ | level: of the false discovery rate, or of a test | 6.2 |
| $\mathbf{w}_i$, $\hat{H}_i$, $d_i$ | covariates of the Cox model; expected number of events; event indicator | 6.3 |
| $h_i(t)$, $h_0(t)$, $\boldsymbol{\zeta}$ | hazard of patient $i$ at time $t$; baseline hazard; slopes of the Cox model | 6.3 |
| $M_i$, $\theta_0$, $\theta_1$ | martingale residual; intercept and slope on $M_i$ | 6.3 |
| $t_k$, $\nu_k$, $z_k$ | statistic, degrees of freedom and normal score at radius $r_k$ | 6.4 |
| $\Phi$ | standard normal distribution function | 6.4 |
| $z_{\max}$ | largest $z_k$ over the radii | 6.4 |
| $\psi_{ik}$, $\hat{\rho}_{kl}$, $\mathbf{R}$ | patient $i$'s signed term of the CR2 variance of $D$ at radius $r_k$; correlation between radii; their matrix | 6.4 |
| $\Lambda$, $p_{\mathrm{D}}$ | length of the path through the radii; Davies' bound | 6.4 |
| $\Delta$ | difference between the group means that a study should detect | 6.5 |
| $\varphi$, $\hat{\varphi}$ | ratio of the true variance of $D$ to the model's; its estimate from a cohort | 6.5 |
| $\bar{u}_c$, $\bar{u}$ | mean patient weight of group $c$; harmonic mean of $\bar{u}_1$ and $\bar{u}_2$ | 6.5 |
| $m^*$ | patients needed per group for 80% power | 6.5 |
| $\Delta_{\min}$ | smallest difference a cohort can detect with 80% power | 6.5 |
| *Algorithm and cost* | | |
| $G$ | number of cell types | 7 |
| $J$ | total number of images, $\sum_i J_i$ | 7 |
